# Multiscale Parcellation of Dynamic Causal Models of the Brain

**DOI:** 10.1101/2025.06.14.659698

**Authors:** Tahereh S. Zarghami

## Abstract

The hierarchical organization of the brain’s distributed network has received growing interest from the neuroscientific community, largely because of its potential to enhance our understanding of human cognition and behavior, in health and disease. This interest is motivated by the hypothesis that near-critical brain dynamics enable multiscale integration and segregation of neural dynamics. While most multiscale connectivity analyses focus on structural and functional networks, characterizing the effective connectome across multiple scales has been somewhat overlooked—primarily for computational reasons. The difficulty of estimating large cyclic causal models, together with the scarcity of theoretical frameworks for systematically moving between scales, has hindered progress in this direction. This technical note introduces a top-down multiscale parcellation scheme for dynamic causal models, with application to neuroimaging data. The method is based on Bayesian model comparison, as a generalization of the well-known Δ*BIC* method. To facilitate computation, recent developments in linear dynamic causal modeling (DCM) and Bayesian model reduction (BMR) are deployed. Specifically, a naïve version of BMR is introduced, enabling the parcellation scheme to scale to hundreds or thousands of regions. Notably, the derivations reveal an analytical relationship between reduced model evidence and minimum cut problem in graph theory. This duality puts the tools of graph theory at the service of model evidence optimization and significance testing. The proposed method was applied to simulated and empirical causal models to establish face and construct validity. Consequently, the large empirical causal network, inferred from a neuroimaging dataset, exhibited log–log scaling trends, suggestive of scale invariance in multiple dynamical measures. Future generalizations of this technique and its potential applications in systems and clinical neuroscience are discussed.

## Introduction

Brain connectomic studies have been progressing towards multiscale analyses in recent years (Axer & Amunts, 2022; Betzel & Bassett, 2017; D’Angelo & Jirsa, 2022; Lu et al., 2022; Schirner et al., 2018; Shine et al., 2021; Van Den Heuvel et al., 2019). A key motivation for this advancement is the hypothesis that the brain operates near a *critical*^1^ regime (Beggs & Plenz, 2003; Chialvo, 2010) that optimally enables and coordinates the brain’s multiscale phenomena, from microscopic neuronal activity to macroscopic cognition (Boonstra et al., 2013; Cocchi et al., 2017; Haimovici et al., 2013; Hengen & Shew, 2025; Hesse & Gross, 2014; Linkenkaer-Hansen et al., 2001; Rubinov et al., 2011; Shew & Plenz, 2013; Tognoli & Kelso, 2014; Van de Ville et al., 2010; Zimmern, 2020, 2020). As such, understanding the full richness of the brain’s multiscale structure and function—across time, space and topology— has motivated a large body of empirical work (Hansen et al., 2023; Maksimenko et al., 2018; Morante, Frølich, Shahzaib, et al., 2024; Munn et al., 2024) and multiple methodological developments (Doucet et al., 2011; Farahibozorg et al., 2024; Morante, Frølich, & Rehman, 2024; Presigny & Fallani, 2022). Multiscale analyses are specifically attractive, because they hold the potential to illuminate the complex relationship between the brain’s hierarchical organization, its self-organized criticality, and human behavior and cognition in health, disease, injury, development, and aging (Bastiani & Roebroeck, 2015; Betzel & Bassett, 2017; Bonifazi et al., 2018; He et al., 2025; Latifi & Carmichael, 2024; Lu et al., 2022; Pines et al., 2022; Sala et al., 2017; Zhou et al., 2023; Zou et al., 2024).

While hierarchical decomposition of structural and functional networks are ubiquitous in the neuroimaging literature (Ashourvan et al., 2019; Ferrarini et al., 2009; Iraji et al., 2023; Meng et al., 2022; Meunier et al., 2010; Mirzaeian et al., 2025; Park et al., 2022; Rousseau et al., 2025), parcellating effective (i.e. causal) networks of the brain remains largely unexplored, with a few exceptions (D’Acunto et al., 2024; Friston et al., 2021; Wu et al., 2011; Wang & Shanechi, 2019). The challenges are mostly computational, as well as theoretical (Bhattacharya et al., 2024). While learning the structure and parameters of large (cyclic) Bayesian networks is computationally demanding, testing hypotheses about their potential modular organization calls for further theoretical work. But the effort is warranted, because neither structural nor functional network analyses can offer the mechanistic insight that a model-based causal framework can. For instance, (Friston et al., 2021) have recently shown that recursive coarse-graining and demarcation of neuronal states based on their effective connectivity (plus summarizing their dynamics at each scale based on the renormalization group theory) can explain the emergence of large-scale networks, self-organized criticality and dynamical instability in the brain. These phenomena can be elegantly studied within a causal modeling framework.

In this work, we present a top-down^2^ parcellation method for effective connectivity graphs. As we will see, the advantage of the top-down treatment is the systematic identification of subordinate components, as opposed to the more arbitrary designation of internal states in the bottom-up approach (Friston et al., 2021). This advantage comes at the cost of further computational challenges in the top-down scheme. Fortunately, recent developments in the estimation of large linearized DCMs (Frässle et al., 2017, 2021; Friston et al., 2021) and sparsification of densely connected Bayesian networks (Friston et al., 2019; Marković et al., 2024; Prando et al., 2020; Wright et al., 2024) have facilitated the estimation of large sparse DCMs—including hundreds or thousands of regions. Hence, the present paper tackles the remaining issue of revealing the hierarchical modular structure of a (large) causal network, starting at the coarsest scale.

In short, the present study offers a top-down recursive parcellation scheme for a (potentially large) effective connectivity graph, such that at each scale the neuronal states are (bi)partitioned into statistically significant subordinate parcels. To ease the computations, the proposed scheme relies on a naive formulation of Bayesian model reduction (BMR) that eliminates costly matrix multiplications, and reveals an analytical relationship between the reduced model evidence and the minimum cut problem in graph theory.

The rest of this paper is organized as follows: In the Theory section, we go through the rationale of the proposed approach and its relation to the better-known Δ*BIC* method. Therein, we will see how causal model comparison intersects with concepts from graph theory. Assessing the significance of the identified partitions is covered in the same section. Thereafter, illustrative examples of bi-partitioning simulated (modular and nonmodular) causal networks are presented to show the functionality of the proposed approach. Next, the same method is applied to the causal graph estimated from an empirical neuroimaging dataset—to reveal the hierarchical organization in a dynamic causal model of the brain. We will show the scaling behavior of multiple dynamical measures for this empirical causal network. Furthermore, we will study whether the partitions identified from the causal graph can be interpreted as statistically independent subsystems conditioned upon their boundary (i.e. mediating) states. This question is of interest because recent (simulation) work (Aguilera et al., 2022) has suggested that recurrent connections in cyclic causal models can induce statistical dependencies that cross causal boundaries. We will provide a hypothesis testing framework that can inspect this phenomenon for any given dataset and causal model. Finally, potential applications of the proposed parcellation scheme in systems and clinical neuroscience are discussed, and some future lines of research are outlined.

### Theory

Dividing a group of data points into several cohesive groups is a recurrent problem in many domains, and has been addressed using a multitude of unsupervised methods. These methods are described under different umbrella terms^3^: clustering, partitioning, segmentation, community or module detection, change point detection, blind source separation, and data decomposition, among others. The solution is usually formulated to either minimize within-group^4^ *dispersion* or maximize within-group *cohesion*. The measure of *dispersion* may be, for example, *variance* in K-means clustering (Lloyd, 1982) or *entropy* in infomax-based independent component analysis (ICA) (Bell & Sejnowski, 1995). Likewise, *cohesion* may be defined in terms of *connectedness* in density-based clustering (Ester et al., 1996) and modularity maximization (Newman, 2006), or in terms of data *likelihood* under distributional assumptions in Gaussian mixture models (Dempster et al., 1977) and maximum likelihood-based ICA (Ans et al., 1985; Hyvärinen, 2013), among others.

Specifically, maximum likelihood, when penalized for model complexity, has commonly been used as a model selection criterion to divide a dataset (usually timeseries) into two or more coherent segments. The goal is to find the breakpoint(s) such that fitting a separate model to each segment would yield a more plausible explanation for the data, than fitting a single model to the entire data^5^. The plausibility of each model is determined by a criterion of the form: −2 log likelihood + a penalty term for model complexity as a function of the number of model parameters and sample size. Depending on the form of the penalty, the criterion becomes: Akaike information criterion (AIC), Bayesian information criterion (BIC), or minimum description length (MDL) (Gao et al., 2024). For example, in a bisegmentation problem, the optimal breakpoint is the one that maximally reduces *BIC*_1_ + *BIC*_2_ compared to *BIC*_0_; where *BIC*_1_ and *BIC*_2_ result from fitting separate models to the two segments (before and after the breakpoint) and *BIC*_0_ is computed for the single model fitted to the entire data. This approach is known as the Δ*BIC* method, since the objective is to minimize the following cost function (Chen & Gupta, 1997; Gao et al., 2024):

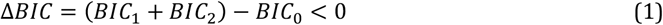

From a more statistically principled perspective, the same problem has been formulated within a hypothesis testing framework, where the null hypothesis *H*_0_ corresponds to a model without any breakpoints and the alternative hypothesis *H*_*a*_ posits the existence of a breakpoint (Gao et al., 2024). Hypothesis testing based on Δ*BIC* is a hybrid approach (Chen & Gupta, 1997), in which a significance level *α* can be associated with the model selection result. This is realized by introducing a critical value *C*_*α*_ ≥ 0 such that the null hypothesis is rejected only when:

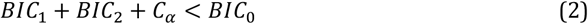

Intuitively, the role of *C*_*α*_ is to ensure that the difference between *BIC*_1_ + *BIC*_2_ and *BIC*_0_ is statistically convincing and not merely driven by noise. Approximate *C*_*α*_ values have been computed by (Chen & Gupta, 1997) for different significance levels *α*, based on the asymptotic null distribution of Δ*BIC*. The Δ*BIC* method—with or without hypothesis testing—has been used in a variety of contexts to segment different kinds of temporal (Chung-Hsien Wu et al., 2006; Costa et al., 2016; Theodorou et al., 2014; Tritschler & Gopinath, 1999; Xue et al., 2010) and spatial data (Vatsavai et al., 2011; Zhao et al., 2008); see (Gao et al., 2024) for a recent review and numerous references.

Here we adopt a conceptually similar and statistically grounded model comparison approach to justify the (multiscale) parcellation of a dynamic causal model fitted to regional timeseries. The goal is to partition the regions associated with the timeseries. As we will shortly see, once we move beyond maximum likelihood to a fully Bayesian treatment, the heuristic formulations of the Δ*BIC* method emerge naturally and turn out to be simplified forms of established constructs in Bayesian model comparison. Note that the same methods can be used to divide the data into more than two partitions (Chen & Gupta, 1997; Gao et al., 2024). Hence, we begin with the bi-partitioning problem and later discuss its generalization to multi-partitions.

Problem statement:

Given a dynamic causal model fitted to a distributed network’s time series, how can the inferred causal information be used to derive a plausible bipartition of the system, and how can the plausibility of such a partition be quantified?

As should be clear from the preceding introduction, this is fundamentally a model comparison problem; and the solution offered here is inspired by the Δ*BIC* method. In short, we compare the *null hypothesis* (*H*_0_) of fitting a single DCM to all regional timeseries with the *alternative hypothesis* (*H*_*a*_) of fitting two DCMs separately to the partitioned regions. Notably, each candidate bipartition of the network constitutes a distinct alternative hypothesis, associated with its own model evidence. The winning model, among these alternatives, reveals the most plausible way to divide the overall causal network into two interacting causal modules. This constitutes the central theme of the present work. In what follows, we elaborate on the implementation of this model comparison and propose solutions to the associated computational challenges, organized around six key points. Key statistical and Bayesian modeling terms referenced throughout are defined both intuitively and formally in Glossary Table 1.

**Table 1:** Glossary of statistical and Bayesian modeling and inference terms.

| Term | Definition | Formula |
| --- | --- | --- |
| **$\Delta BIC$** | $\Delta BIC$ is a method for comparing models by examining the difference in their Bayesian information criterion (BIC) values.<br>$\Delta BIC$ can be interpreted as a large-sample approximation to (negative twice) the log Bayes factor, under regularity conditions.<br>See **BIC** and **Bayes factor**. | $\Delta BIC$ $= BIC(M1) - BIC(M2)$ $\approx -2 \log BF_{12}$<br>$M1$: Model 1<br>$M2$: Model 2 |
| **$\Delta F$** | $\Delta F$ reflects the relative explanatory power of two models.<br>Technically, $\Delta F$ is the difference in the variational free energies of two models, which (when optimized) approximate the respective log model evidences. As such, $\Delta F$ approximates the log Bayes factor (BF).<br>See **Free energy** and **Bayes factor**. | $\Delta F = F(M1) - F(M2)$ $\approx \log \frac{P(y\vert M1)}{P(y\vert M2)} = \log BF_{12}$ |
| **Bayes factor (BF)** | Bayes factor quantifies how strongly the data supports one model over another.<br>Technically, it is the ratio of model evidences (also known as marginal likelihoods).<br>See **Model evidence**. | $BF_{12} = \frac{P(y\vert M1)}{P(y\vert M2)}$ |
| **Bayesian information criterion (BIC)** | BIC is a criterion for model selection that balances model fit and complexity to promote parsimony. Models with lower BIC are generally preferred.<br>Technically, BIC is an asymptotic approximation to (negative twice) the log model evidence, which penalizes model complexity via the number of free parameters. | $BIC = -2 \log \hat{L} + k \log n$<br>$\hat{L} = P(y\vert \hat{\theta}, M)$: maximized likelihood<br>$k$: number of parameters<br>$n$: sample size |
| **Bayesian model reduction (BMR)** | BMR evaluates simpler nested models relative to a full model efficiently.<br>Technically, it computes the model evidence and posteriors of reduced (i.e. nested) models from a full model's posterior and evidence, without re-estimating parameters, using analytic expressions. | $\frac{P(y\vert M^R)}{P(y\vert M)}$ $= \int P(\theta\vert y, M) \frac{P(\theta\vert M^R)}{P(\theta\vert M)} d\theta$<br>(see Appendix B) |
| **Dynamic causal modeling (DCM)** | DCM explains observed time-series data by modeling hidden causal interactions among system components.<br>Technically, DCM is a Bayesian generative modeling and hypothesis testing framework based on state-space models, which infers effective connectivity, latent dynamics and model evidence from observed responses. | $\dot{x} = f(x, u, \theta) + \omega$ $y = g(x, \theta) + \epsilon$ |
| **Free energy (F)** | Free energy quantifies the trade-off between model accuracy and complexity; higher F indicates a better model.<br>Technically, F (also known as the evidence lower bound, ELBO) is a variational functional that bounds the log model evidence from below<br>See **Model evidence**. | $F(Q; P)$ $= E_Q[\log P(y\vert \theta, M)]$ $- D_{KL}[Q(\theta)\vert \vert P(\theta)]$ $\leq \log P(y\vert M)$ $F^{opt} \approx \log P(y\vert M)$ $Q^{opt} \approx P(\theta\vert y, M)$ |
| **Markov blanket (MB)** | Markov blanket refers to the set of variables that fully shield a variable from the rest of the system.<br>Technically, it is the minimal set of variables—comprising parents, children, and co-parents of children—that renders a variable conditionally independent of all other variables in a graphical model. | $P(X\vert rest) = P(X\vert MB(X))$ $MB(X)$ $= Pa(X) \cup Ch(X)$ $\cup Pa(Ch(X))$ |
| **Minimum cut (minCut) problem** | The minCut problem seeks the most economical way to partition a network into disjoint parts.<br>Technically, it is a combinatorial optimization problem that partitions a (weighted) graph into disjoint subsets by minimizing the total (weight of) edges crossing the partitions. | $\min Cut(A, B)$ $Cut(A, B) = \sum_{i \in A, j \in B} W_{ij}$ |
| **Model evidence** | Model evidence measures how plausible a model is given the observed data.<br>Technically, it is the marginal likelihood obtained by integrating the likelihood over the model's parameter prior. Model evidence serves as the cornerstone of Bayesian model comparison. | $P(y\vert M)$ $= \int P(y\vert \theta, M)P(\theta\vert M)d\theta$ |
| **Naïve mean field approximation (MFA)** | The naïve mean field approximation simplifies inference by breaking dependencies between components of a complex system.<br>Naïve MFA is a variational inference method that assumes a fully factorized posterior distribution over parameters, enabling tractable optimization at the cost of neglecting posterior correlations. | $Q(\theta) = \prod_i Q_i(\theta_i)$ |

#### I) Δ*F* relation to Δ*BIC*

In general, the logarithm of the model evidence ratio for the alternative versus the null hypothesis can be written as:

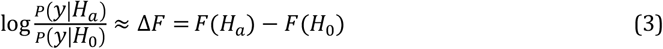

where *F* denotes the variational free energy optimized for each model (i.e. *F* ≈ log *P*(*y*|*model*)) and *y* denotes the observed data. Notably, Δ*F* has the interpretation of a log Bayes factor^6^, which is conventionally taken to provide *strong* evidence for one hypothesis over another when it exceeds 3 nats, and *very strong* evidence when it exceeds 5 nats (Kass & Raftery, 1995). By contrast, the (negative) BIC can be viewed as a special case of the free energy approximation to log model evidence, obtained by discarding all terms that do not scale with the number of data points (Penny, 2012; Penny et al., 2004). This raises the question: under what conditions can Δ*F* be decomposed analogously to Δ*BIC* (Eq. 1), namely

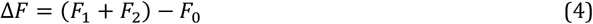

The answer lies in the implicit statistical independence assumption underlying the Δ*BIC* method. If we assume that the two local partitions can be modeled independently, then the joint prior distribution over their parameters can be written as the product of partition-specific distributions; i.e., *P*(*θ*|*H*_*a*_) = *P*(*θ*_1_)*P*(*θ*_2_). Under this assumption, the generative model—as well as the true and approximate posterior distributions—also admit factorized forms under the alternative hypothesis:

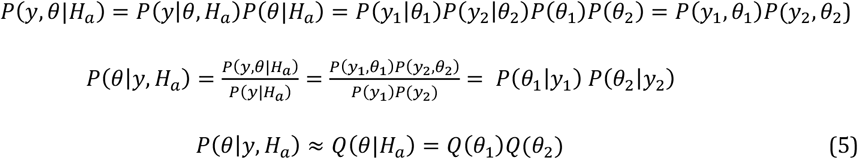

Here, *y* denotes the observed data (i.e. timeseries), *θ* denotes the model parameters and latent variables^7^, *Q*(*θ*) denotes the variational (approximate) posterior, and subscripts denote partition-specific quantities. For a schematic illustration of this factorization, see Fig. 1. Consequently, identifying maximally independent partitions of nodes comes down to optimizing a factorization over subsets of model parameters (i.e. causal connections) so as to maximize model evidence. This observation provides the formal link between Bayesian model comparison and network partitioning that underpins the proposed approach.

**Fig. 1:**
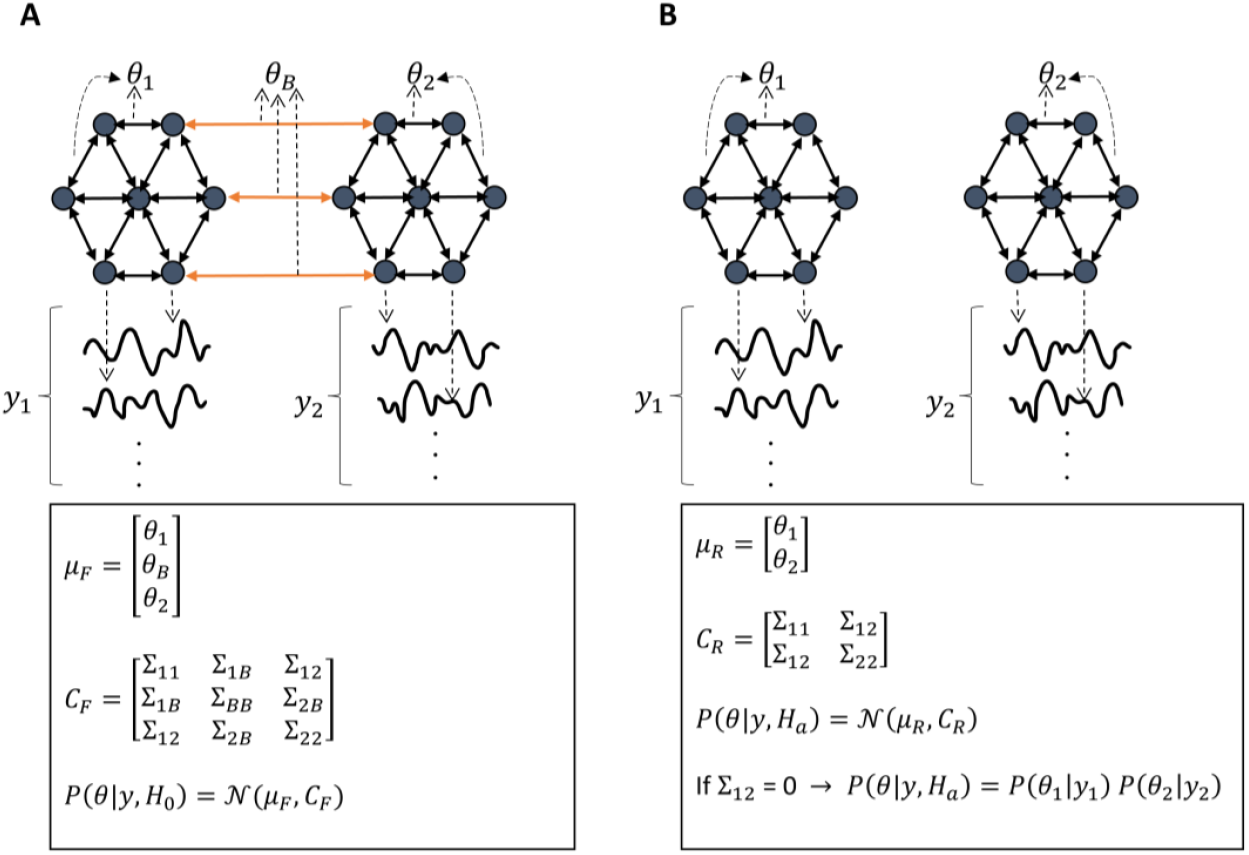
Schematic of the full and partitioned (i.e. reduced) models. (A) A dynamic causal model (DCM) fitted to the timeseries from 14 nodes/regions. Intra-partition connections are denoted by *θ*_*i*_, whereas *θ*_*B*_ represents between-partition connections. The causal connections are normally distributed with mean *μ*_*F*_ and covariance matrix *C*_*F*_. The full model

As shown in Supplementary Appendix A, substituting the factorized model directly into the variational free energy expression reveals that the free energy of the partitioned model decomposes additively as the sum of the free energies of the two partitions; that is, *F*(*H*_*a*_) = *F*_1_ + *F*_2_. This additive accumulation of evidence (akin to Δ*BIC*) is a direct mathematical consequence of modeling local subsets of the data independently under *H*_*a*_. Accordingly, Δ*BIC* can be understood as a special case of the more general formulation in Eq. (3), predicated on an explicit independence assumption. However, the validity of this assumption depends critically on the structure of the underlying data, which raises the question of how violations of independence impact model evidence and performance.

When the independence assumption under *H*_*a*_is well aligned with the true generative process, it leads to an increase in model evidence. This provides the rationale for the success of the Δ*BIC* method in applications such as change-point detection and speech segmentation (Cheng et al., 2008; Chung-Hsien Wu et al., 2006; Costa et al., 2016; Theodorou et al., 2014; Tritschler & Gopinath, 1999; Xue et al., 2010). By contrast, in spatially distributed systems characterized by strong interactions—such as brain networks—assuming fully independent partitions is generally unrealistic. In such cases, this assumption typically reduces model evidence relative to models that explicitly account for cross-partition interactions, yielding a negative^8^ Δ*F*. Importantly, this does not undermine the utility of the approach. Among all candidate bi-partitions, the partition that maximizes Δ*F* (i.e., is least negative) identifies the weakest or most vulnerable breakpoint(s) in the system—those along which the network can be most plausibly decomposed into modules^9^. If subsequent significance testing supports this solution, it constitutes principled statistical evidence for modular organization. Consequently, after computing and maximizing Δ*F*, the next essential step is to assess the statistical significance of Δ*F*^*max*^, as described below. serves as the null hypothesis, *H*_0_. (B) The reduced model, which serves as the alternative hypothesis *H*_*a*_, comprises two DCMs fitted separately to the two partitions. Practically this means that between-partition connections have been removed from the full model, such that the partitioned model is *nested* inside the full model.

#### II) Partition significance

To claim that an integrated system is partitionable (i.e. modular), we need to check whether Δ*F*^*max*^ is statistically significant^10^, at some significance level *α*. This is achieved by assessing whether^11^ Δ*F*^*max*^ ≥ *C*_*α*_, where C_α_ is a critical value for rejecting the null hypothesis, determined from the null distribution of Δ*F*^*max*^ such that:

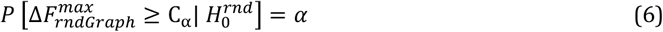

Here 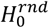 stands for the null hypothesis of optimally bi-partitioning a randomly connected network (with some properties matching the original network). We will discuss how to build the null distribution of Δ*F*^*max*^ (denoted 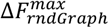 for clarity) later in this section. In summary, our goal is to (1) compute and maximize Δ*F* and (2) accept a bi-partition only when Δ*F*^*max*^ ≥ *C*_α_, or equivalently when the 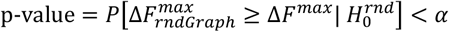. The remaining points below offer pragmatic solutions to achieve these goals.

#### III) Bayesian model reduction (BMR)

The third point concerns the efficient computation of Δ*F*—before attempting to maximize it—for any alternative hypothesis *H*_*a*_. Unlike the Δ*BIC* method, the proposed technique does not require computing *F*_1_ and *F*_2_ by fitting separate models to subsets of the data. Since the structure of any partitioned model is *nested* inside the full (unpartitioned) model, we can use the machinery of *Bayesian model reduction* (BMR) to compute *F*(*H*_*a*_) in one shot and analytically^12^ (Friston et al., 2019, 2016; Friston & Penny, 2011). This is realized by treating the alternative hypothesis as a reduced (i.e. nested) *prior* in the BMR framework, and computing *F*(*H*_*a*_) based on information obtained from fitting the full model—namely, its free energy, prior, and posterior distributions. As shown in Appendix B, this formulation amounts to:

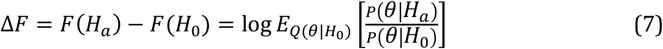

Notably, under Gaussian assumptions on the distributions of the causal connections in variational Laplace, the BMR formulae admit simple forms based on the prior and posterior mean vectors and covariance matrices of the causal connections (Friston et al., 2007, 2019, 2016; Friston & Penny, 2011; Zeidman et al., 2023):

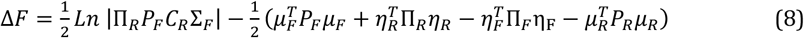

For the notation and a succinct introduction to BMR, please refer to Appendix B, in the Supplementary. Although these expressions admit closed-form solutions, their direct evaluation becomes impractical as network size increases—due to the cost of large matrix multiplications and inversions—which motivates a simplified approximation.

#### IV) Naïve BMR

This point is computationally crucial for large networks. When the network contains many nodes (e.g. n = 1,000), multiplying and inverting covariance matrices (with e.g. *n*^2^ = 10^6^ entries) quickly becomes prohibitive in BMR calculations. In this case, naïve mean-field approximations^13^ (MFA) (Dauwels, 2007; Geiger & Meek, 2005; Jaakkola & Jordan, 1998; Jordan et al., 1998; Khemakhem et al., 2020; Kingma & Welling, 2013; Marković et al., 2024; Xing, 2004; Xing et al., 2002) can be used to factorize the variational posterior and simplify the BMR computations. As demonstrated in Appendix C, MFA leads to a naïve BMR formulation with an intuitive Δ*F* expression that avoids matrix operations:

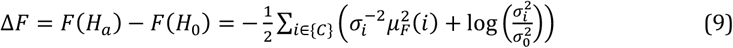

Here, *μ*_*F*_ and 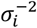 denote the posterior means and precisions of the connections, while 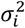 and 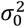 denote their posterior and prior variances, respectively. The summation is performed over the set of removed (i.e. cross-partition) connections, {*C*}. Hence, to maximize Δ*F*, the causal network should be partitioned to minimize the right-hand side of Eq. (9):

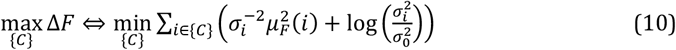

#### V) Δ*F*^*max*^relation to minCut

This point is a bridge to the realm of graph theory, which provides a rich set of optimization tools. Notably, the objective function in Eq. (10) resembles the *minimum cut* (minCut) problem in graph theory (Von Luxburg, 2007), where the *cut size* refers to the sum of between-partition connections (which must be *cut* through to partition the model). In this case, the graph for which we seek the minCut is a function of the original graph, where the connections are squared and precision-weighted 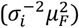, and adjusted to reflect prior to posterior changes in variance (i.e. uncertainty), through log 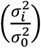. This transformation establishes a direct correspondence between reductions in free energy and graph cut size, the implications of which are unpacked below.

Squaring the connections 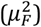 implies that the magnitudes—not signs—of the connections matter, so removing weaker connections minimizes the cut size. Scaling by 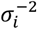 further steers the cut towards less precise connections. Conversely, the log variance-ratio term favors removing overly precise connections, as these contribute disproportionately to the complexity term in free energy and result in unfavorable “brittle” models (Penny et al., 2004). For typical DCM values, the log-ratio term (which is negative^14^) is several orders of magnitude smaller than the positive squared-connection term, rendering it negligible in practice. Thus, solving the minCut problem for the transformed graph comes down to finding and cutting through relatively *weak and uncertain* connections, thereby carving the graph into disjoint partitions.

The well-known challenge with minCut is that it becomes an NP-hard combinatorial optimization problem when the partitions are required to be balanced (Von Luxburg, 2007). Accordingly, a computationally efficient approximate solution— called *spectral clustering*— has been developed. In spectral clustering, the eigenvectors of the graph Laplacian^15^ matrix constitute low-dimensional representations of the graph vertices, a process known as spectral (or Laplacian) embedding. A clustering algorithm then partitions the nodes in this lower dimensional space to minimize a relaxed balanced-cut objective, yielding partitions of comparable size. For a comprehensive review of spectral clustering, see (Von Luxburg, 2007). Here, we intend to apply spectral clustering to the transformed graph defined by Eq. (10). Because classical spectral clustering is applied to symmetric graphs^16^, the transformed graph is symmetrized by adding the transpose of its adjacency matrix to itself—and dividing by two, to preserve the cut size. In summary, applying spectral clustering^17^ to the transformed and symmetrized causal graph provides an approximate solution to the balanced minCut problem, which in turn maximizes Δ*F* (Eq. 10). This achieves our first goal—maximizing ΔF—leaving statistical validation as the final step.

#### VI) Null distribution of Δ*F*^*max*^

Assessing the statistical significance of a candidate partition requires specifying the null distribution of the maximized reduced free energy, Δ*F*^*max*^. Crucially, the key source of stochasticity underlying this distribution does not arise from the free energy itself—which is deterministic for a fixed model and variational posterior—but from variability in network structure. That is, randomizing the connectivity pattern of the graph induces a distribution over model evidences^18^.

Recall that Eq. (10) established a direct relationship between Δ*F* and the cut size of the transformed graph, a correspondence we to refer to as the *free energy-graph cut duality*:

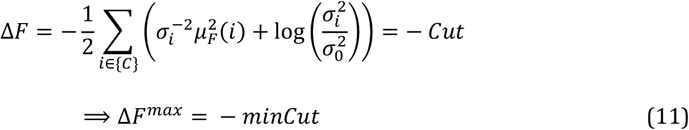

Thus, testing whether Δ*F*^*max*^ ≥ *C*_α_ is equivalent to assessing whether *minCut* ≤ −*C*_*α*_. As we will see, the null distribution of minCut is more straightforward to compute. The intuition is that modular networks are more likely—by design—to exhibit topological bottlenecks^19^ than randomly connected networks. Because minCut solutions typically cut through these bottleneck edges, they often result in *smaller* cut sizes for modular networks than for randomly connected networks with comparable degree and weight distributions. This property can be used to infer modularity—or lack thereof—in the topology of a given network, and to determine stopping criteria in hierarchical partitioning schemes.

The relevant question in the modularity field is called “To cut or not to cut?” (Chang et al., 2014; Reichardt & Bornholdt, 2006), which seeks stopping criteria for graph division based on statistical significance of the modules. Methods that assess the significance of modularity draw either from random matrix theory (Alexander-Bloch et al., 2010; He et al., 2009; Meunier, 2009; Mirshahvalad et al., 2013), network perturbation (Hu et al., 2010; Karrer et al., 2008; Lancichinetti et al., 2010; Mirshahvalad et al., 2012; Seifi et al., 2013), or analytical methods (Chang et al., 2014; Li & Qi, 2020; Newman, 2006; Reichardt & Bornholdt, 2007). Although the statistic used here (minCut) is not identical to modularity (Newman, 2006), their goals and methods are closely related. Below, we describe the randomization procedure used to construct the null distribution of minCut, and comment on potential analytical alternatives in the Discussion.

In brief, we generate a large ensemble of random graphs—connected and matched to the transformed graph in degree and weight distributions—bi-partition each using spectral clustering, and record the resulting cut sizes to form an empirical null distribution of the minCut. Based on this null distribution, we compute the probability of getting minCut values as small as or smaller than the observed minCut. This probability constitutes the p-value, and we adopt the conventional significance threshold *α* = 0.05 to reject the null hypothesis of a nonmodular (i.e. indivisible) graph. Based on the duality in Eq. (11), the same p-value approximately adjudicates the significance of Δ*F*^*max*^. Multiple-comparison correction is then required in the hierarchical setting.

Several approaches exist to control Type I error in hierarchical hypothesis testing. Family-wise error rate (FWER) control can be achieved via resolution-dependent adjustments (e.g. Bonferroni-type corrections) based on the number of hypotheses tested simultaneously (Hochberg & Tamhane, 1987; Meinshausen, 2008; Nichols & Hayasaka, 2003), but this often leads to substantial power loss at finer scales. Alternatively, hierarchical false discovery rate (FDR) control has been proposed at the full-tree, scale-restricted, or outer-node level (Singh & Phillips, 2010; Yekutieli, 2008; Yekutieli et al., 2006), when tests of the parents and their child nodes are independent. A more recent approach that allows for ancestral dependence in the tests controls the average FDR over (all or selected) families of hypotheses (Benjamini & Bogomolov, 2014), where a family^20^ of hypotheses share a common parent hypothesis (Yekutieli et al., 2006). Notably, controlling FDR for each family at level q guarantees that the average FDR across all families is also bounded by q (Benjamini & Bogomolov, 2014). We apply this procedure at each scale, thereby controlling average FDR across all families of hypotheses tested at a given scale^21^ at q = 0.05.

To quantify the magnitude of the modularity effect^22^, we report the standardized effect size (Cohen, 1988), SES = (*X*_*obs*_ − *μ*_*null*_)/*σ*_*null*_, where *X*_*obs*_ = Δ*F*^*max*^, *μ*_*null*_ and *σ*_*null*_ denote the mean and standard deviation of the empirical null distribution of Δ*F*^*max*^, respectively (Botta-Dukát, 2018; De Clerck et al., 2024; Han et al., 2023; Hervías-Parejo et al., 2025; Pérez-Ortega et al., 2023; Tatsumi et al., 2019; Veech, 2012).

This completes the theoretical framework. For a visual summary, the parcellation procedure is presented as a flowchart in Fig. 2. In the following section, we apply this procedure first to simulated examples and then to an empirical causal graph of the brain.

**Fig. 2:**
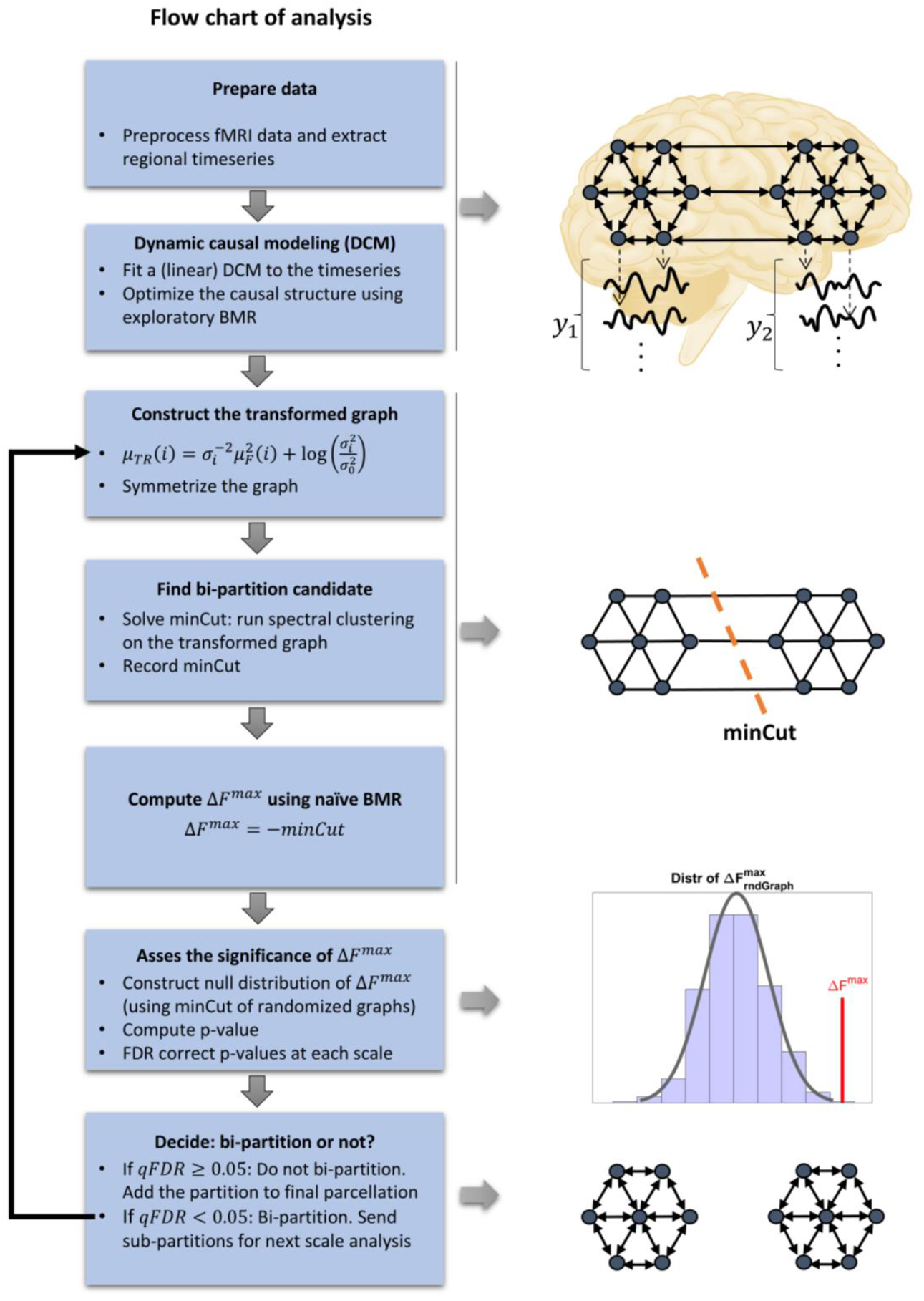
Flow chart of the recursive top-down parcellation scheme. (Brain schematic: Injurymap, https://commons.wikimedia.org/wiki/File:Human_Brain.png, Recolored by Author, CC BY 4.0).

## Materials and Methods

### 3.1 Simulated networks

To demonstrate the functionality of the bi-partitioning scheme, it is initially applied to small simulated causal networks (Fig. 3-A) and subsequently to a large empirical network. The first simulated network, originally used in (Friston et al., 2019), consists of eight interacting nodes with classical state-space dynamics of the form:

**Fig. 3:**
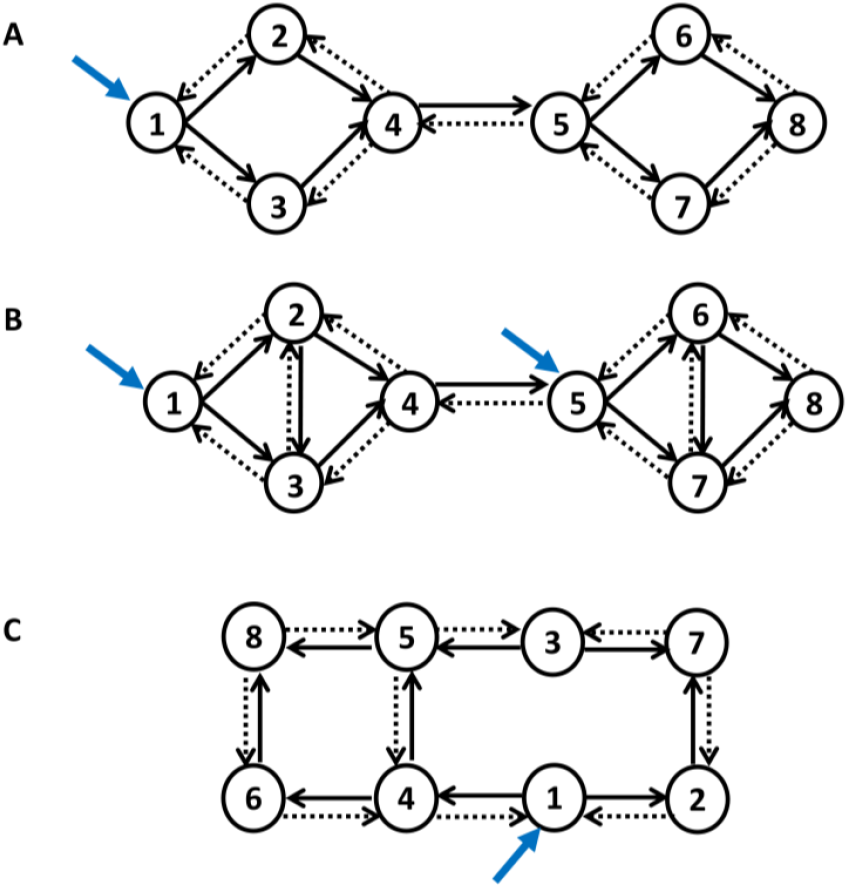
Simulated causal networks. (A) An 8-node causal network that exhibits weak modular structure. (B) A more modular causal network. (C) A randomly connected nonmodular causal network. Solid black arrows represent excitatory connections, whereas dashed arrows signify inhibitory connections. Self-connections are always inhibitory, but are not shown explicitly. Blue arrows represent external perturbations. Synthetic timeseries were generated from each network. A fully-connected DCM was fitted to each network and pruned post-hoc, using exploratory Bayesian model reduction, to yield the optimal (i.e. most accurate and sparsest) causal structure. The estimated model structure and parameters are depicted in Fig. 4 for the first network, and in Supplementary Figs. S1-S2 for the other two networks.

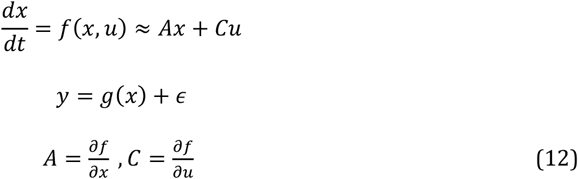

where the time dependencies of *x, y, u* and *ϵ* are implicit, for brevity. The state vector *x* holds a hidden quantity associated with each node. For example, if the nodes represent brain regions, *x*_*i*_ encodes the ensemble neuronal activity of region *i*—which affects and is affected by the activity in the regions connected to it. The general state equation 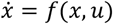 was approximated by a Taylor series in the first line of Eq. (12); hence, the Jacobian or connectivity matrix, *A*_*N*×*N*_, encodes the interactions among the *N* nodes, such that *A*_*ij*_ is the influence from node *j* on node *i*. Moreover, matrix *C* parametrizes the effect of external perturbations applied to the network, encoded as time series in the columns of matrix *u*(*t*). The connectivity parameters *θ* = (*A, C*) are rate constants in units of Hz. The second line of Eq. (12) describes the generation of measurements given the state vector *x*, through an observation function *g* and additive measurement error *ϵ* (Friston et al., 2019).

To generate the data, the connectivity matrix *A* was designed to have excitatory feedforward connections (+0.5 Hz), inhibitory feedback connections (−1 Hz), and self-inhibition on each region for stability (−0.25 Hz). The network was driven by twenty events of one-second duration each that perturbed the first node with temporally jittered onsets (*C*_11_ = 1 Hz). The observation function *g* was set to identity for simplicity, and Gaussian observation noise (with variance = 1/400) was added to the measurements to yield a signal to noise ratio of approximately 8 dB (Friston et al., 2019).

Having generated the synthetic data, the connectivity structure was next assumed to be unknown. To infer it, a fully-connected DCM was fitted to the data, using a variational Laplace scheme (*spm_nlsi_GN*.*m*), which estimated the posterior distribution over the parameters of this full model and a free energy approximation of its log evidence. The full model’s posterior connectivity was then pruned using exploratory Bayesian model reduction (*spm_dcm_bmr_all*.*m*)(Friston & Penny, 2011; Penny et al., 2004). The results are illustrated in Fig. 4. Notably, the partitioning procedure begins at this stage, i.e., *after* optimization of the causal model’s structure and parameters.

**Fig. 4:**
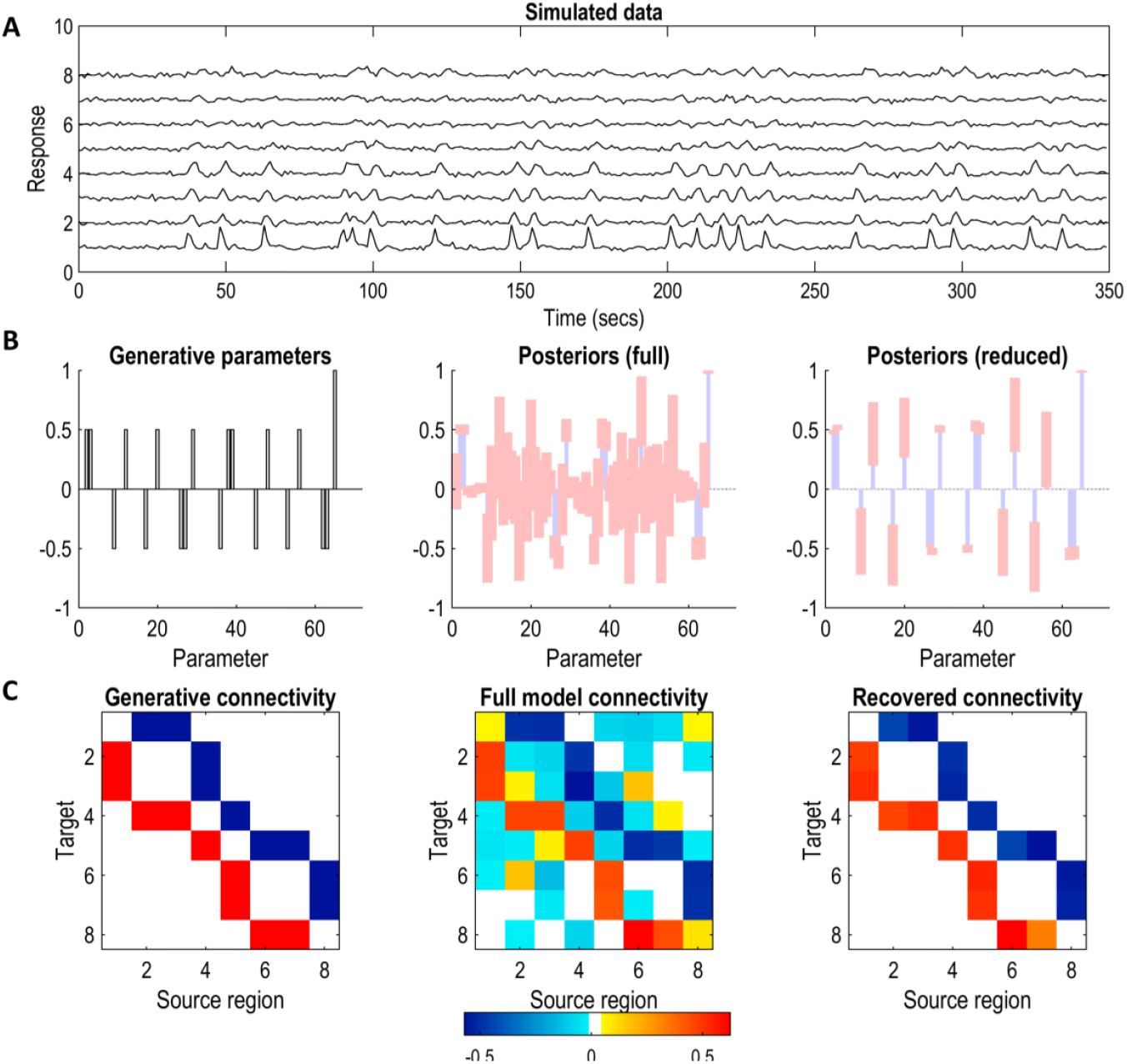
Synthetic data generation and DCM model inversion. (A) Data simulated from the causal model in Fig. 3-A. Each row contains the timeseries from one of the eight nodes. (B) Left: Generative model parameters (i.e. the ground truth connections). Middle: Posterior parameter strengths, from fitting a fully-connected DCM to the synthetic timeseries. Blue bars represent expected values and pink error bars represent 95% credible intervals. Right: Posterior parameter strengths following exploratory Bayesian model reduction. Weak and uncertain parameters have been pruned automatically to improve model evidence. For an unpacked version of the bar plots please refer to Supplementary Fig. S3. (C) Left: Generative (i.e. benchmark) connectivity in matrix format. Red and blue connections represent excitatory and inhibitory effects, respectively. Middle: Connectivity matrix following variational inversion of the fully-connected model. Right: The final connectivity matrix, following Bayesian model reduction. Only significant parameters (with 90% credible intervals not containing zero) are shown. Similar results for the other two simulated networks (Fig. 3-B and Fig. 3-C) are included as Supplementary Figs. S1-S2.

Since the simulated network was sufficiently small, we were able to perform an exhaustive search over the space of all possible bi-partitions, to show that spectral clustering solves the minCut (=max Δ*F*) problem while respecting the balance constraint. In this setting, each possible bisection of the graph corresponds to an alternative hypothesis (*H*_*a*_ is Eq. 3), Practically, generating random bisections is equivalent to performing random cuts on the graph. For each random cut, the associated cut size was converted to a Δ*F* value using Eq. (11)—yielding a set of values denoted by Δ*F*_*rndCut*_. Fig. 5-B shows the distribution of Δ*F*_*rndCut*_, with the Δ*F*^*max*^ value obtained from the spectral cut indicated by a red circle. These histograms illustrate the anticipated peaked, approximately Gaussian form of the random-cut distribution (Schreiber & Martin, 1999)—even for relatively small graphs—and demonstrate the near-optimality of the spectral solution among all possible cuts. Interpretation of these results is revisited in the Results section.

**Fig. 5:**
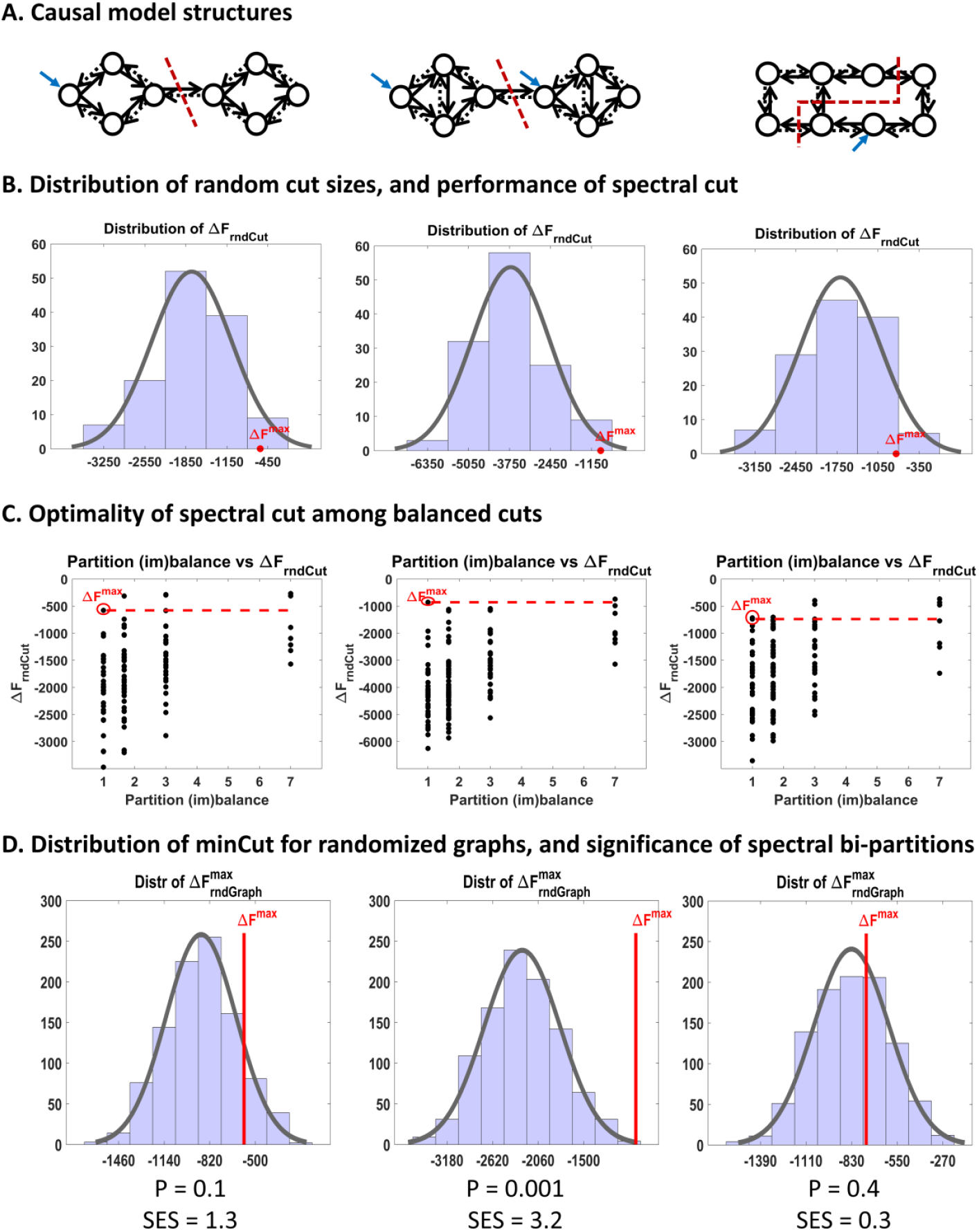
Bi-partitioning of simulated causal networks. (A) Posterior effective connectivity structures of the three simulated causal networks shown in Fig. 3. Each network was bi-partitioned by applying spectral clustering to its corresponding transformed graph (see Theory). Dashed red lines indicate the resulting spectral cut locations. (B) Distributions of *ΔF*_*rndCut*_ obtained by evaluating all possible random cuts of each transformed graph (one column per network). Cut sizes were converted to Δ*F* values using Eq. (11), and Gaussian fits are overlaid for comparison with theoretical predictions (Schreiber & Martin, 1999). In each case, the *ΔF*^*max*^ value obtained from the spectral cut is indicated by a red circle, demonstrating that it is approximately optimal among all possible cuts. (C) Scatter plot of partition imbalance versus Δ*F*, showing that *ΔF*^*max*^ is exactly optimal among balanced cuts (partition imbalance = 1), in these examples. Partition imbalance was defined as the ratio of partition sizes following each cut. Only unbalanced cuts (partition imbalance > 1) yield Δ*F* values exceeding *ΔF*^*max*^. (D) Null distributions of 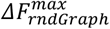 obtained by solving balanced minCut problems (via spectral clustering) for 1,000 randomized graphs matched to the original transformed graph in degree and weight distributions. Cut sizes were translated to Δ*F* values using Eq. (11), and Gaussian fits are overlaid for reference. The vertical red line indicates *ΔF*^*max*^ from the original graph. The associated p-value corresponds to the normalized tail probability to the right of *ΔF*^*max*^, and the standardized effect size (SES) is defined as SES = (*X*_*obs*_ − *μ*_*null*_)/*σ*_*null*_, where *X*_*obs*_ = *ΔF*^*max*^, and *μ*_*null*_ and *σ*_*null*_ denote the mean and standard deviation of the null distribution of *ΔF*^*max*^.

To assess the significance of the spectral bi-partitions, we approximated the empirical null distribution of Δ*F*^*max*^ by generating 1,000 randomized graphs^23^ matched to the original transformed graph in degree and weight distributions while remaining connected. Each randomized graph was bi-partitioned using spectral clustering, and the resulting cut sizes were mapped to Δ*F* values using Eq. (11), yielding the 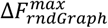 null distribution in Fig. 5-D. This distribution was used to compute the p-value^24^ of the spectral bi-partition as 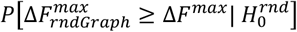 .

The same procedure was applied to a more modular simulated network (Fig. 3-B) and to a randomly connected nonmodular network (Fig. 3-C), to demonstrate that the proposed hypothesis testing framework can reliably distinguish between modular and nonmodular causal structures and appropriately recommend when (not) to partition. The corresponding results are presented in the Results section.

### 3.2. Empirical network

At this point, the multiscale parcellation procedure was applied to a large DCM fitted to empirical fMRI timeseries from the widely used attention-to-visual-motion dataset^25^ (Friston et al., 2021). To summarize, a linearized DCM (Appendix D) was fitted to 1,024 regional timeseries, where each timeseries represented the first eigenvariate from the voxels in the 4 mm vicinity of the local voxel with the highest variance. The Jacobian (i.e. effective connectivity) between regions was then subjected to a Bayesian model reduction procedure, which pruned the redundant connections based on the improvement in model evidence following assumptions of reciprocal local connectivity, and yielded a sparse directed coupling matrix (Fig. 6). This sparse effective connectivity matrix was recursively bi-partitioned by applying spectral clustering to the transformed graph based on Eq. (10), and the partitions at each scale were assessed for significance based on the null distribution of minCut constructed from degree and weight matched randomized graphs (Eqs. 6 and 11).

**Fig. 6:**
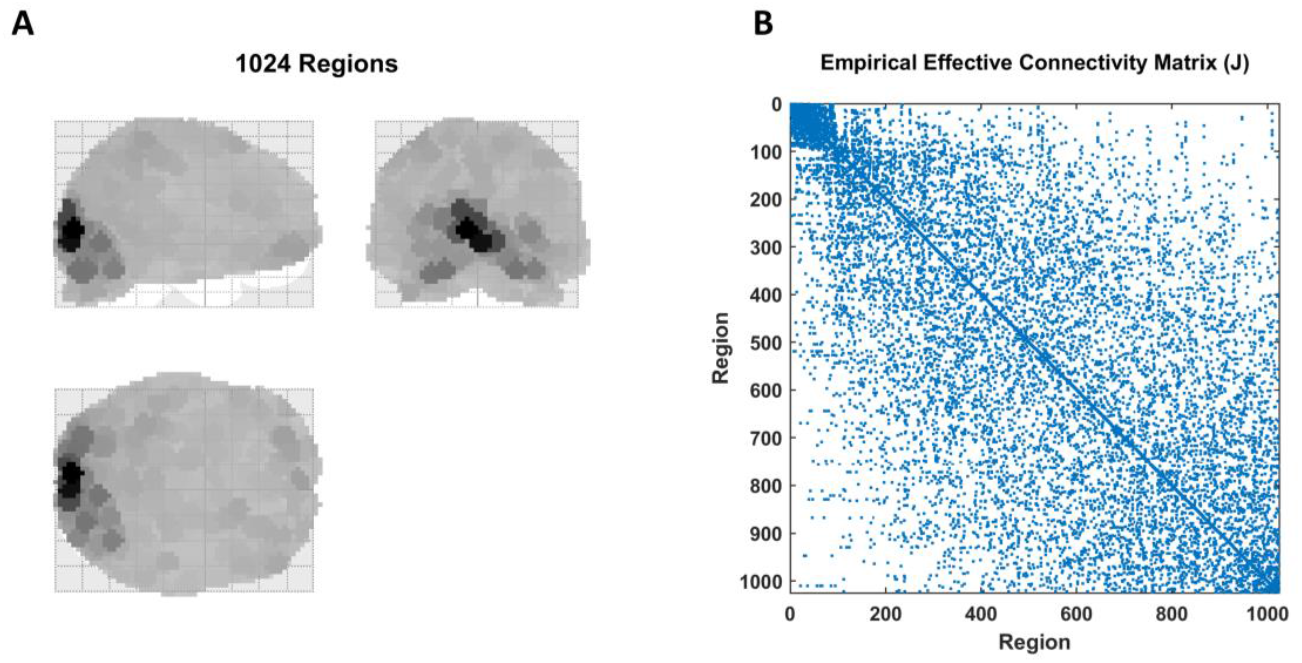
Empirical causal network, inferred from the attention-to-visual-motion dataset, comprising the 1^st^ scale. (A) A maximum intensity projection of the 1024 regions that nearly cover the entire brain, where the spatial support of each region has been color-coded according to the variance explained by its principal eigenmode. (B) Sparsity pattern of the empirical effective connectivity matrix, following DCM inversion and Bayesian model reduction. The matrix is negative definite and structurally symmetric (i.e. bidirectional connections), but asymmetric in coupling values. The connectivities lie in the [-1.82, +1.64] Hz range, but are shown as binary values here to improve the image contrast. This constitutes the model at the first scale of analysis, which was then recursively bi-partitioned using the procedure outlined in the Theory section. The resultant partitions at four consecutive scales have been plotted in the next few figures.

The results, up to the 5^th^ scale, are shown in Figs. 7-9. We elaborate on the findings in the Results section.

**Fig. 7:**
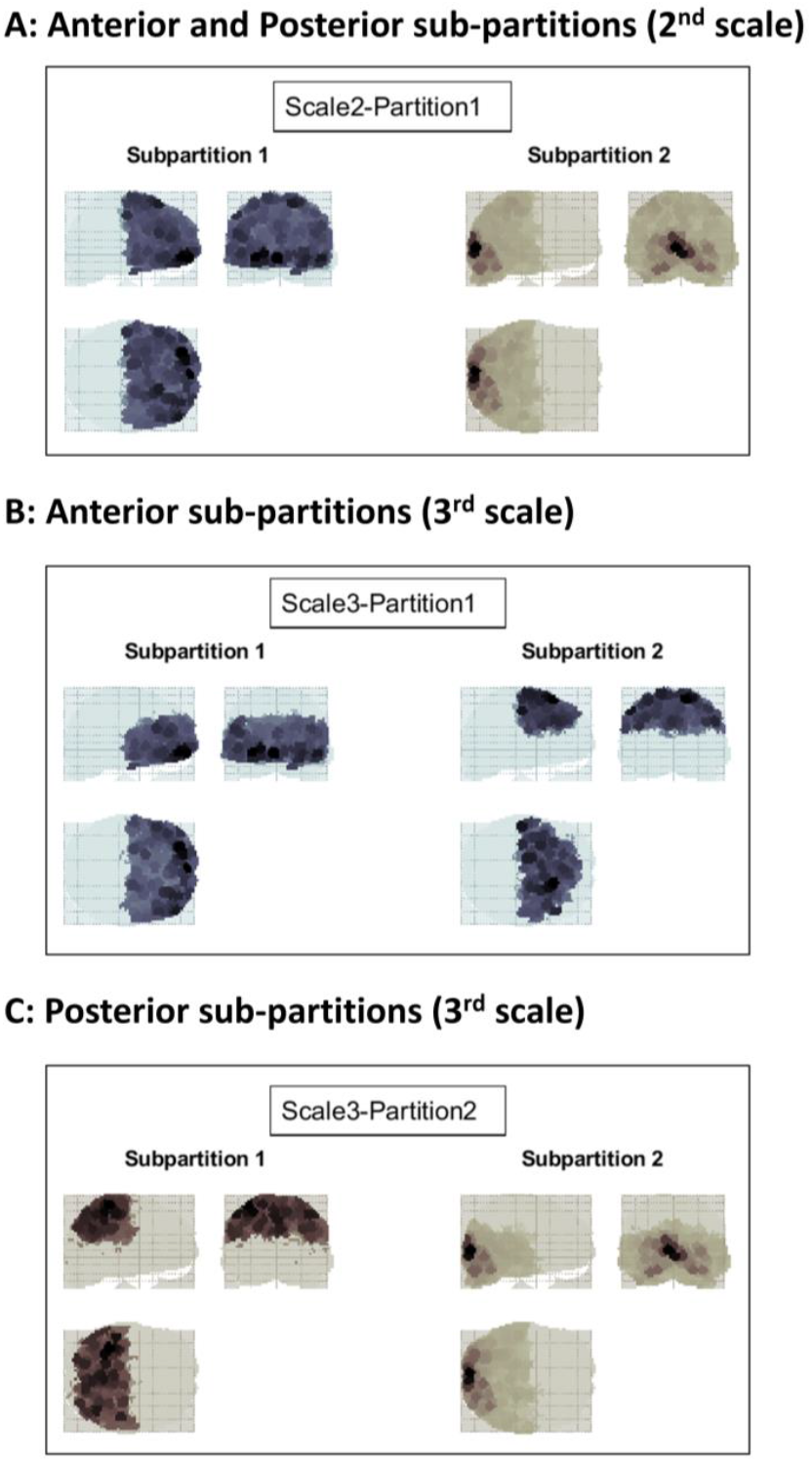
Recursive bi-partitioning of the empirical causal network at the 2^nd^ and 3^rd^ scales. (A) At the 2^nd^ scale, the brain is partitioned into anterior and posterior components. (B) At the 3^rd^ scale, the anterior partition from the 2^nd^ scale is subdivided into superior and inferior components. (C) Also at the 3^rd^ scale, the posterior partition from the 2^nd^ scale is further subdivided into superior and inferior components. In each case, the glass brain shows maximum intensity projections, on the sagittal, coronal and axial planes, where the spatial support of each region has been color-coded according to the variance explained by its principal eigenmode. Each box contains the two subpartitions resulting from one round of bi-partitioning. Each of these subpartitions was further bi-partitioned, as shown in the next figure.

## Results

### 4.1. Simulated networks

Figure 4 shows the model inversion results for the first causal network in Fig. 3-A. Specifically, we show the parameters of the generative model used to simulate the data, the simulated timeseries, the parameter posteriors following DCM inversion, and the posteriors after pruning the network using exploratory BMR. The whole procedure faithfully recovers the original causal structure. Similar plots have been prepared for the other two networks and are provided as Supplementary Figs. S1-S2.

Figure 5 summarizes the bi-partitioning results for the simulated causal networks. In Fig. 5-A, dashed red lines indicate the cut locations obtained by applying spectral clustering to the transformed graphs. For each network, Fig. 5-B shows the distribution of reduced model evidence values arising from random cuts performed on the same graph (Δ*F*_*rndCut*_), computed via the free energy–graph cut duality in Eq. (11). The value of Δ*F*^*max*^ (=−minCut) associated with the spectral cut is marked by a red circle. As expected, Δ*F*_*rndCut*_ distributions exhibit an approximately Gaussian form, consistent with theoretical results showing that random cut sizes converge to normality in the large-network limit (Schreiber & Martin, 1999). Only a small fraction of random cuts yield lower cut sizes than the spectral cut (or equivalently, higher Δ*F* values than *ΔF*^*max*^); however, these correspond to highly unbalanced partitions (e.g., isolating a single node). In contrast, the spectral cut satisfies the balance constraint, as evinced by the cut locations in Fig. 5-A and quantified in Fig. 5-C. The scatter plots in Fig. 5-C further demonstrate that, for these examples, *ΔF*^*max*^ is exactly optimal among all balanced cuts, where partition imbalance was defined as the ratio of partition sizes following each (random) cut.

Figure 5-D shows the empirical null distributions of 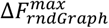 for each causal model, obtained by performing balanced minCuts (via spectral clustering) on randomized graphs matched to the original transformed graphs in degree and weight distributions. Consistent with theoretical expectations, these null distributions are again approximately Gaussian^26^, because minCut distributions for random graphs converge to normality in the large-network limit (Schreiber & Martin, 1999). The p-values reported at the bottom of Fig. 5-D quantify the statistical significance of the observed Δ*F*^*max*^ values relative to their respective null distributions, while the standardized effect sizes (SES) provide a scale-normalized measure of the strength of the modularity effect. Based on these results, only the second model is deemed statistically divisible and modular (p = 0.001, SES = 3.2). In contrast, the first model (p = 0.10, SES = 1.3) and the third model (p = 0.40, SES = 0.3)—obtained by randomizing the connections of the first model—are classified as indivisible at the *α* = 0.05 significance level.

### 4.2. Empirical network

The empirical effective connectivity matrix of Fig. 6 was recursively bi-partitioned, by applying spectral clustering to the transformed graphs, across multiple scales. Figures 7-9 demonstrate the resultant partitions at four consecutive scales, projected on glass brains for anatomical visualization. Fig. 7-A shows the first bi-partition (on the 2^nd^ scale), which divides the causal model into anterior and posterior components of the brain—colored in blue and brown, respectively (here and at subsequent scales)—where color intensity reflects the variance explained by the principal regional eigenmode. The anterior partition was further bi-partitioned, at the next (3^rd^) scale, into superior and inferior portions (Fig. 7-B). Likewise, the posterior partition was further divided into superior and inferior partitions (Fig. 7-C).

At the subsequent (4^th^) scale (Fig. 8), each partition was further divided into either anterior-posterior or lateral-medial components. The same trend continued at the subsequent (5^th^) scale (Fig. 9). Bi-partitions were only accepted if Δ*F*^*max*^ was deemed significant based on the empirical null distribution constructed from graph randomization. As such, the causal model was partitioned top-down over ten scales, until the evidence no longer supported the modularity (i.e. divisibility) of the remaining partitions.

**Fig. 8:**
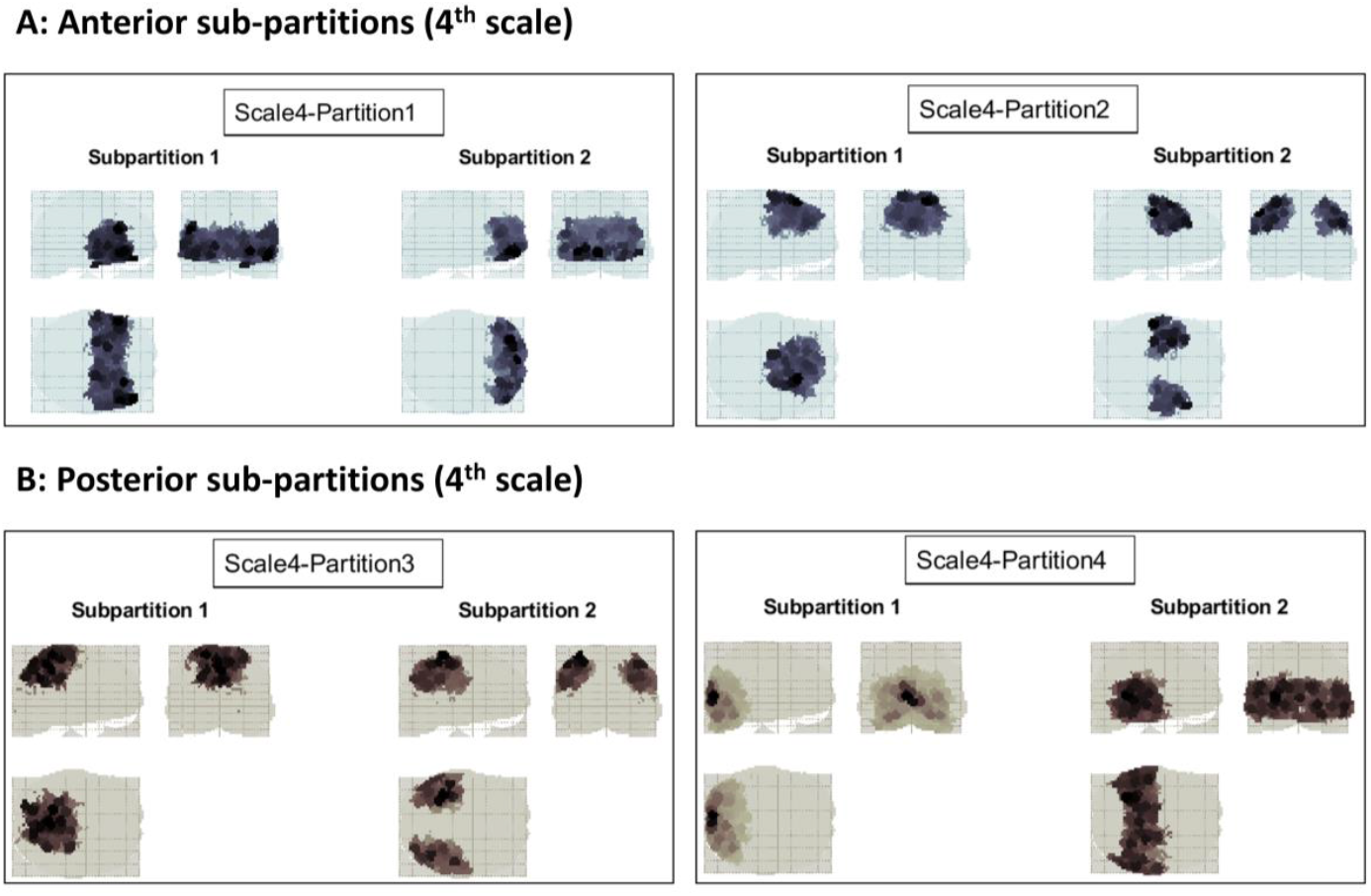
Recursive bi-partitioning of the empirical causal network at the 4^th^ scale. (A) At the 4^th^ scale, the inferior anterior partition of the 3^rd^ scale is further divided into anterior and posterior components, whereas the superior anterior partition is divided into medial and lateral parts. (B) Also at the 4^th^ scale, the superior posterior partition from the 3^rd^ scale is divided into lateral and medial components, whereas the inferior posterior partition is divided into anterior and posterior parcels. Bi-partitioning continues in the next figure.

**Fig. 9:**
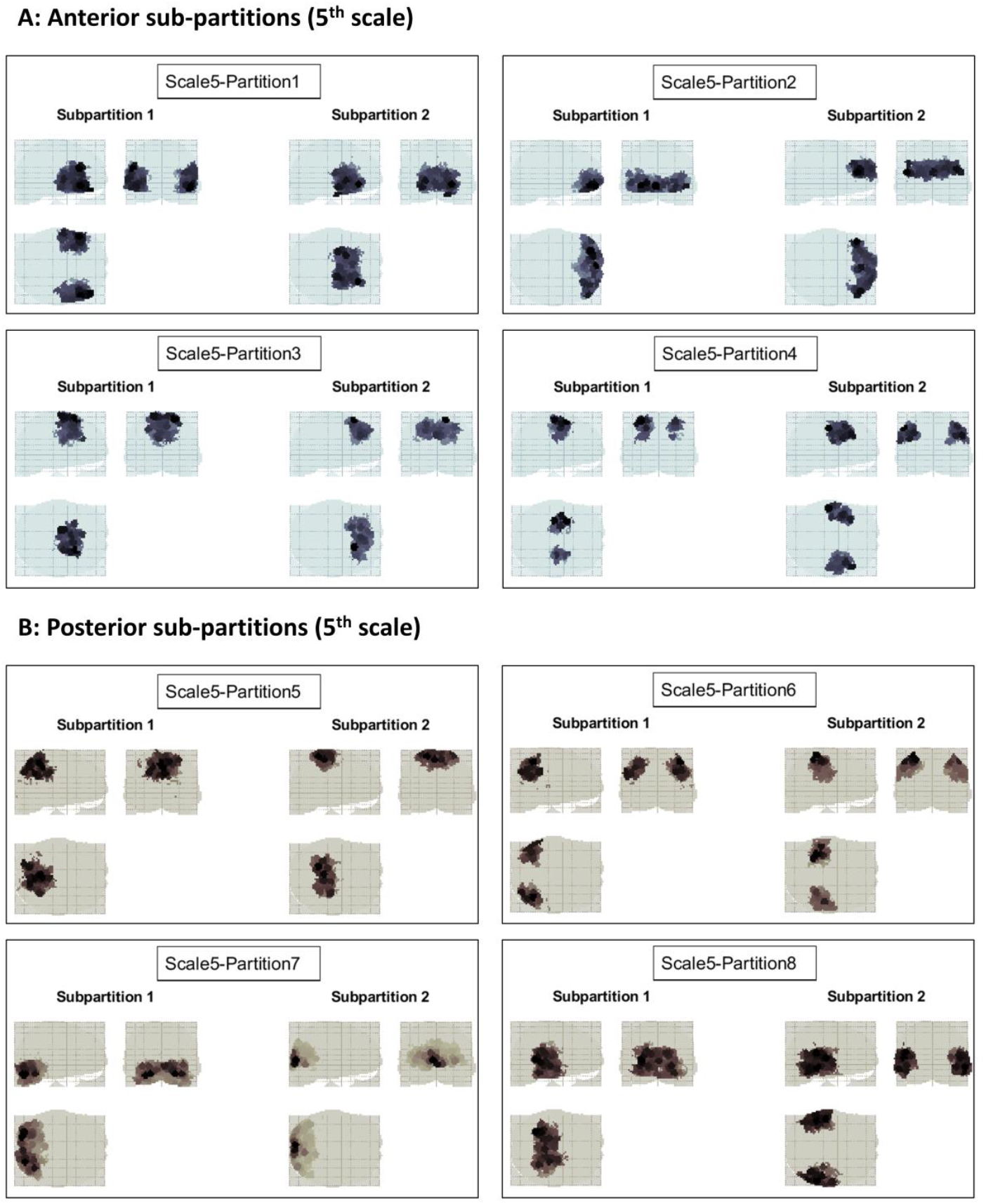
Recursive bi-partitioning of the empirical causal network at the 5^th^ scale. (A) Further bi-partitioning of the anterior brain network. (B) Further bi-partitioning of the posterior brain network. Note how in each case the divisions are either anterior-posterior, lateral-medial or superior-inferior. Initially the number of partitions grows exponentially with scale, but then decreases towards finer scales, where progressively more partitions turn out to be indivisible—as judged by the p-value of the partition significance test.

#### Scale invariance

Figure 10 shows how the standardized effect size, average cardinality (size), dissipative time constant and kinetic energy of the partitions vary across ten scales. The latter two quantities are defined based on the eigenvalues of the Jacobian (i.e. the effective connectivity matrix) in Appendix E. The approximate linearity of the log-log plots is suggestive of power-law relationships and scale invariance—although not definitive (see the Discussion). The same dataset, when parcellated in a bottom-up fashion in (Friston et al., 2021) also exhibited a linear log-log relationship between the expected time constant and the spatial scale. The present top-down decomposition corroborates those findings.

**Fig. 10:**
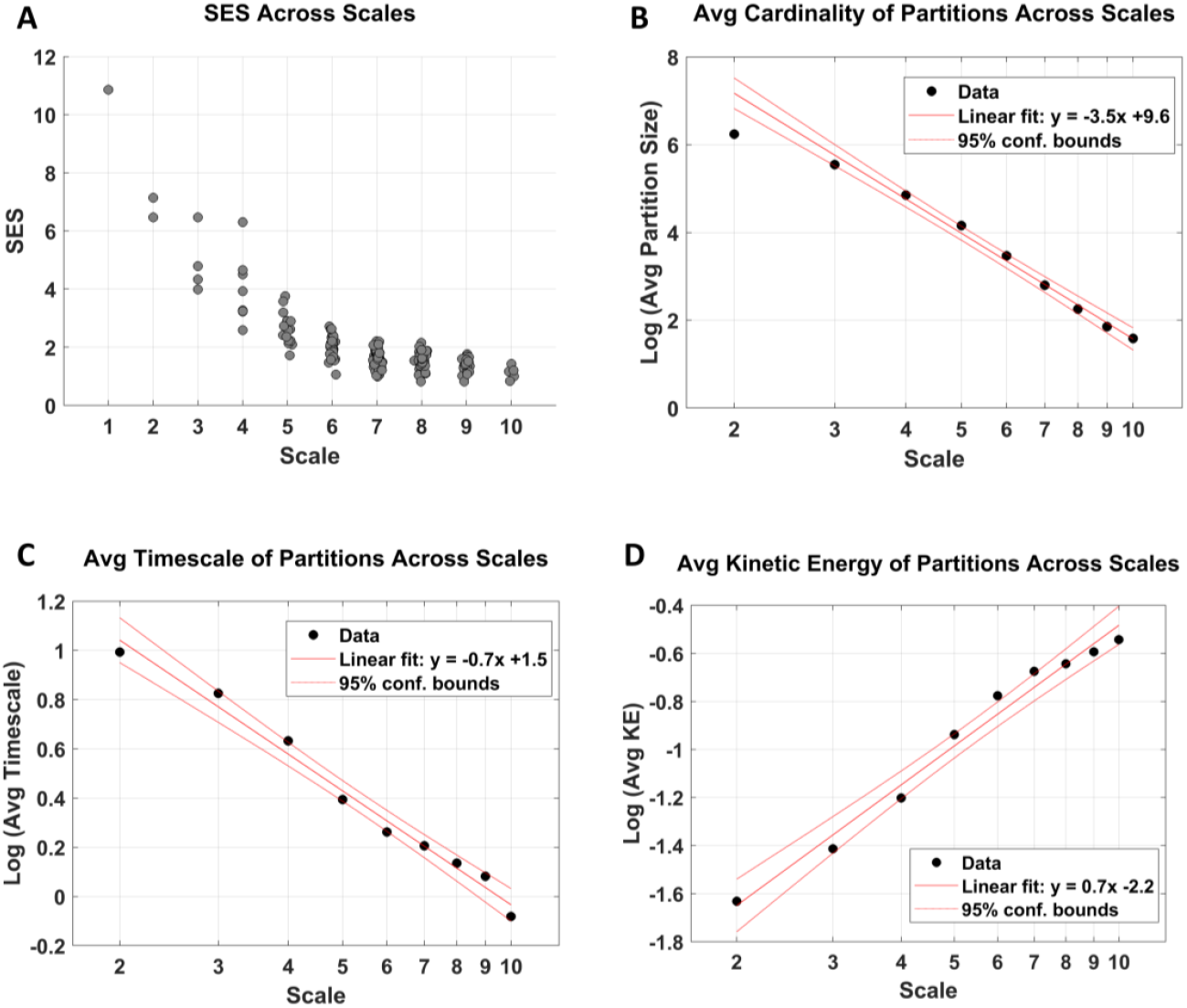
Scaling behavior across hierarchical levels, following multiscale parcellation. (A) Standardized effect sizes (SES) across scales. At finer (higher) scales, partitions are on average smaller and progressively less likely to be further subdivided, as reflected by decreasing SES values. (B) Average partition size (cardinality) as a function of scale. When plotted on log–log axes, the average cardinality decreases approximately linearly. (C) Average intrinsic timescale of partitions across scales, which likewise decreases approximately linearly in logarithmic coordinates. (D) Average kinetic energy of partitions across scales, which increases approximately linearly on a log–log plot. Collectively, these scaling relationships are suggestive of scale-invariant organization across hierarchical levels.

#### Final parcellation and hierarchical depth

The final parcellation of the empirical causal network is shown in Fig. 11-A, where distinct indivisible partitions are indicated by different colors. This finest-scale parcellation corresponds to the leaves of the resulting (dendrogram) tree and yields spatially contiguous parcels. As expected, the multiscale nature of the approach also allows coarser parcellations to be obtained by cutting the dendrogram at earlier scales.

**Fig. 11:**
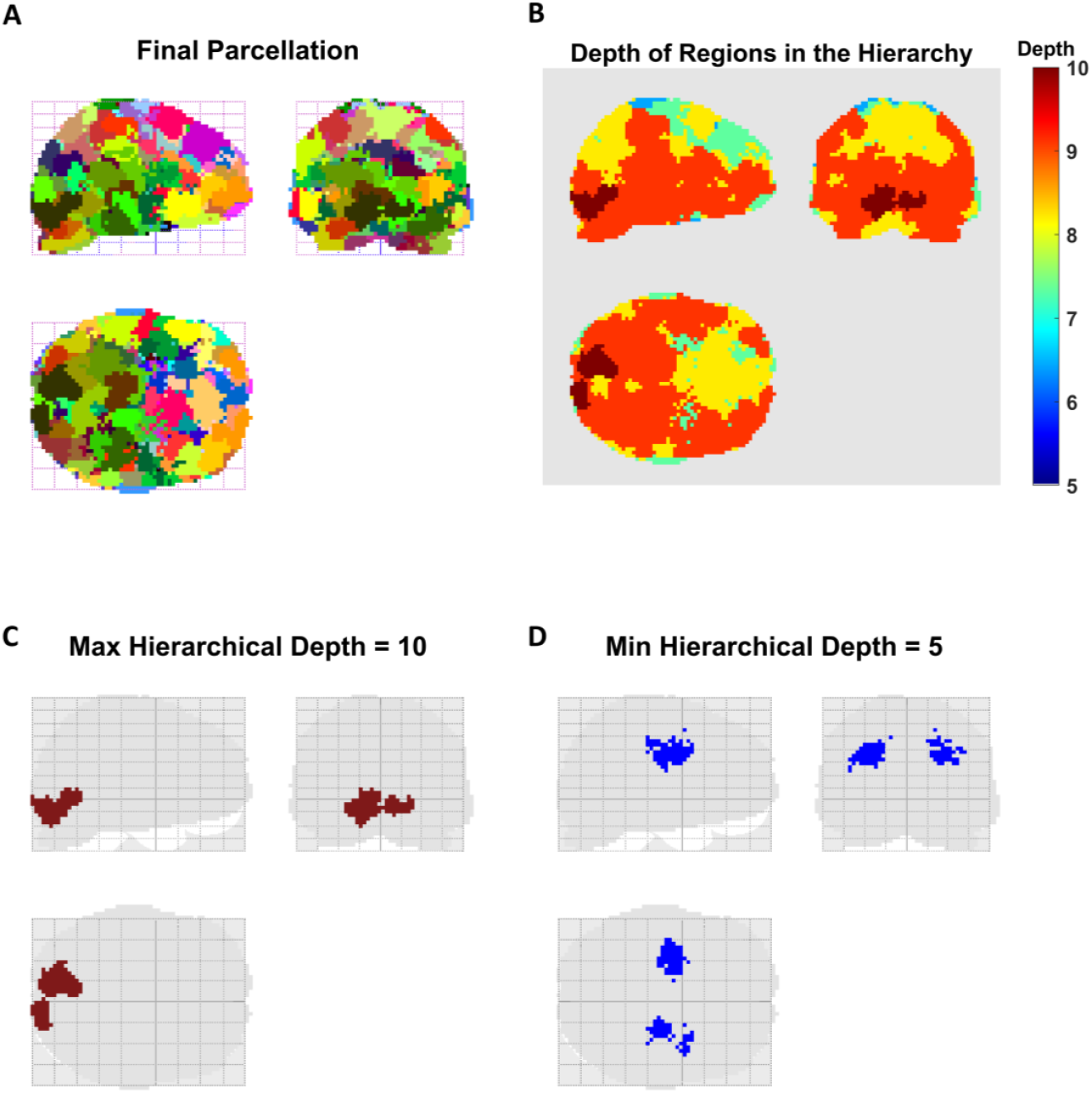
Parcellation results for the empirical network. (A) Finest-scale parcellation of the network, consisting of indivisible partitions (i.e., leaf nodes of the dendrogram), color-coded to distinguish the spatial extent of different parcels. (B) Hierarchical depth of brain regions in the top-down multiscale parcellation, shown using a color scale. Warmer colors indicate regions whose causal structures persist across a greater number of recursive bi-partitioning steps, reflecting their embedding within a deeper multiscale modular organization of the effective connectome. (C) The deepest partition is located in the occipital cortex, whose causal structure passes through ten scales before being identified as nonmodular. (D) The shallowest partition is found in the bilateral medial temporal regions, where an indivisible effective structure is identified by the fifth scale.

The hierarchical depth of brain regions in the top-down multiscale parcellation is visualized in Fig. 11-B using a color-coded representation. Warmer colors indicate regions whose causal structures persist across a greater number of recursive bi-partitioning steps before being identified as indivisible, reflecting the embedding of their local effective connectome within a deeper multiscale modular organization. The deepest partition was located in the occipital cortex (Fig. 11-C), which passed through ten hierarchical levels before being identified as nonmodular. In contrast, the shallowest partition was found in bilateral medial temporal regions (Fig. 11-D), where an indivisible^27^ effective structure emerged by the fifth scale. Overall, for this task and individual, hierarchical depth exhibited a gradual decrease from inferior–posterior to superior–anterior brain regions.

#### Conditional independence and Markov blanket

Another question we can ask of the parcellation results is the conditional independence of the partitions. This question is of interest because recent (simulation) work has suggested that recurrent connections in cyclic causal models can induce statistical dependencies (i.e. functional connectivity) that cross the structural boundaries of the causal model, especially at nonequilibrium steady state (Aguilera et al., 2022). Consequently, the Markov blanket^28^ (MB) identified based on effective connectivity may not coincide with the MB implied by functional couplings. Practically, this means that if certain functional couplings “cross” the boundaries defined by effective connectivity, then the boundary states of the effective connectome are not *the only* MB states (see Fig. 12 for the intuition). This question can be posed for any given dataset and causal model by casting the problem in a hypothesis testing framework:

**Fig. 12:**
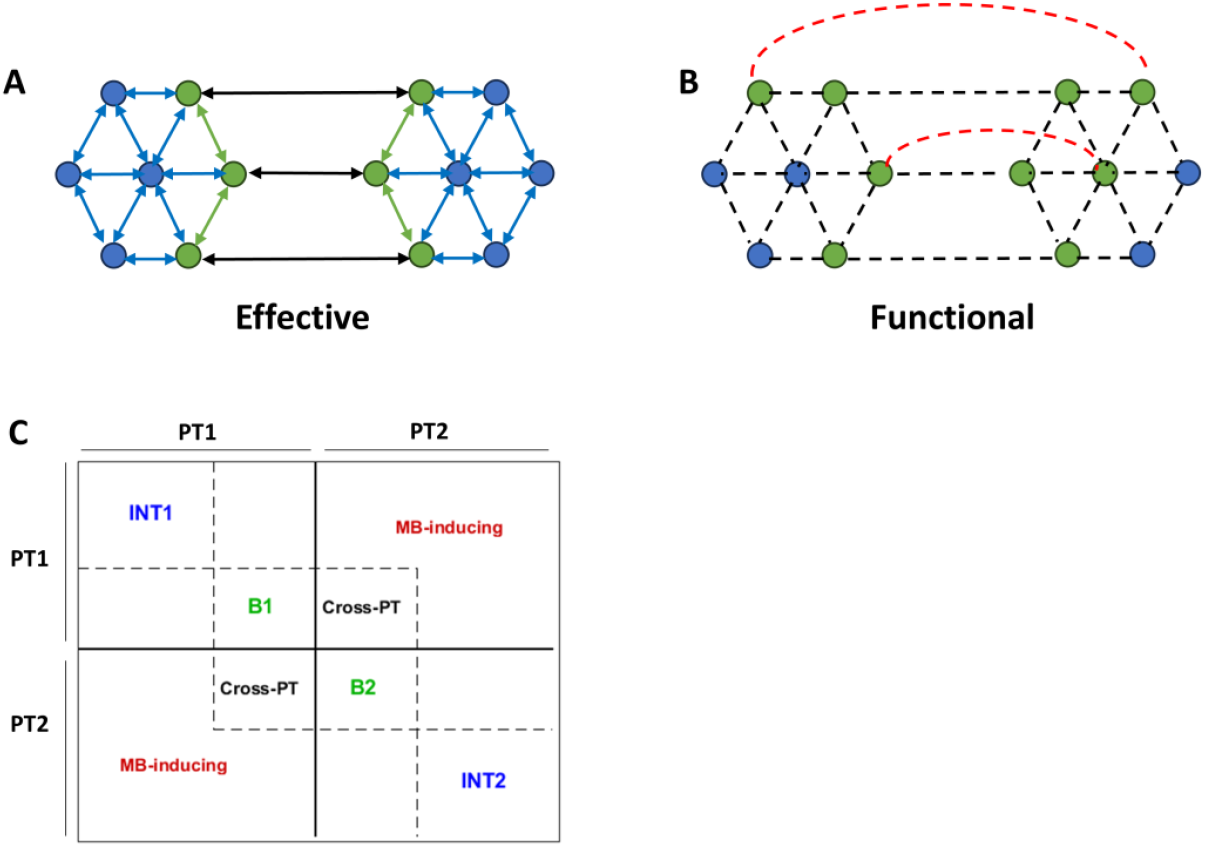
Identifying Markov blanket (MB) states from effective and functional couplings. (A) Schematic of a small directed causal network, comprising two partitions. Directed effective couplings within and between the two partitions are indicated by double-headed arrows. Blue circles denote internal (INT) states, while green circles represent boundary/blanket (B) states. (B) Schematic of functional connectivity for the network in panel A. Undirected functional couplings are shown as dashed lines. If functional couplings (red) are detected between INT1-INT2 states, or between INT2-B1 (or INT1-B2) states, then the corresponding INT states—identified from effective connectivity—must be reclassified as B (i.e. blanket) states at the functional level. (C) Schematic structure of the effective connectivity matrix after reordering states to expose intra- and inter-partition couplings. PT1 and PT2 denote Partition 1 and Partition 2, respectively. INT and B blocks comprise connections among internal states and among blanket states of each partition, respectively, based on effective connectivity. The *Cross-PT* block contains cross-partition connections that mediate interactions between partitions via the effective-connectivity-based boundary states. The *MB-inducing* block contains entries that are zero in the effective connectivity matrix but correspond to functional connections whose presence would violate the conditional independence implied by the effective-connectivity-based Markov blanket, thereby requiring the promotion of the associated internal states to additional MB states. This matrix layout is used to visualize the empirical results in the following figure.

> Having identified a set of Markov blanket (i.e. boundary) states based on the *effective* couplings in a recurrent causal network, what are the odds that some *functional* couplings cross this boundary and thereby induce additional MB states?

To answer this question, we compute and examine the functional network, which can be derived from the effective couplings. The details of computing the inverse covariance (i.e. precision) matrix of the neuronal states 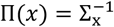 from the effective connectome *J*(*x*), under Gaussian assumptions, are included in Appendix F. After computing the precision matrix (with elements *p*_*ij*_), the partial correlation between any two states is simply computed as:

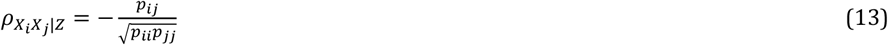

where 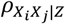 quantifies the correlation between two variables *X*_*i*_, *X*_*j*_, while excluding the effect of a third variable or set of variables *Z*. Under a jointly normal assumption on the distribution of the variables, partial correlation coincides with conditional correlation and can therefore be used to assess conditional (in)dependence between the variables (Baba et al., 2004; Baba & Sibuya, 2005; Lawrance, 1976).

To test whether a population partial correlation is zero, frequentist tests are available (Weatherburn, 1949, pp. 242-263). However, frequentist hypothesis testing cannot quantify the evidence in favor of the null hypothesis. In other words, “the frequentist test does not discriminate between ‘evidence of absence’ and ‘absence of evidence’” (Keysers et al., 2020; Kucharský et al., 2023). This limitation can be addressed within a Bayesian framework through the use of Bayes factors, which allow direct comparison of evidence for competing hypotheses. Here, we adopt a recently proposed Bayes factor test for partial correlations that operates directly on the partial correlation coefficient itself, rather than on proxy statistics such as regression coefficients. For each functional connection, we quantify the evidence in favor of the alternative hypothesis, 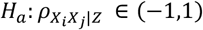, against the null hypothesis,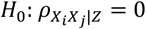. Under the alternative model, the population partial correlation is treated as an unknown parameter that may take any value within the (−1,1) interval, and inference proceeds by estimating this parameter from the data. The resulting marginal posterior distribution of the partial correlation, along with an analytic expression for the corresponding Bayes factor, has been derived by (Kucharský et al., 2023), for a class of well-defined priors. Specifically, the default prior on the partial correlation is a symmetric beta distribution stretched to the interval (−1,1). This stretched beta prior is governed by a single hyperparameter *α*, which controls the concentration of prior mass around zero^29^; representative priors for different values of *α* are shown in Fig. 14-A. To satisfy the information-consistency^30^ criterion for Bayes factors, we restrict 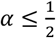 (Jeffreys, 1961; Kucharský et al., 2023). Importantly, the critical functional connections tested in this analysis are restricted to the L-shaped regions in the lower or upper corners of the partial correlation matrix 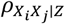 (see Fig. 12-C). These connections are of particular interest because their presence would indicate functional interactions crossing the effective-connectivity-based Markov blanket, thereby inducing additional MB states. For this reason, they are referred to as *MB-inducing* connections.

Figure 13 summarizes the results of applying this procedure to an exemplar empirical effective connectivity matrix (*J*) from the present dataset. Briefly, *J* was first bi-partitioned using spectral clustering on the transformed graph. The inverse covariance matrix 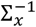 was then estimated from *J* (Appendix F) and converted to partial correlations 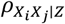 using Eq. (13). A stretched beta prior with default parameter *α* = 0.5 was specified for each partial correlation. Bayes factors comparing the alternative model to the null model (denoted *BF*_10_) were subsequently computed for each functional connection using the formulation of (Kucharský et al., 2023) and reported on the log scale for ease of interpretation. Following conventional thresholds (Kass & Raftery, 1995), log *BF*_10_ ≤ −3 (equivalently log *BF*_01_ ≥ +3) was interpreted as *strong* evidence for the null hypothesis (i.e., absence of functional coupling), while log *BF*_10_ ≤ −1 (equivalently log *BF*_01_ ≥ +1) indicated *positive* evidence for the null model. Conversely, log *BF*_10_ ≥ +3 and log *BF*_10_ ≥ +1 were interpreted as *strong* and *positive* evidence, respectively, in favor of the alternative hypothesis (i.e., presence of functional coupling). The resulting patterns under both thresholds are shown in Fig. 13.

**Fig. 13:**
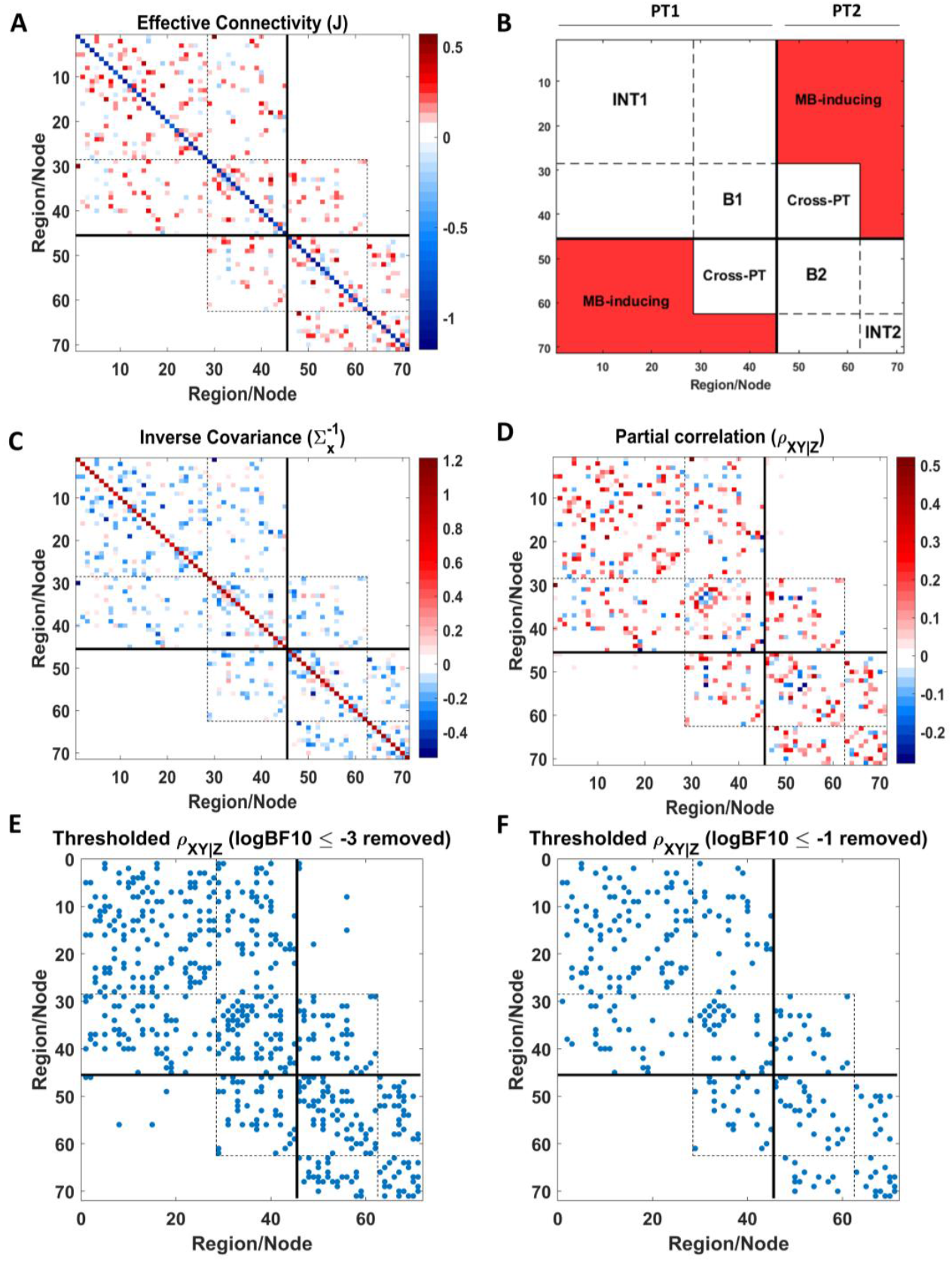
Identification of Markov blanket states in a (reciprocally connected, but asymmetric) empirical effective connectivity matrix. (A) Empirical effective connectivity matrix (71 × 71), bi-partitioned using spectral clustering on the transformed graph (see Theory) and reordered to highlight the modular structure. (B) Schematic indicating the ordering and identity of the states and their interconnections; abbreviations were defined previously in the caption of Fig. 12. (C) Inverse covariance matrix of neuronal states, 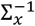, derived from effective connectivity using the relations in Appendix F. (D) Partial correlation matrix computed from 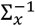 using Eq. (13); diagonal entries (fixed at −1) were removed to enhance contrast. (E) Sparsity pattern of the partial correlation matrix after thresholding (i.e. removing) connections with log BF_10_ ≤ −3 (equivalently, log BF_01_ ≥ +3), corresponding to *strong* evidence for the null hypothesis of zero partial correlation. Bayes factors were computed using a stretched beta prior with hyperparameter *α* = 0.5. Only a small number of connections remain in the MB-inducing region after thresholding, for which evidence in favor of nonzero partial correlation is examined in the next panel. (F) Sparsity pattern after a more liberal threshold, removing only connections with log BF_10_ ≤ −1 (equivalently, log BF_01_ ≥ +1), corresponding to *positive* evidence for the null hypothesis. Although the remaining connections in panel E do not show *strong* evidence for nullity, they do exhibit *positive* evidence for absence. Consequently, the internal states involved in these putative MB-inducing connections are not eligible to be added to the blanket, confirming the validity of the original Markov blanket identified on the basis of effective connectivity. Overall, these results support the conditional independence of the partitions identified by the proposed parcellation method, once conditioned on the effective-connectivity-based boundary states. The robustness of these conclusions to alternative prior specifications is examined in the subsequent figure.

To assess the sensitivity of these findings to the choice of prior, we recomputed *log BF*_10_ across a range of stretched beta priors with *α* < 0.5, thereby preserving information consistency. Figure 14-B shows *log BF*_10_ as a function of 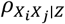 for the connections in the L-shaped MB-inducing region. As *α* decreases, *log BF*_10_ consistently decreases, indicating increasing evidence in favor of the null hypothesis (i.e., absence of these functional connections). This pattern further reinforces the boundary states identified from effective connectivity as the sole Markov blanket states of the system.

These analyses were repeated, over a range of priors, for multiple empirical effective connectivity matrices sampled from different scales of the recursive parcellation hierarchy (results not shown for brevity), yielding qualitatively similar conclusions. Overall, for the present high-dimensional neuroimaging dataset and the assumed (linear, sparse, locally and reciprocally coupled) DCM, the evidence predominantly supports the *absence* of MB-inducing functional connections. This suggests that the Markov blanket states inferred from effective connectivity suffice to establish conditional independence between the recursively identified parcels. While the generality of this finding across datasets and DCM formulations remains to be investigated, the present work provides a principled framework for such future studies.

**Fig. 14:**
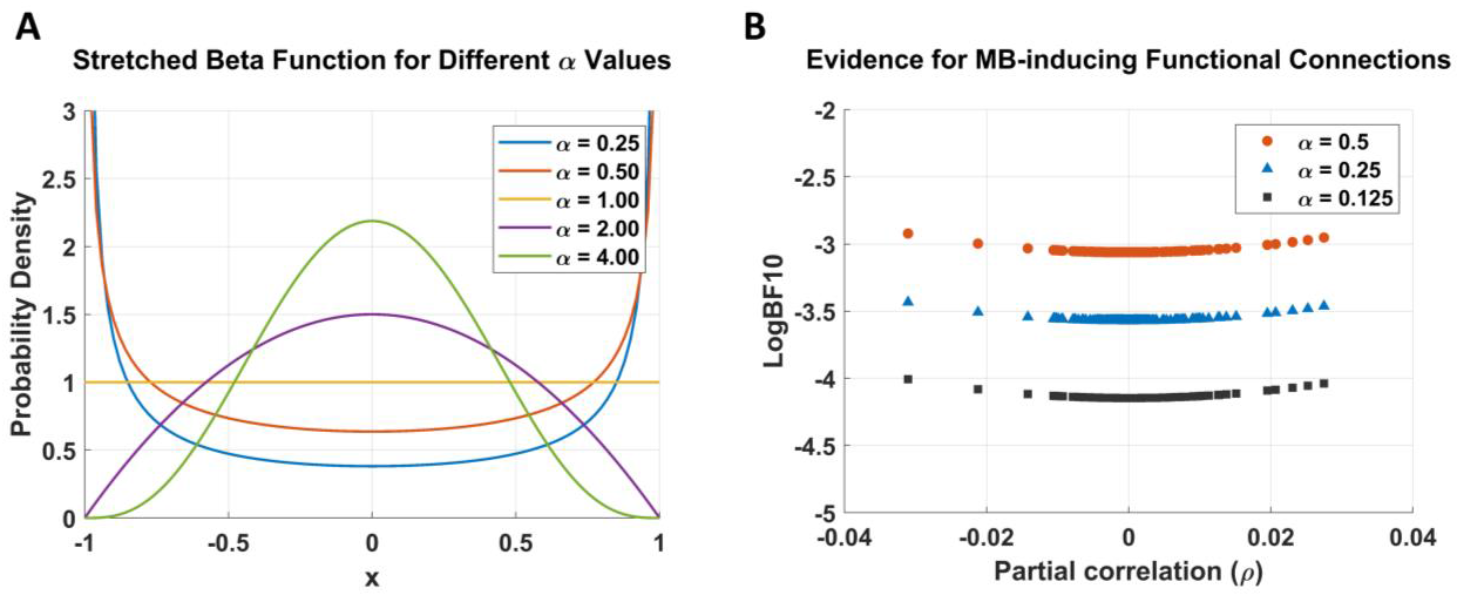
Sensitivity of inference on MB-inducing functional connections to prior specification. Stretched beta priors on the partial correlation, shown for different values of the hyperparameter *α*. (B) *log BF*_10_ as a function of the partial correlation, computed for the connections identified in the L-shaped MB-inducing region in Fig. 13 across several prior settings. All displayed hyperparameter values (*α* ≤ 0.5) satisfy the information-consistency requirement for Bayes factors. As *α* decreases, *log BF*_10_ shifts further towards negative values, indicating increasing evidence for the *absence* of boundary-crossing (i.e. MB-inducing) functional connections, and hence the *absence* of additional MB states.

## Discussion

A top-down recursive parcellation scheme was proposed for (bi)partitioning dynamic causal models. This approach is a generalization of the Δ*BIC* method, framed as Bayesian model comparison. However, unlike the Δ*BIC* method, the proposed technique does not require fitting local models to subsets of the data in order to compute Δ*F* for model comparison. Instead, model evidence for the partitioned model is computed analytically from the full (unpartitioned) model using Bayesian model reduction—because the former model is nested inside the latter. Specifically, this nesting enables analytic evaluation of reduced model evidence without re-estimating parameters.

Furthermore, a naïve version of BMR was introduced, which avoids matrix multiplications and thus considerably reduces the computational cost for large networks. Under the naïve formulation, we derived an analytical relationship between Δ*F* and the graph cut size for a transformed version of the original graph (Eq. 11: Δ*F* = −*Cut*). This duality facilitated both the optimization of Δ*F* and the assessment of its significance, using tools from graph theory. Specifically, we chose spectral clustering to solve the balanced minCut problem, and thereby indirectly maximized Δ*F*. Moreover, the distribution of minCut under the null hypothesis of randomly connected graphs served as a proxy for the null distribution of minus 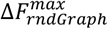, which was subsequently used to assess the significance of Δ*F*^*max*^ and determine whether the causal network was partitionable. The recursive application of this procedure yielded a multiscale hierarchical decomposition of the network. This procedure was applied to simulated causal networks and to a high-dimensional causal network inferred from a neuroimaging dataset. In the following, we review the key findings and highlight important aspects of the proposed method.

In the simulations, top-down parcellation correctly identified the location of the minCut for each causal network using spectral clustering on the transformed graph. Crucially, the parcellation outcome depends on the structure and uncertainty of the *estimated* network, not the ground truth—to which we do not have access in real situations. This emphasizes that the method operates on the inferred causal structure rather than the assumed ground truth, aligning with the longstanding debate of whether a “true” model exists or whether some models are simply more useful (i.e., have higher evidence) than others (Litvak et al., 2019). Although in the three small simulated networks the posterior structures were faithful to the benchmarks, in general the accuracy of the recovered network depends on the size of the network, the amount of data, the type of DCM, and the difficulty of model inversion (i.e., ill-posedness and local optima). In any case, the proposed parcellation scheme is applied only *after* the causal model’s structure and parameters have been optimized—to inspect its hierarchical modular organization.

A second point concerns the use of spectral clustering as an approximate solution to the balanced minCut problem. In general, minCut is a combinatorial optimization problem, requiring brute-force search over all possible bi-partitions, the number of which grows exponentially with network size^31^. Spectral clustering is therefore typically used as a computationally tractable approximation. Importantly, the present framework is agnostic to the specific minCut solver employed. It is possible to improve upon this initial solution using incremental refinements, akin to hill climbing algorithms (Li et al., 2014). More recent methods can even solve minCut exactly in near-linear time (Henzinger et al., 2024).

The third point pertains to the free energy-graph cut duality in Eq. 11. This relationship was derived under the naïve mean field approximation (MFA) (Appendix C) and proved useful for both optimization and statistical inference. Beyond its computational utility, this duality establishes a conceptual bridge between Bayesian model evidence and graph-theoretic modularity. The graph theoretical link is also noteworthy because prior work in integrated information theory (IIT) has shown *numerically* that spectral clustering can plausibly approximate the minimum information bi-partition (MIB) of a network when applied to correlation matrices of neural timeseries (Toker & Sommer, 2019). Based on thus-identified MIB, the authors proceeded to compute the “geometric” version of integrated information, defined as the KL-divergence between the distributions over temporal evolutions of the full and disconnected models (Oizumi et al., 2016; Toker & Sommer, 2019). In comparison, we also used spectral clustering for bi-partitioning the network, but since the present approach is model-based, the (transformed) effective connectivity matrix was used instead of the correlation matrix. We further provided an *analytical* relationship between the cut size and the change in model evidence (Δ*F*) from disconnecting the full model. Future work can compare the MIBs identified using the causal model (i.e. effective connectivity) against those found based on the correlation matrix (i.e. functional connectivity). A potential advantage of the former is that a model-based framework offers a standard measure (Δ*F* = log Bayes factor) to score and compare the plausibility of different bi-partitions against each other.

Note that without naïve MFA, reduced model evidence has the general form of:

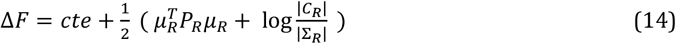

where the terms associated with the full model have been lumped into a constant (cte); *μ*_*R*_ and *P*_*R*_ denote the posterior mean and posterior precision of the retained (i.e. within-partition) connections, while *C*_*R*_ and Σ_*R*_ denote their posterior and prior covariance matrices, respectively. So, so maximize Δ*F*, the model has to be partitioned to maximize the contribution of within-partition connections (i.e. ∑_*i,j*∈\{*C*}_ *μ*_*i*_*μ*_*j*_*P*_*ij*_) while penalizing overly precise configurations via the log-ratio term. This objective still closely parallels the goals of modularity maximization and graph partitioning (Wang et al., 2015) in network science, and suggests that graph partitioning algorithms can provide useful approximations or initial solutions, even beyond the naïve MFA setting, with subsequent refinement to maximize Δ*F*.

The fourth point concerns the statistical significance of the identified partitions. We constructed an empirical null distribution of minCut by applying spectral clustering to a large ensemble of randomized graphs that were connected and matched to the transformed graph in both degree and edge-weight distributions. This permutation-based strategy provides a nonparametric reference distribution that is well matched to the empirical network topology, but can become computationally demanding for very large networks, motivating the consideration of parametric alternatives. Progress in this direction requires a clearer understanding of the distribution of minCut values in random graphs. Early work on binary random graphs established bounds and expectations for the minCut (Bui, 1983; Fu & Anderson, 1986; Kanter & Sompolinsky, 1987), and subsequent work showed that, for sparse random graphs at large network sizes, the minCut distribution converges to a Gaussian^32^ whose mean and variance scale linearly with the number of vertices, *N* (Schreiber & Martin, 1999). Notably, as *N* → ∞ in this asymptotic regime, the relative fluctuations of the minCut about its mean vanish, a property known as “self-averaging”^33^ (Schreiber & Martin, 1999). We showed these properties numerically on our randomized graphs, in supplementary Fig. S4^34^.

Even though these numerical results provide useful intuition, rigorous parametric testing requires further theoretical work to generalize^35^ existing results on minCut statistics to sparse *weighted* random graphs, potentially for different edge-weight distributions^36^. Alternatively, a non-graph-theoretical route to parametric testing could focus directly on the null distribution of Δ*F*^*max*^, which is itself a log Bayes factor. Relevant insights may be drawn from (Zhou & Guan, 2018), who showed that under the null hypothesis, 2log BF is distributed as a weighted sum of chi-squared random variables with a shifted mean, for linear regression models; and that the p-value associated with such BF can be computed analytically—without permutations—using the polynomial method of (Bausch, 2013). Notably, the analytic form of Δ*F* in Eq. (9) resembles the structure of the log BF expression in Eq. (10) of (Zhou & Guan, 2018), suggesting a promising avenue for future development.

Once a parametric form for the null distribution of minCut or Δ*F*^*max*^ is established, one could perform (frequentist or Bayesian) parametric null hypothesis testing. A Bayesian treatment is particularly appealing here, as Δ*F* is itself a log Bayes factor and naturally admits probabilistic interpretation. Bayesian inference additionally requires the specification of priors but has the intrinsic advantage of allowing one to quantify evidence in favor of the null hypothesis, rather than only against it in frequentist testing (Friston, Glaser, et al., 2002; Friston, Penny, et al., 2002; Friston & Penny, 2003; Masharipov et al., 2021; Morey & Rouder, 2011).

Fifth, we review the empirical findings. Recursive bi-partitioning of the empirical causal network revealed cohesive parcels across all ten hierarchical scales. At coarser scales, the spatial maps showed that the divisions followed anterior–posterior, superior–inferior, or lateral–medial axes, as illustrated in Figs. 7–9. Interestingly, several of these partitions closely resemble parcels obtained using the bottom-up (i.e., merging) scheme introduced by (Friston et al., 2021), which is grounded in renormalization group theory. Although the top-down and bottom-up approaches adopt different inferential perspectives, the reproducibility of multiple parcels across methods suggests that both are likely uncovering the same underlying modular organization. This convergence provides some construct validity for the resulting effective parcellation.

Notably, the finest parcellation comprised spatially contiguous parcels (Fig. 11-A). This property arises in part from the locality assumption imposed on the empirical effective connectome, the spatial range of which was optimized in (Friston et al., 2021). More generally, effective connectivity models can be constrained or informed using structural or functional priors. Although structural, functional, and effective connectivities are defined and estimated quintessentially differently (Friston, 2011; Moghimi et al., 2022), there is substantial evidence that they can meaningfully inform one another (Deco et al., 2013, 2014; Greaves et al., 2025; Razi et al., 2017; Sokolov et al., 2019; Stephan et al., 2009). In this context, it would be of interest to examine how such structural or functional priors influence the resulting multiscale effective parcellations within a Bayesian model comparison framework, and whether they enhance anatomical and functional plausibility, particularly at finer spatial scales.

Beyond the spatial organization of the resulting parcels, examining the scaling behavior across hierarchical levels revealed several systematic trends. In particular, the modularity effect size (SES) generally decreased towards finer scales, indicating that smaller partitions were progressively less likely to be deemed further partitionable. In addition, when expressed in logarithmic coordinates, both the average parcel size and the average intrinsic timescale decreased approximately linearly with scale, whereas the average kinetic energy increased linearly across scales, collectively suggesting the presence of scale-invariant organization. Importantly, these trends were observed consistently across multiple summary dynamical measures, and mirror findings obtained previously using a bottom-up parcellation approach (Friston et al., 2021).

Nevertheless, linearity in log-log plots constitutes a necessary but not sufficient condition for genuine power-law behavior. Reliable detection and characterization of power-law distributions typically require data spanning at least two orders of magnitude along both axes, with sufficiently dense sampling across that range (*n* ≥ 50) (Clauset et al., 2009; Stumpf & Porter, 2012). The current dataset and hierarchical depth did not meet these criteria; accordingly, the observed log–log linear trends should be interpreted as suggestive rather than conclusive evidence of scale invariance. This caution is especially pertinent when interpreting scaling behavior in finite, task-specific hierarchies. A more rigorous investigation of power-law behavior in deeper hierarchies would likely require starting from voxel-level effective connectivity rather than parcel-level representations, ideally using datasets with higher spatial resolution.

Another notable empirical observation was the marked inhomogeneity in the hierarchical depth of causal structures across the brain (Fig. 11-B). This indicates that different regions participate in the multiscale causal hierarchy to markedly different extents, with occipital cortex embedded deepest in the hierarchy for the present visual task. More generally, a gradual decrease in hierarchical depth was observed from inferior-posterior to anterior-superior regions of the brain—for this specific task and subject. This apparent spatial gradient raises the possibility of applying gradient-based analyses (Bernhardt et al., 2022; Haak et al., 2018; He et al., 2024; Huntenburg et al., 2018; Margulies et al., 2016; Nashed et al., 2025; Royer et al., 2024; Vos de Wael et al., 2020) to identify the principal axes underlying smooth spatial transitions in hierarchical depth, within condition-specific effective connectomes. Clearly, assessing the robustness and generality of such analyses would require group-level inference, which we will discuss shortly.

Sixth, additional hypothesis testing on the empirically derived parcels revealed that the evidence predominantly supported the absence of boundary-crossing (i.e. MB-inducing) functional connections (Figs. 12–13). In the present high-dimensional neuroimaging dataset, modeled using a sparse, locally and reciprocally coupled linear DCM, this implies that the boundary states identified from the cyclic effective connectivity structure were, in most cases, the only Markov blanket states of the system. Under the assumed causal model, this finding supports the conditional independence of the identified partitions given the Markov blanket states inferred from effective connectivity.

Importantly, the framework introduced here is sufficiently general to enable systematic investigation of this phenomenon across different datasets and modeling assumptions. For example, one may vary the type of data, the number of regions, or the form of the generative model (e.g. nonlinear DCMs), as well as examine the effects of coupling sparsity and asymmetry on the emergence of boundary-crossing functional connections. For instance, simulation work by (Aguilera et al., 2022) showed that increasing asymmetry in effective couplings—thereby driving the system further from equilibrium—can promote functional dependencies that cross the boundaries implied by effective connectivity. While the authors suggested that such effects may challenge the existence of a Markov blanket in cyclic systems, an alternative interpretation is that they reflect the presence of a more extended Markov blanket than that prescribed by effective connectivity alone. Disentangling these possibilities is precisely the type of question that can be addressed within the proposed hypothesis testing framework grounded in causal modeling.

Seventh, although we restricted the recursive parcellation to bi-partitions, it is straightforward to generalize to multi-partitions, in which case the model comparison would be conducted for *H*_0_: *n* = 1 versus *H*_*a*_: *n* ≥ 2 partitions. Note that the derivations for Δ*F* using naïve BMR and the Δ*F* − *Cut* duality do not depend on the *number* of cuts (which is more than one in multi-partitioning), so the same framework would still be applicable. The only difference is that we would have to conduct spectral clustering for a range of *n*′*s* ≥ 2 and compare the resulting Δ*F*(*n*)’s to choose the optimal number of sub-partitions.

Eighth, the derivations of naïve BMR are generic and can be used for purposes other than parcellation as well. We mention two such applications. For instance, if the goal is to prune an over-parametrized Bayesian network, Eq. (9) says that we can *sort* the causal connections based on how much free energy improvement each removal would bring about. Moreover, to compute the free energy improvement from removing a *group* of connections, we just have to add up the analytically computed individual improvements. Another potential application of naïve BMR is in conducting group Bayesian analyses using a random effects scheme known as parametric empirical Bayes (PEB)(Friston et al., 2016, 2016; Zeidman et al., 2019). Since PEB formulation relies on BMR, naïve BMR can be specifically useful for performing group analysis over large networks—by avoiding matrix operations. Note that the whole top-down parcellation scheme is also generic and can be used for different data types—other than neuroimaging—and different DCMs developed for other applications (Friston et al., 2022).

In future work, we aim to extend the proposed framework to perform multiscale parcellation at the group level within a random-effects setting. One natural approach is to pool individual posterior estimates using PEB and subsequently parcellate the resulting group-level network. Alternatively, modularity assumptions could be integrated into the PEB framework, enabling an iterative form of modularity-informed causal inference. In this setting, empirically derived group-level partitions could act as soft priors that constrain or inform iterative subject and group level inference, effectively encouraging individual networks to conform to a shared modular organization while preserving individual differences. Developing such principled group-level extensions constitutes both a conceptually important and practically valuable direction for future research—particularly for constructing task- and population-specific effective connectivity atlases.

Finally, in clinical neuroscience, this approach may be of particular interest for investigating alterations in the brain’s hierarchical organization in neurological and psychiatric disorders. While most existing work has focused on abnormalities in the hierarchies of *functional* and *structural* networks, the present method enables principled multiscale parcellation of (potentially very large) *effective* networks, allowing direct interrogation of the hierarchical organization of causal interactions in both healthy and pathological brains. This framework makes it possible to examine, for example, differences in scaling behavior between control and patient populations, alterations in the spatial configuration of parcels across groups, abnormal fragmentation or excessive integration of causal networks in specific disorders, and the particular scales at which such abnormalities emerge. More broadly, by providing access to the brain’s causal architecture across scales, the proposed multiscale parcellation scheme offers a new avenue for studying how multiscale organization and near-critical dynamics jointly support the balance between integration and segregation in brain networks—and how disruptions of this multiscale balance may underlie dysfunction in disease.

## Conclusions

This technical note introduced a principled multiscale top-down parcellation scheme for dynamic causal models, with application to large-scale neuroimaging data. The proposed method generalizes the Δ*BIC* approach by formulating parcellation explicitly as a Bayesian model comparison problem. Because the partitioned model is nested within the full model, Δ*F* could be computed efficiently using Bayesian model reduction, thereby circumventing the need to fit separate submodels and enabling scalability to whole-brain causal networks.

At the theoretical level, transforming the original directed graph (into precision-weighted squared connections, adjusted for posterior change in uncertainty) revealed an intuitive relationship under the naïve independence assumption: changes in free energy map onto the negative graph cut, Δ*F* = −*Cut*. This result establishes a direct analytical link between Bayesian model evidence and the minCut problem in graph theory, allowing causal network partitioning to be solved efficiently using spectral clustering. The statistical significance of candidate bi-partitions was further assessed by comparing the observed minCut to its empirical null distribution obtained from matched randomized graphs.

The method was validated using both simulated and empirical causal models, establishing face and construct validity. Applied to empirical data, the resulting multiscale partitions revealed scale-dependent regularities and alluded to scale-invariant behavior across several dynamical measures. In addition, we examined whether the resulting partitions were conditionally independent given their boundary states. To this end, we identified Markov blanket states of cyclic causal models operationally as boundary states of the recurrent effective connectome, and introduced a hypothesis testing framework to assess evidence for additional Markov blanket states inferred from functional connectivity. In the present dataset and model, such additional states were largely absent, supporting the conditional independence of the identified partitions.

Finally, we discussed several natural extensions of the proposed framework, including multi-partitioning, group-level inference with random effects, and parametric alternatives to permutation-based significance testing, as well as potential applications in systems and clinical neuroscience. By providing a scalable and statistically grounded approach to uncovering hierarchical structure in causal brain networks, this work offers a foundation for future investigations of multiscale organization, as well as integration–segregation trade-offs that emerge near criticality, and their potential disruption in brain disorders.

## Software and Data availability

MATLAB code to reproduce the results presented in this paper is available at: https://github.com/tszarghami/Parcellation. Linearized DCM, PEB, and BMR procedures are implemented using MATLAB routines (spm_dcm_J.m, spm_dcm_peb.m, spm_dcm_bmr.m) in SPM12 software package (https://www.fil.ion.ucl.ac.uk/spm/). Spectral clustering is implemented using the Compressive Spectral Clustering Toolbox (http://cscbox.gforge.inria.fr/), and graph randomization is performed using functions from the Brain Connectivity Toolbox (https://sites.google.com/site/bctnet/). The attention-to-visual-motion fMRI dataset analyzed in this study is publicly available at: https://www.fil.ion.ucl.ac.uk/spm/data/attention/. Appendices are provided as Supplementary Material.

## Author Contributions

TSZ: Conceptualization, Methodology, Analysis, Interpretation, Writing.

## Acknowledgements

An early version of this work was presented online to colleagues from the Methods Group at the UCL Wellcome Center for Human Neuroimaging, in November 2020. I appreciate the feedback and the discussions generated during and after that session. I also sincerely thank the three anonymous reviewers for their careful, thoughtful, and constructive comments, which substantially strengthened the clarity and rigor of this work.

## Footnotes

1 Criticality is a dynamical state at the edge of a phase transition, characterized by scale-free activity, long-range correlations and optimal information processing; see (Hengen & Shew, 2025) for a recent review and perspective.

2 Conventionally, (biological) systems are modeled either bottom-up or top-down. Bottom-up models explain how elementary causes lead to ensemble-level functions, whereas top-down models start from ensemble descriptions and try to infer the underlying components (D’Angelo & Jirsa, 2022).

3 Hence, the following terms are used generically and interchangeably in this paper, without implying the specialized meanings they may carry in specific domains: segment, partition, module, subset, cluster, community, parcel.

4 Minimizing within-group dispersion is usually equivalent to maximizing between-group separation, for example based on the law of total variance in k-means clustering.

5 This is conceptually similar to piecewise regression.

6 A Bayes factor is the ratio of the evidences of two competing statistical models and is used to quantify the support for one model over the other (Kass & Raftery, 1995).

7 A dynamic causal model usually includes latent variables (*ϕ*) other than the causal connections (*θ*)—e.g. hemodynamic variables—in which case the partitioned model takes the form: *P*(*θ, ϕ*|*H*_*a*_) = *P*(*θ*_1_, *ϕ*_1_) *P*(*θ*_2_, *ϕ*_2_). For brevity, the *ϕ* variables are not shown explicitly in this paper; they can be assumed to be grouped with the *θ* variables.

8 If Δ*F* is positive, this indicates that the removed connections were most likely redundant in the original causal model, suggesting that the structure-learning stage of model development may have been inadequate. In SPM12, a DCM can be structurally optimized post hoc (using greedy search and Bayesian model reduction) via the routine *spm_dcm_bmr_all*.*m*. Bayesian model reduction is discussed further below.

9 Note that a distinct but related line of work models *interactions* between the presumed modules in a spatially distributed dynamical system—such as the brain—as neuronal messages containing mean-fields passed between the modules (Parr et al., 2020). A pre-requisite for modeling such between-module interactions is identifying the effective modules in the first place, for example using the parcellation scheme presented here.

10 The idea of computing a p-value associated with Bayes factor (BF) dates back to Good (Good, 1957) as a symbol of “Bayes/non-Bayes compromise” (Good, 1992). While empirical null distributions of BF are typically constructed via permutations or simulations (Feldl et al., 2025; Modrák et al., 2025; Schad et al., 2024; Schönbrodt & Wagenmakers, 2018; Sekulovski et al., 2024; Servin & Stephens, 2007; Xu & Guan, 2014), the theoretical null distribution of BF—and the associated p-value—can be derived analytically for specific models, such as linear regression (Zhou & Guan, 2018). Here, we rely on empirical null distributions but discuss potential analytical approaches in the Discussion.

11 Note the resemblance with Eq. (2): −Δ*BIC* > *C*_*α*_.

12 BMR has been successfully applied to both linear and nonlinear DCMs (Almgren et al., 2018; Friston et al., 2016; Jafarian et al., 2019, 2021, 2024, 2025; Papadopoulou et al., 2017; Pinotsis et al., 2019; Razi et al., 2017; Sokolov et al., 2019; Van de Steen et al., 2019; Zarghami, 2023; Zarghami et al., 2023). Notably, BMR has been shown to provide more robust posteriors for nonlinear models than fitting models to the data separately, and to be less susceptible to violations of the Laplace (i.e. Gaussian) assumption and local maxima problems (Friston et al., 2016). However, parcellating high-dimensional nonlinear DCMs—such as those used to model EEG/MEG/LFP data (David et al., 2006; Jafarian et al., 2025; Kiebel et al., 2008; Medrano et al., 2024; Pereira et al., 2021)—may be more challenging than for linear DCMs if the assumptions of BMR are strongly violated and affect the free energy approximation. While this primarily impacts absolute evidence values, the present results rely on relative comparisons across partitions, which are expected to be more robust. Quantifying the reliability of BMR in such high-dimensional nonlinear settings remains an important topic for future work.

13 In the context of DCM, (Frässle et al., 2017, 2021) also adopt naïve MFA to estimate large-scale effective brain networks. This assumption has two consequences: First, it ignores the fact that redundant paths between two nodes tend to introduce correlations between parameters that reflect linear symmetries in the model (e.g., increasing one connection having the same effect as decreasing another), as articulated by the Reviewer. Second, adopting a factorized variational posterior leads to underestimation of the (approximate) model evidence, as implied by the Bogolyubov inequality (Bogolubov Jr, 1966; Parr et al., 2020). These consequences for model accuracy and evidence are typically accepted in exchange for reduced model complexity and more efficient inference. In practice, sensitivity analyses comparing naïve and full BMR could be used to assess the impact of this assumption more rigorously.

14 Typically, posterior variances of the connections are several orders of magnitude smaller than the prior variance, meaning that estimates become more *precise* after seeing the data; that is, 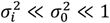, which implies 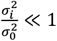 and 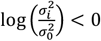.

15 The unnormalized graph Laplacian is computed as *L* = *D* − *A*, where *D* is a diagonal matrix of node degrees and *A* is the adjacency matrix.

16 Generalized versions of spectral clustering have been developed for directed graphs (Meilă & Pentney, 2007; Rohe et al., 2016; Sevi et al., 2022; Zhou et al., 2005). These methods are particularly useful for capturing directional flow in highly asymmetric graphs with low reciprocity. The current empirical DCM exhibits reciprocal connections by design. In cases where a highly nonreciprocal causal network is to be parcellated, it would be worthwhile to compare classical and generalized spectral clustering approaches—for example by comparing their induced cut sizes in terms of log Bayes factors (since |*Cut*| = |Δ*F*| = |*F*(*H*_*a*_) − *F*(*H*_0_)| → Δ*Cut* = *F*(*H*_*a*_) − *F*(*H*^′^_*a*_) = log *BF*).

17 We used a modified version of the spectral clustering MATLAB routine in the Compressive Spectral Clustering Toolbox (Tremblay et al., 2016): http://cscbox.gforge.inria.fr/

18 Free energy, 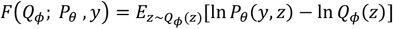, is normally a functional of the variational posterior (*Q*_*ϕ*_) and a function of the fixed generative model (*P*_*θ*_), parametrized by *θ*. However, when generating randomized networks, *θ* turns into random variable(s), following which *F* and Δ*F*^*max*^ (as functions of *θ*) also become random variables. This provides the rationale for defining a null distribution over Δ*F*^*max*^. In practice, this null distribution is approximated using the empirical distribution of minCut, as described below.

19 Bottlenecks are the small number of links that connect different modules in the topology of a modular graph (Tavara & Schliep, 2021).

20 The first-scale parent-free hypothesis is treated as a family by itself.

21 A simpler way of controlling the false discovery rate (FDR) at each scale would be to apply the direct FDR procedure to all hypotheses at that scale, regardless of family membership. In our analyses, the family-based approach of (Benjamini & Bogomolov, 2014) exhibited higher statistical power than the direct approach.

22 We thank the Reviewer for suggesting to report the modularity effect size.

23 Using *randmio_dir_connected*.*m* routine in Brain Connectivity Toolbox: http://www.brain-connectivity-toolbox.net/

24 In practice, a conservative and unbiased empirical p-value (for a limited number of permutations) was computed (Feldl et al., 2025; Phipson & Smyth, 2016): 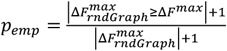. The addition of 1 to both numerator and denominator prevents the p-value from being exactly zero when none of the 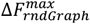 values exceed the observed Δ*F*^*max*^.

25 FMRI timeseries were acquired from a normal subject at 2 Tesla using a gradient echo-planar sequence (TE = 40ms; TR = 3.22 seconds; matrix size = 64×64×32, voxel size 3×3×3mm). Four consecutive 100-scan sessions were acquired, comprising a sequence of 10-scan blocks under five conditions: *Dummy, Fixation, Attention, No attention*, and *Stationary*. In the *Attention* condition, the subject was asked to detect changes in the radial velocity of moving dots. In *No attention*, the subject was asked to simply look at the moving dots. In the *Stationary* condition, the subject viewed stationary dots. The order of the conditions alternated between *Fixation* and photic stimulation. The subject fixated at the center of the screen in all conditions. No overt response was required in any condition and there were no actual speed changes in the moving dots. For more details see (Büchel & Friston, 1997; Friston et al., 2021).

26 The approximate Gaussianity of both Δ*F*_*rndCut*_ and 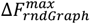 is not accidental. The statistical physics of graph partitioning allows interpolation between minimum cuts and random cuts, as elaborated in (Schreiber & Martin, 1999).

27 At first glance, this partition may appear divisible—but it is not. Only the spatial extent of the *nodes* in the causal graph is shown here; *edges* are omitted to avoid visual clutter. The two seemingly disjoint bilateral regions are, in fact, strongly causally connected, due to the assumption in (Friston et al., 2021) that homologous regions are spatially proximal when optimizing the range of local effective connections.

28 In a Bayesian network, the Markov blanket of a random variable is the minimal set of variables that renders it conditionally independent of all others. It comprises the parents, children, and co-parents of the children of the variable.

29 The stretched beta prior has a single hyperparameter α governing the concentration of probability mass around zero. When *α* = 1, the distribution is uniform; when *α* > 1, it is peaked around 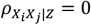; and when 0 < *α* < 1, the distribution concentrates toward the extremes (−1, +1), assigning lower probability to values near zero (Kucharský et al., 2023). See Fig. 14-A.

30 A Bayes factor is information-consistent if it diverges to infinity—thus falsifying the null hypothesis—when presented with overwhelmingly informative data (Jeffreys, 1961; Kucharský et al., 2023).

31 The number of candidate bi-partitions is given by the Stirling number of the second kind, *S*(*n*, 2) = 2^*n*−1^ − 1.

32 Even for small random networks, the minCut distribution shows Gaussian-like peakedness (see Fig. 5-D).

33 Some of the results in (Schreiber & Martin, 1999) extend to *weighted* random graphs. For example, although the “self-averaging” property of the minCut distribution has been primarily discussed for unweighted graphs,, it is typical of processes involving many contributing terms, as discussed in (Schreiber & Martin, 1999) and demonstrated numerically in Fig. S4.

34 In Supplementary Fig. S4, average minCut values are plotted against the number of nodes N for weighted random graphs matched to the transformed empirical graphs in degree and weight distributions across scales. The mean and variance of the minCut scale approximately linearly with N, consistent with theoretical and numerical results in (Schreiber & Martin, 1999). Fig. S4-C further demonstrates the self-averaging property and Gaussian-like shape of the distributions.

35 To generalize theoretical results from unweighted to weighted graphs, one may map weighted graphs onto unweighted multigraphs, which allow multiple parallel edges between vertex pairs (Newman, 2004).

36 For comparison, parameters of a transformed Tracy–Widom distribution have been estimated numerically to define an asymptotic null distribution for *modularity* in Gaussian random graphs (i.e., with Gaussian distributed edge weights) (Chang et al., 2014).

## References

Aguilera, M., Poc-López, Á., Heins, C., & Buckley, C. L. (2022). Knitting a Markov blanket is hard when you are out-of-equilibrium: Two examples in canonical nonequilibrium models (arXiv:2207.12914). arXiv. 10.48550/arXiv.2207.12914

Alexander-Bloch, A. F., Gogtay, N., Meunier, D., Birn, R., Clasen, L., Lalonde, F., Lenroot, R., Giedd, J., & Bullmore, E. T. (2010). Disrupted modularity and local connectivity of brain functional networks in childhood-onset schizophrenia. Frontiers in Systems Neuroscience, 4, 147.

Almgren, H., Van de Steen, F., Kühn, S., Razi, A., Friston, K., & Marinazzo, D. (2018). Variability and reliability of effective connectivity within the core default mode network: A multi-site longitudinal spectral DCM study. Neuroimage, 183, 757–768.

Ans, B., Hérault, J., Jutten, C., & others. (1985). Architectures neuromimétiques adaptatives: Détection de primitives. Proceedings of Cognitiva, 85, 593–597.

Ao, P. (2004). Potential in stochastic differential equations: Novel construction. Journal of Physics A: Mathematical and General, 37(3), L25.

Ashourvan, A., Telesford, Q. K., Verstynen, T., Vettel, J. M., & Bassett, D. S. (2019). Multi-scale detection of hierarchical community architecture in structural and functional brain networks. PLOS ONE, 14(5), e0215520. 10.1371/journal.pone.0215520

Axer, M., & Amunts, K. (2022). Scale matters: The nested human connectome. Science, 378(6619), 500–504. 10.1126/science.abq2599

Baba, K., Shibata, R., & Sibuya, M. (2004). Partial correlation and conditional correlation as measures of conditional independence. Australian & New Zealand Journal of Statistics, 46(4), 657–664.

Baba, K., & Sibuya, M. (2005). Equivalence of partial and conditional correlation coefficients. Journal of the Japan Statistical Society, 35(1), 1–19.

Bartels, R. H., & Stewart, G. W. (1972). Algorithm 432 [C2]: Solution of the matrix equation AX+ XB= C [F4]. Communications of the ACM, 15(9), 820–826.

Bastiani, M., & Roebroeck, A. (2015). Unraveling the multiscale structural organization and connectivity of the human brain: The role of diffusion MRI. Frontiers in Neuroanatomy, 9. 10.3389/fnana.2015.00077

Bausch, J. (2013). On the efficient calculation of a linear combination of chi-square random variables with an application in counting string vacua. Journal of Physics A: Mathematical and Theoretical, 46(50), 505202.

Beggs, J. M., & Plenz, D. (2003). Neuronal avalanches in neocortical circuits. Journal of Neuroscience, 23(35), 11167–11177.

Bell, A. J., & Sejnowski, T. J. (1995). An information-maximization approach to blind separation and blind deconvolution. Neural Computation, 7(6), 1129–1159.

Benjamini, Y., & Bogomolov, M. (2014). Selective inference on multiple families of hypotheses. Journal of the Royal Statistical Society Series B: Statistical Methodology, 76(1), 297–318.

Benozzo, D., Baggio, G., Baron, G., Chiuso, A., Zampieri, S., & Bertoldo, A. (2024). Analyzing asymmetry in brain hierarchies with a linear state-space model of resting-state fMRI data. Network Neuroscience, 8(3), 965–988. 10.1162/netn_a_00381

Bernhardt, B. C., Smallwood, J., Keilholz, S., & Margulies, D. S. (2022). Gradients in brain organization. In NeuroImage (Vol. 251, p. 118987). Elsevier.

Betzel, R. F., & Bassett, D. S. (2017). Multi-scale brain networks. NeuroImage, 160, 73–83. 10.1016/j.neuroimage.2016.11.006

Bhattacharya, A., Linero, A., & Oates, C. J. (2024). Grand Challenges in Bayesian Computation (arXiv:2410.00496). arXiv. 10.48550/arXiv.2410.00496

Bogolubov Jr, N. N. (1966). On model dynamical systems in statistical mechanics. Physica, 32(5), 933–944.

Bonifazi, P., Erramuzpe, A., Diez, I., Gabilondo, I., Boisgontier, M. P., Pauwels, L., Stramaglia, S., Swinnen, S. P., & Cortes, J. M. (2018). Structure–function multi-scale connectomics reveals a major role of the fronto-striato-thalamic circuit in brain aging. Human Brain Mapping, 39(12), 4663–4677. 10.1002/hbm.24312

Boonstra, T. W., He, B. J., & Daffertshofer, A. (2013). Scale-free dynamics and critical phenomena in cortical activity. In Frontiers in physiology (Vol. 4, p. 79). Frontiers Media SA.

Botta-Dukát, Z. (2018). Cautionary note on calculating standardized effect size (SES) in randomization test. Community Ecology, 19(1), 77–83.

Büchel, C., & Friston, K. J. (1997). Modulation of connectivity in visual pathways by attention: Cortical interactions evaluated with structural equation modelling and fMRI. Cerebral Cortex (New York, NY: 1991), 7(8), 768–778.

Bui, T. N. (1983). On Bisecting Random Graphs.

Chang, Y.-T., Pantazis, D., & Leahy, R. M. (2014). To cut or not to cut? Assessing the modular structure of brain networks. NeuroImage, 91, 99–108. 10.1016/j.neuroimage.2014.01.010

Chen, J., & Gupta, A. K. (1997). Testing and Locating Variance Changepoints with Application to Stock Prices. Journal of the American Statistical Association, 92(438), 739–747. 10.1080/01621459.1997.10474026

Cheng, S.-S., Wang, H.-M., & Fu, H.-C. (2008). BIC-based audio segmentation by divide-and-conquer. 2008 IEEE International Conference on Acoustics, Speech and Signal Processing, 4841–4844.

Chialvo, D. R. (2010). Emergent complex neural dynamics. Nature Physics, 6(10), 744–750.

Chung-Hsien Wu, Yu-Hsien Chiu, Chi-Jiun Shia, & Chun-Yu Lin. (2006). Automatic segmentation and identification of mixed-language speech using delta-BIC and LSA-based GMMs. IEEE Transactions on Audio, Speech and Language Processing, 14(1), 266–276. 10.1109/TSA.2005.852992

Clauset, A., Shalizi, C. R., & Newman, M. E. (2009). Power-law distributions in empirical data. SIAM Review, 51(4), 661–703.

Cocchi, L., Gollo, L. L., Zalesky, A., & Breakspear, M. (2017). Criticality in the brain: A synthesis of neurobiology, models and cognition. Progress in Neurobiology, 158, 132–152.

Cohen, J. (1988). Statistical power analysis for the behavioral sciences. routledge.

Costa, M., Gonçalves, A. M., & Teixeira, L. (2016). Change-point detection in environmental time series based on the informational approach. Electronic Journal of Applied Statistical Analysis, 9(2).

D’Acunto, G., Bonchi, F., Morales, G. D. F., & Petri, G. (2024). Extracting the Multiscale Causal Backbone of Brain Dynamics. In F. Locatello & V. Didelez (Eds.), Proceedings of the Third Conference on Causal Learning and Reasoning (Vol. 236, pp. 265–295). PMLR. https://proceedings.mlr.press/v236/d-textsc-char13acunto24a.html

D’Angelo, E., & Jirsa, V. (2022). The quest for multiscale brain modeling. Trends in Neurosciences, 45(10), 777–790. 10.1016/j.tins.2022.06.007

Dauwels, J. (2007). On Variational Message Passing on Factor Graphs. 2007 IEEE International Symposium on Information Theory, 2546–2550. 10.1109/ISIT.2007.4557602

David, O., Kiebel, S. J., Harrison, L. M., Mattout, J., Kilner, J. M., & Friston, K. J. (2006). Dynamic causal modeling of evoked responses in EEG and MEG. NeuroImage, 30(4), 1255–1272.

De Clerck, B., Van Utterbeeck, F., & Rocha, L. E. (2024). Comparative analysis of graph randomization: Tools, methods, pitfalls, and best practices. arXiv Preprint arXiv:2405.05400.

Deco, G., McIntosh, A. R., Shen, K., Hutchison, R. M., Menon, R. S., Everling, S., Hagmann, P., & Jirsa, V. K. (2014). Identification of optimal structural connectivity using functional connectivity and neural modeling. Journal of Neuroscience, 34(23), 7910–7916.

Deco, G., Ponce-Alvarez, A., Mantini, D., Romani, G. L., Hagmann, P., & Corbetta, M. (2013). Resting-state functional connectivity emerges from structurally and dynamically shaped slow linear fluctuations. Journal of Neuroscience, 33(27), 11239–11252.

Dempster, A. P., Laird, N. M., & Rubin, D. B. (1977). Maximum likelihood from incomplete data via the EM algorithm. Journal of the Royal Statistical Society: Series B (Methodological), 39(1), 1–22.

Doucet, G., Naveau, M., Petit, L., Delcroix, N., Zago, L., Crivello, F., Jobard, G., Tzourio-Mazoyer, N., Mazoyer, B., Mellet, E., & Joliot, M. (2011). Brain activity at rest: A multiscale hierarchical functional organization. Journal of Neurophysiology, 105(6), 2753–2763. 10.1152/jn.00895.2010

Ester, M., Kriegel, H.-P., Sander, J., & Xu, X. (1996). Simoudis, Evangelos; Han, Jiawei; Fayyad, Usama M., eds. A Density-Based Algorithm for Discovering Clusters in Large Spatial Databases with Noise, 226–231.

Farahibozorg, S. R., Harrison, S. J., Bijsterbosch, J. D., Woolrich, M. W., & Smith, S. M. (2024). Multiscale Modes of Functional Brain Connectivity. Neuroscience. 10.1101/2024.05.28.596120

Feldl, M., Olayo-Alarcon, R., Amstalden, M. K., Zannoni, A., Peschel, S., Sharma, C. M., Brochado, R., & Müller, C. L. (2025). Statistical end-to-end analysis of large-scale microbial growth data with DGrowthR. bioRxiv, 2025–03.

Ferrarini, L., Veer, I. M., Baerends, E., Van Tol, M., Renken, R. J., Van Der Wee, N. J. A., Veltman, Dirk. J., Aleman, A., Zitman, F. G., Penninx, B. W. J. H., Van Buchem, M. A., Reiber, J. H. C., Rombouts, S. A. R. B., & Milles, J. (2009). Hierarchical functional modularity in the resting-state human brain. Human Brain Mapping, 30(7), 2220–2231. 10.1002/hbm.20663

Frässle, S., Lomakina, E. I., Kasper, L., Manjaly, Z. M., Leff, A., Pruessmann, K. P., Buhmann, J. M., & Stephan, K. E. (2018). A generative model of whole-brain effective connectivity. NeuroImage, 179, 505–529. 10.1016/j.neuroimage.2018.05.058

Frässle, S., Lomakina, E. I., Razi, A., Friston, K. J., Buhmann, J. M., & Stephan, K. E. (2017). Regression DCM for fMRI. NeuroImage, 155, 406–421. 10.1016/j.neuroimage.2017.02.090

Frässle, S., Manjaly, Z. M., Do, C. T., Kasper, L., Pruessmann, K. P., & Stephan, K. E. (2021). Whole-brain estimates of directed connectivity for human connectomics. NeuroImage, 225, 117491. 10.1016/j.neuroimage.2020.117491

Friston, K. J. (2011). Functional and effective connectivity: A review. Brain Connectivity, 1(1), 13–36.

Friston, K. J., Fagerholm, E. D., Zarghami, T. S., Parr, T., Hipólito, I., Magrou, L., & Razi, A. (2021). Parcels and particles: Markov blankets in the brain. Network Neuroscience, 5(1), 211–251. 10.1162/netn_a_00175

Friston, K. J., Flandin, G., & Razi, A. (2022). Dynamic causal modelling of COVID-19 and its mitigations. Scientific Reports, 12(1), 12419. 10.1038/s41598-022-16799-8

Friston, K. J., Glaser, D. E., Henson, R. N., Kiebel, S., Phillips, C., & Ashburner, J. (2002). Classical and Bayesian inference in neuroimaging: Applications. Neuroimage, 16(2), 484–512.

Friston, K. J., Litvak, V., Oswal, A., Razi, A., Stephan, K. E., Van Wijk, B. C. M., Ziegler, G., & Zeidman, P. (2016). Bayesian model reduction and empirical Bayes for group (DCM) studies. NeuroImage, 128, 413–431. 10.1016/j.neuroimage.2015.11.015

Friston, K. J., & Penny, W. (2003). Posterior probability maps and SPMs. Neuroimage, 19(3), 1240–1249.

Friston, K. J., Penny, W., Phillips, C., Kiebel, S., Hinton, G., & Ashburner, J. (2002). Classical and Bayesian inference in neuroimaging: Theory. NeuroImage, 16(2), 465–483.

Friston, K., Mattout, J., Trujillo-Barreto, N., Ashburner, J., & Penny, W. (2007). Variational free energy and the Laplace approximation. NeuroImage, 34(1), 220–234. 10.1016/j.neuroimage.2006.08.035

Friston, K., Parr, T., & Zeidman, P. (2019). Bayesian model reduction (arXiv:1805.07092). arXiv. 10.48550/arXiv.1805.07092

Friston, K., & Penny, W. (2011). Post hoc Bayesian model selection. NeuroImage, 56(4), 2089–2099. 10.1016/j.neuroimage.2011.03.062

Fu, Y., & Anderson, P. W. (1986). Application of statistical mechanics to NP-complete problems in combinatorial optimisation. Journal of Physics A: Mathematical and General, 19(9), 1605.

Gao, Z., Xiao, X., Fang, Y.-P., Rao, J., & Mo, H. (2024). A Selective Review on Information Criteria in Multiple Change Point Detection. Entropy, 26(1), 50. 10.3390/e26010050

Geiger, D., & Meek, C. (2005). Structured variational inference procedures and their realizations. International Workshop on Artificial Intelligence and Statistics, 104–111.

Good, I. J. (1957). Saddle-point methods for the multinomial distribution. The Annals of Mathematical Statistics, 28(4), 861–881.

Good, I. J. (1992). The Bayes/non-Bayes compromise: A brief review. Journal of the American Statistical Association, 87(419), 597–606.

Greaves, M. D., Novelli, L., & Razi, A. (2025). Structurally informed resting-state effective connectivity recapitulates cortical hierarchy. bioRxiv, 2024–04.

Haak, K. V., Marquand, A. F., & Beckmann, C. F. (2018). Connectopic mapping with resting-state fMRI. Neuroimage, 170, 83–94.

Haimovici, A., Tagliazucchi, E., Balenzuela, P., & Chialvo, D. R. (2013). Brain Organization into Resting State Networks Emerges at Criticality<? Format?> on a Model of the Human Connectome. Physical Review Letters, 110(17), 178101.

Han, Z., Yang, X., Zhao, X., Jiguet, F., Tryjanowski, P., & Wang, H. (2023). Mongolian Lark as an indicator of taxonomic, functional and phylogenetic diversity of steppe birds. Avian Research, 14, 100124.

Hansen, J. Y., Shafiei, G., Voigt, K., Liang, E. X., Cox, S. M. L., Leyton, M., Jamadar, S. D., & Misic, B.(2023). Integrating multimodal and multiscale connectivity blueprints of the human cerebral cortex in health and disease. PLOS Biology, 21(9), e3002314. 10.1371/journal.pbio.3002314

He, Y., Wang, J., Wang, L., Chen, Z. J., Yan, C., Yang, H., Tang, H., Zhu, C., Gong, Q., Zang, Y., & others. (2009). Uncovering intrinsic modular organization of spontaneous brain activity in humans. PloS One, 4(4), e5226.

He, Yirong, Zeng, D., Li, Q., Chu, L., Dong, X., Liang, X., Sun, L., Liao, X., Zhao, T., Chen, X., Lei, T., Men, W., Wang, Y., Wang, D., Hu, M., Pan, Z., Zhang, H., Liu, N., Tan, S., … Li, S. (2025). The multiscale brain structural re-organization that occurs from childhood to adolescence correlates with cortical morphology maturation and functional specialization. PLOS Biology, 23(4), e3002710. 10.1371/journal.pbio.3002710

He, Z., Zhang, T., Wang, Q., Zhang, S., Cao, G., Liu, T., Zhao, S., Jiang, X., Guo, L., Yuan, Y., & others. (2024). Brain functional gradients are related to cortical folding gradient. Cerebral Cortex, 34(11), bhae453.

Hengen, K. B., & Shew, W. L. (2025). Is criticality a unified setpoint of brain function? Neuron, 113(16), 2582–2598.

Henzinger, M., Li, J., Rao, S., & Wang, D. (2024). Deterministic near-linear time minimum cut in weighted graphs. Proceedings of the 2024 Annual ACM-SIAM Symposium on Discrete Algorithms (SODA), 3089–3139.

Hervías-Parejo, S., Traveset, A., Nogales, M., Heleno, R., Llewelyn, J., & Strona, G. (2025). Sampling biases across interaction types affect the robustness of ecological multilayer networks. Ecological Informatics, 89, 103183.

Hesse, J., & Gross, T. (2014). Self-organized criticality as a fundamental property of neural systems. Frontiers in Systems Neuroscience, 8, 166.

Hochberg, Y., & Tamhane, A. C. (1987). Multiple comparison procedures. John Wiley & Sons, Inc.

Hu, Y., Nie, Y., Yang, H., Cheng, J., Fan, Y., & Di, Z. (2010). Measuring the significance of community structure in complex networks. Physical Review E—Statistical, Nonlinear, and Soft Matter Physics, 82(6), 066106.

Huntenburg, J. M., Bazin, P.-L., & Margulies, D. S. (2018). Large-scale gradients in human cortical organization. Trends in Cognitive Sciences, 22(1), 21–31.

Hyvärinen, A. (2013). Independent component analysis: Recent advances. Philosophical Transactions of the Royal Society A: Mathematical, Physical and Engineering Sciences, 371(1984), 20110534.

Iraji, A., Fu, Z., Faghiri, A., Duda, M., Chen, J., Rachakonda, S., DeRamus, T., Kochunov, P., Adhikari, B. M., Belger, A., Ford, J. M., Mathalon, D. H., Pearlson, G. D., Potkin, S. G., Preda, A., Turner, J. A., Van Erp, T. G. M., Bustillo, J. R., Yang, K., … Calhoun, V. D. (2023). Identifying canonical and replicable multi-scale intrinsic connectivity networks in 100k+ RESTING-STATE FMRI datasets. Human Brain Mapping, 44(17), 5729–5748. 10.1002/hbm.26472

Jaakkola, T. S., & Jordan, M. I. (1998). Improving the mean field approximation via the use of mixture distributions. In Learning in graphical models (pp. 163–173). Springer.

Jafarian, A., Assem, M. K., Kocagoncu, E., Lanskey, J. H., Fye, H., Williams, R., Quinn, A. J., Pitt, J., Raymont, V., Lowe, S., & others. (2025). Neurophysiological progression in Alzheimer’s disease: Insights from dynamic causal modelling of longitudinal magnetoencephalography. Human Brain Mapping, 46(8), e70234.

Jafarian, A., Assem, M. K., Kocagoncu, E., Lanskey, J. H., Williams, R., Cheng, Y.-J., Quinn, A. J., Pitt, J., Raymont, V., Lowe, S., & others. (2024). Reliability of dynamic causal modelling of resting-state magnetoencephalography. Human Brain Mapping, 45(10), e26782.

Jafarian, A., Zeidman, P., Litvak, V., & Friston, K. (2019). Structure learning in coupled dynamical systems and dynamic causal modelling. Philosophical Transactions of the Royal Society A: Mathematical, Physical and Engineering Sciences, 377(2160), 20190048. 10.1098/rsta.2019.0048

Jafarian, A., Zeidman, P., Wykes, R. C., Walker, M., & Friston, K. J. (2021). Adiabatic dynamic causal modelling. NeuroImage, 238, 118243.

Jeffreys, H. (1961). The theory of probability. OuP Oxford.

Jordan, M. I., Ghahramani, Z., Jaakkola, T. S., & Saul, L. K. (1998). An Introduction to Variational Methods for Graphical Models. In M. I. Jordan (Ed.), Learning in Graphical Models (pp. 105–161). Springer Netherlands. 10.1007/978-94-011-5014-9_5

Kanter, I., & Sompolinsky, H. (1987). Graph optimisation problems and the Potts glass. Journal of Physics A: Mathematical and General, 20(11), L673.

Karrer, B., Levina, E., & Newman, M. E. (2008). Robustness of community structure in networks. Physical Review E—Statistical, Nonlinear, and Soft Matter Physics, 77(4), 046119.

Kass, R. E., & Raftery, A. E. (1995). Bayes factors. Journal of the American Statistical Association, 90(430), 773–795.

Keysers, C., Gazzola, V., & Wagenmakers, E.-J. (2020). Using Bayes factor hypothesis testing in neuroscience to establish evidence of absence. Nature Neuroscience, 23(7), 788–799. 10.1038/s41593-020-0660-4

Khemakhem, I., Kingma, D., Monti, R., & Hyvarinen, A. (2020). Variational autoencoders and nonlinear ica: A unifying framework. International Conference on Artificial Intelligence and Statistics, 2207–2217.

Kiebel, S. J., Garrido, M. I., Moran, R. J., & Friston, K. J. (2008). Dynamic causal modelling for EEG and MEG. Cognitive Neurodynamics, 2(2), 121–136.

Kingma, D. P., & Welling, M. (2013). Auto-encoding variational bayes. arXiv Preprint arXiv:1312.6114.

Kucharský, Š., Wagenmakers, E.-J., Van Den Bergh, D., & Ly, A. (2023). Analytic Posterior Distribution and Bayes Factor for Pearson Partial Correlations. PsyArXiv. 10.31234/osf.io/6muwy

Lancichinetti, A., Radicchi, F., & Ramasco, J. J. (2010). Statistical significance of communities in networks. Physical Review E—Statistical, Nonlinear, and Soft Matter Physics, 81(4), 046110.

Latifi, S., & Carmichael, S. T. (2024). The emergence of multiscale connectomics-based approaches in stroke recovery. Trends in Neurosciences, 47(4), 303–318. 10.1016/j.tins.2024.01.003

Lawrance, A. (1976). On conditional and partial correlation. The American Statistician, 30(3), 146–149.

Li, S., Zhang, J., Huang, K., & Gao, C. (2014). A graph partitioning approach for Bayesian Network structure learning. Proceedings of the 33rd Chinese Control Conference, 2887–2892. 10.1109/ChiCC.2014.6897098

Li, Y., & Qi, Y. (2020). Asymptotic distribution of modularity in networks. Metrika, 83(4), 467–484. 10.1007/s00184-019-00740-7

Linkenkaer-Hansen, K., Nikouline, V. V., Palva, J. M., & Ilmoniemi, R. J. (2001). Long-range temporal correlations and scaling behavior in human brain oscillations. Journal of Neuroscience, 21(4), 1370–1377.

Litvak, V., Jafarian, A., Zeidman, P., Tibon, R., Henson, R. N., & Friston, K. (2019). There’s no such thing as a ‘true’ model: The challenge of assessing face validity. 2019 IEEE International Conference on Systems, Man and Cybernetics (SMC), 4403–4408. 10.1109/SMC.2019.8914255

Lloyd, S. (1982). Least squares quantization in PCM. IEEE Transactions on Information Theory, 28(2), 129–137.

Lu, M., Guo, Z., Gao, Z., Cao, Y., & Fu, J. (2022). Multiscale Brain Network Models and Their Applications in Neuropsychiatric Diseases. Electronics, 11(21), 3468. 10.3390/electronics11213468

Maksimenko, V. A., Runnova, A. E., Frolov, N. S., Makarov, V. V., Nedaivozov, V., Koronovskii, A. A., Pisarchik, A., & Hramov, A. E. (2018). Multiscale neural connectivity during human sensory processing in the brain. Physical Review E, 97(5), 052405. 10.1103/PhysRevE.97.052405

Margulies, D. S., Ghosh, S. S., Goulas, A., Falkiewicz, M., Huntenburg, J. M., Langs, G., Bezgin, G., Eickhoff, S. B., Castellanos, F. X., Petrides, M., & others. (2016). Situating the defaultmode network along a principal gradient of macroscale cortical organization. Proceedings of the National Academy of Sciences, 113(44), 12574–12579.

Marković, D., Friston, K. J., & Kiebel, S. J. (2024). Bayesian sparsification for deep neural networks with Bayesian model reduction. IEEE Access, 12, 88231–88242. 10.1109/ACCESS.2024.3417219

Masharipov, R., Knyazeva, I., Nikolaev, Y., Korotkov, A., Didur, M., Cherednichenko, D., & Kireev, M. (2021). Providing evidence for the null hypothesis in functional magnetic resonance imaging using group-level bayesian inference. Frontiers in Neuroinformatics, 15, 738342.

Medrano, J., Friston, K. J., & Zeidman, P. (2024). Dynamic Causal Models of Time-Varying Connectivity. arXiv Preprint arXiv:2411.16582.

Meilă, M., & Pentney, W. (2007). Clustering by weighted cuts in directed graphs. Proceedings of the 2007 SIAM International Conference on Data Mining, 135–144.

Meinshausen, N. (2008). Hierarchical testing of variable importance. Biometrika, 95(2), 265–278.

Meng, X., Iraji, A., Fu, Z., Kochunov, P., Belger, A., Ford, J., McEwen, S., Mathalon, D. H., Mueller, A. A., Pearlson, G., & others. (2022). Multimodel order independent component analysis: A data-driven method for evaluating brain functional network connectivity within and between multiple spatial scales. Brain Connectivity, 12(7), 617–628.

Meunier, D. (2009). Hierarchical modularity in human brain functional networks. Frontiers in Neuroinformatics, 3. 10.3389/neuro.11.037.2009

Meunier, D., Lambiotte, R., & Bullmore, E. T. (2010). Modular and Hierarchically Modular Organization of Brain Networks. Frontiers in Neuroscience, 4. 10.3389/fnins.2010.00200

Mirshahvalad, A., Beauchesne, O. H., Archambault, E., & Rosvall, M. (2013). Resampling effects on significance analysis of network clustering and ranking. PloS One, 8(1), e53943.

Mirshahvalad, A., Lindholm, J., Derlén, M., & Rosvall, M. (2012). Significant communities in large sparse networks. PloS One, 7(3), e33721.

Mirzaeian, S., Jensen, K. M., Ballem, A. R., Calhoun, V. D., & Iraji, A. (2025). Ultra-High Order Independent Component Analysis for Intrinsic Connectivity Networks in Resting-State Functional Magnetic Resonance Imaging Data. 2025 IEEE 22nd International Symposium on Biomedical Imaging (ISBI), 1–4. 10.1109/ISBI60581.2025.10981312

Modrák, M., Stroppel, S., & Bürkner, P.-C. (2025). Simulation-based validation of Bayes factor computation. arXiv Preprint arXiv:2508.11814.

Moghimi, P., Dang, A. T., Do, Q., Netoff, T. I., Lim, K. O., & Atluri, G. (2022). Evaluation of functional MRI-based human brain parcellation: A review. Journal of Neurophysiology, 128(1), 197–217.

Morante, M., Frølich, K., & Rehman, N. ur. (2024). Multiscale Functional Connectivity: Exploring the brain functional connectivity at different timescales (arXiv:2406.19041). arXiv. 10.48550/arXiv.2406.19041

Morante, M., Frølich, K., Shahzaib, M., Shakil, S., & Rehman, N. U. (2024). Multiscale Functional Connectivity analysis of episodic memory reconstruction. Frontiers in Cognition, 3, 1433234. 10.3389/fcogn.2024.1433234

Morey, R. D., & Rouder, J. N. (2011). Bayes factor approaches for testing interval null hypotheses. Psychological Methods, 16(4), 406.

Munn, B. R., Müller, E. J., Favre-Bulle, I., Scott, E., Lizier, J. T., Breakspear, M., & Shine, J. M. (2024). Multiscale organization of neuronal activity unifies scale-dependent theories of brain function. Cell, 187(25), 7303-7313.e15. 10.1016/j.cell.2024.10.004

Nashed, J. Y., Sandarage, R., Pasarikovski, C. R., Gallivan, J. P., & Cook, D. J. (2025). From gradients to cognition: Linking cortical manifolds to brain flexibility and disorder. Frontiers in Cognition, 4, 1690469.

Newman, M. E. J. (2004). Analysis of weighted networks. Physical Review E, 70(5), 056131. 10.1103/PhysRevE.70.056131

Newman, M. E. J. (2006). Modularity and community structure in networks. Proceedings of the National Academy of Sciences, 103(23), 8577–8582. 10.1073/pnas.0601602103

Nichols, T., & Hayasaka, S. (2003). Controlling the familywise error rate in functional neuroimaging: A comparative review. Statistical Methods in Medical Research, 12(5), 419–446.

Oizumi, M., Tsuchiya, N., & Amari, S. (2016). Unified framework for information integration based on information geometry. Proceedings of the National Academy of Sciences, 113(51), 14817–14822.

Papadopoulou, M., Cooray, G., Rosch, R., Moran, R., Marinazzo, D., & Friston, K. (2017). Dynamic causal modelling of seizure activity in a rat model. Neuroimage, 146, 518–532.

Park, B., Paquola, C., Bethlehem, R. A. I., Benkarim, O., Neuroscience in Psychiatry Network (NSPN) Consortium, Mišić, B., Smallwood, J., Bullmore, E. T., Bernhardt, B. C., Bullmore, E., Dolan, R., Goodyer, I., Fonagy, P., Jones, P., Moutoussis, M., Hauser, T., Neufeld, S., Romero-Garcia, R., St Clair, M., … Chamberlain, S. (2022). Adolescent development of multiscale structural wiring and functional interactions in the human connectome. Proceedings of the National Academy of Sciences, 119(27), e2116673119. 10.1073/pnas.2116673119

Parr, T., Sajid, N., & Friston, K. J. (2020). Modules or Mean-Fields? Entropy, 22(5), 552. 10.3390/e22050552

Penny, W. D. (2012). Comparing Dynamic Causal Models using AIC, BIC and Free Energy. NeuroImage, 59(1), 319–330. 10.1016/j.neuroimage.2011.07.039

Penny, W. D., Stephan, K. E., Mechelli, A., & Friston, K. J. (2004). Comparing dynamic causal models. NeuroImage, 22(3), 1157–1172. 10.1016/j.neuroimage.2004.03.026

Pereira, I., Frässle, S., Heinzle, J., Schöbi, D., Do, C. T., Gruber, M., & Stephan, K. E. (2021). Conductance-based dynamic causal modeling: A mathematical review of its application to cross-power spectral densities. NeuroImage, 245, 118662.

Pérez-Ortega, S., Verdú, M., Garrido-Benavent, I., Rabasa, S., Green, T. A., Sancho, L. G., & de los Ríos, A. (2023). Invariant properties of mycobiont-photobiont networks in Antarctic lichens. Global Ecology and Biogeography, 32(11), 2033–2046.

Phipson, B., & Smyth, G. K. (2016). Permutation P-values should never be zero: Calculating exact P-values when permutations are randomly drawn. arXiv Preprint arXiv:1603.05766.

Pines, A. R., Larsen, B., Cui, Z., Sydnor, V. J., Bertolero, M. A., Adebimpe, A., Alexander-Bloch, A. F., Davatzikos, C., Fair, D. A., Gur, R. C., Gur, R. E., Li, H., Milham, M. P., Moore, T. M., Murtha, K., Parkes, L., Thompson-Schill, S. L., Shanmugan, S., Shinohara, R. T., … Satterthwaite, T. D. (2022). Dissociable multi-scale patterns of development in personalized brain networks. Nature Communications, 13(1), 2647. 10.1038/s41467-022-30244-4

Pinotsis, D. A., Loonis, R., Bastos, A. M., Miller, E. K., & Friston, K. J. (2019). Bayesian modelling of induced responses and neuronal rhythms. Brain Topography, 32(4), 569–582.

Prando, G., Zorzi, Mattia, Bertoldo, A., Corbetta, M., Zorzi, Marco, & Chiuso, A. (2020). Sparse DCM for whole-brain effective connectivity from resting-state fMRI data. NeuroImage, 208, 116367. 10.1016/j.neuroimage.2019.116367

Presigny, C., & Fallani, F. D. V. (2022). Multiscale modeling of brain network organization. Reviews of Modern Physics, 94(3), 031002. 10.1103/RevModPhys.94.031002

Qian, H., & Beard, D. A. (2005). Thermodynamics of stoichiometric biochemical networks in living systems far from equilibrium. Biophysical Chemistry, 114(2–3), 213–220.

Razi, A., Seghier, M. L., Zhou, Y., McColgan, P., Zeidman, P., Park, H.-J., Sporns, O., Rees, G., & Friston, K. J. (2017). Large-scale DCMs for resting-state fMRI. Network Neuroscience, 1(3), 222–241.

Reichardt, J., & Bornholdt, S. (2006). When are networks truly modular? Physica D: Nonlinear Phenomena, 224(1–2), 20–26.

Reichardt, J., & Bornholdt, S. (2007). Partitioning and modularity of graphs with arbitrary degree distribution. Physical Review E—Statistical, Nonlinear, and Soft Matter Physics, 76(1), 015102.

Rohe, K., Qin, T., & Yu, B. (2016). Co-clustering directed graphs to discover asymmetries and directional communities. Proceedings of the National Academy of Sciences, 113(45), 12679–12684.

Rousseau, P.-N., Bazin, P.-L., & Steele, C. J. (2025). Multiscale gradients of corticopontine structural connectivity. Scientific Reports, 15(1), 16399. 10.1038/s41598-025-00886-7

Royer, J., Paquola, C., Valk, S. L., Kirschner, M., Hong, S.-J., Park, B., Bethlehem, R. A., Leech, R., Yeo, B. T., Jefferies, E., & others. (2024). Gradients of brain organization: Smooth sailing from methods development to user community. Neuroinformatics, 22(4), 623–634.

Rubinov, M., Sporns, O., Thivierge, J.-P., & Breakspear, M. (2011). Neurobiologically realistic determinants of self-organized criticality in networks of spiking neurons. PLoS Computational Biology, 7(6), e1002038.

Sala, A., Caminiti, S. P., Presotto, L., Premi, E., Pilotto, A., Turrone, R., Cosseddu, M., Alberici, A., Paghera, B., Borroni, B., Padovani, A., & Perani, D. (2017). Altered brain metabolic connectivity at multiscale level in early Parkinson’s disease. Scientific Reports, 7(1), 4256. 10.1038/s41598-017-04102-z

Schad, D. J., Modrák, M., & Vasishth, S. (2024). How accurate are Bayes factor-based null hypothesis tests? A simulation study. arXiv Preprint arXiv:2406.08022.

Schirner, M., McIntosh, A. R., Jirsa, V., Deco, G., & Ritter, P. (2018). Inferring multi-scale neural mechanisms with brain network modelling. Elife, 7, e28927.

Schönbrodt, F. D., & Wagenmakers, E.-J. (2018). Bayes factor design analysis: Planning for compelling evidence. Psychonomic Bulletin & Review, 25(1), 128–142.

Schreiber, G. R., & Martin, O. C. (1999). Cut Size Statistics of Graph Bisection Heuristics. SIAM Journal on Optimization, 10(1), 231–251. 10.1137/S1052623497321523

Seifi, M., Junier, I., Rouquier, J.-B., Iskrov, S., & Guillaume, J.-L. (2013). Stable community cores in complex networks. In Complex Networks (pp. 87–98). Springer.

Sekulovski, N., Marsman, M., & Wagenmakers, E.-J. (2024). A Good check on the Bayes factor. Behavior Research Methods, 56(8), 8552–8566.

Servin, B., & Stephens, M. (2007). Imputation-based analysis of association studies: Candidate regions and quantitative traits. PLoS Genetics, 3(7), e114.

Sevi, H., Jonckheere, M., & Kalogeratos, A. (2022). Generalized spectral clustering for directed and undirected graphs. arXiv Preprint arXiv:2203.03221.

Shew, W. L., & Plenz, D. (2013). The functional benefits of criticality in the cortex. The Neuroscientist, 19(1), 88–100.

Shine, J. M., Müller, E. J., Munn, B., Cabral, J., Moran, R. J., & Breakspear, M. (2021). Computational models link cellular mechanisms of neuromodulation to large-scale neural dynamics. Nature Neuroscience, 24(6), 765–776. 10.1038/s41593-021-00824-6

Singh, A. K., & Phillips, S. (2010). Hierarchical control of false discovery rate for phase locking measures of EEG synchrony. NeuroImage, 50(1), 40–47.

Sokolov, A. A., Zeidman, P., Erb, M., Ryvlin, P., Pavlova, M. A., & Friston, K. J. (2019). Linking structural and effective brain connectivity: Structurally informed Parametric Empirical Bayes (si-PEB). Brain Structure and Function, 224(1), 205–217.

Stephan, K. E., Tittgemeyer, M., Knösche, T. R., Moran, R. J., & Friston, K. J. (2009). Tractography-based priors for dynamic causal models. Neuroimage, 47(4), 1628–1638.

Stumpf, M. P., & Porter, M. A. (2012). Critical truths about power laws. Science, 335(6069), 665–666.

Tatsumi, S., Cadotte, M. W., & Mori, A. S. (2019). Individual-based models of community assembly: Neighbourhood competition drives phylogenetic community structure. Journal of Ecology, 107(2), 735–746.

Tavara, S., & Schliep, A. (2021). Effects of network topology on the performance of consensus and distributed learning of SVMs using ADMM. PeerJ Computer Science, 7, e397. 10.7717/peerj-cs.397

Theodorou, T., Mporas, I., & Fakotakis, N. (2014). An Overview of Automatic Audio Segmentation. International Journal of Information Technology and Computer Science, 6(11), 1–9. 10.5815/ijitcs.2014.11.01

Tognoli, E., & Kelso, J. S. (2014). The metastable brain. Neuron, 81(1), 35–48.

Toker, D., & Sommer, F. T. (2019). Information integration in large brain networks. PLOS Computational Biology, 15(2), e1006807. 10.1371/journal.pcbi.1006807

Tremblay, N., Puy, G., Gribonval, R., & Vandergheynst, P. (2016). Compressive spectral clustering. International Conference on Machine Learning, 1002–1011.

Tritschler, A., & Gopinath, R. A. (1999). Improved speaker segmentation and segments clustering using the bayesian information criterion. 6th European Conference on Speech Communication and Technology, 679–682. 10.21437/Eurospeech.1999-174x

Van de Steen, F., Almgren, H., Razi, A., Friston, K., & Marinazzo, D. (2019). Dynamic causal modelling of fluctuating connectivity in resting-state EEG. Neuroimage, 189, 476–484.

Van de Ville, D., Britz, J., & Michel, C. M. (2010). EEG microstate sequences in healthy humans at rest reveal scale-free dynamics. Proceedings of the National Academy of Sciences, 107(42), 18179–18184.

Van Den Heuvel, M. P., Scholtens, L. H., & Kahn, R. S. (2019). Multiscale Neuroscience of Psychiatric Disorders. Biological Psychiatry, 86(7), 512–522. 10.1016/j.biopsych.2019.05.015

Vatsavai, R. R., Symons, C. T., Chandola, V., & Jun, G. (2011). GX-Means: A model-based divide and merge algorithm for geospatial image clustering. Procedia Computer Science, 4, 186–195. 10.1016/j.procs.2011.04.020

Veech, J. A. (2012). Significance testing in ecological null models. Theoretical Ecology, 5(4), 611–616.

Von Luxburg, U. (2007). A tutorial on spectral clustering. Statistics and Computing, 17(4), 395–416. 10.1007/s11222-007-9033-z

Vos de Wael, R., Benkarim, O., Paquola, C., Lariviere, S., Royer, J., Tavakol, S., Xu, T., Hong, S.-J., Langs, G., Valk, S., & others. (2020). BrainSpace: A toolbox for the analysis of macroscale gradients in neuroimaging and connectomics datasets. Communications Biology, 3(1), 103.

Wang, C., & Shanechi, M. M. (2019). Estimating Multiscale Direct Causality Graphs in Neural Spike-Field Networks. IEEE Transactions on Neural Systems and Rehabilitation Engineering, 27(5), 857–866. 10.1109/TNSRE.2019.2908156

Wang, Y., Huang, H., Feng, C., & Liu, Z. (2015). Community detection based on minimum-cut graph partitioning. Web-Age Information Management: 16th International Conference, WAIM 2015, Qingdao, China, June 8-10, 2015. Proceedings 16, 57–69.

Weatherburn, C. E. (1949). A first course mathematical statistics (Vol. 158). CUP Archive.

Wright, D., Igel, C., & Selvan, R. (2024). BMRS: Bayesian Model Reduction for Structured Pruning (arXiv:2406.01345). arXiv. 10.48550/arXiv.2406.01345

Wu, G. R., Chen, F., Kang, D., Zhang, X., D. Marinazzo, & Chen, H. (2011). Multiscale Causal Connectivity Analysis by Canonical Correlation: Theory and Application to Epileptic Brain. IEEE Transactions on Biomedical Engineering, 58(11), 3088–3096. 10.1109/TBME.2011.2162669

Xing, E. (2004). Graph partition strategies for generalized mean field inference. Proc Uncertainty in Artificial Intelligence 20 (UAI2004).

Xing, E. P., Jordan, M. I., & Russell, S. (2002). A generalized mean field algorithm for variational inference in exponential families. Proceedings of the Nineteenth Conference on Uncertainty in Artificial Intelligence, 583–591.

Xu, H., & Guan, Y. (2014). Detecting local haplotype sharing and haplotype association. Genetics, 197(3), 823–838.

Xue, H., Li, H., Gao, C., & Shi, Z. (2010). Computationally efficient audio segmentation through a multi-stage BIC approach. 2010 3rd International Congress on Image and Signal Processing, 3774–3777. 10.1109/CISP.2010.5646687

Yekutieli, D. (2008). Hierarchical false discovery rate–controlling methodology. Journal of the American Statistical Association, 103(481), 309–316.

Yekutieli, D., Reiner-Benaim, A., Benjamini, Y., Elmer, G. I., Kafkafi, N., Letwin, N. E., & Lee, N. H. (2006). Approaches to multiplicity issues in complex research in microarray analysis. Statistica Neerlandica, 60(4), 414–437.

Zarghami, T. S. (2023). A new causal centrality measure reveals the prominent role of subcortical structures in the causal architecture of the extended default mode network. Brain Structure and Function, 228(8), 1917–1941. 10.1007/s00429-023-02697-w

Zarghami, T. S., Zeidman, P., Razi, A., Bahrami, F., & Hossein-Zadeh, G.-A. (2023). Dysconnection and cognition in schizophrenia: A spectral dynamic causal modeling study. Human Brain Mapping, 44(7), 2873–2896.

Zeidman, P., Friston, K., & Parr, T. (2023). A primer on Variational Laplace (VL). NeuroImage, 279, 120310. 10.1016/j.neuroimage.2023.120310

Zeidman, P., Jafarian, A., Seghier, M. L., Litvak, V., Cagnan, H., Price, C. J., & Friston, K. J. (2019). A guide to group effective connectivity analysis, part 2: Second level analysis with PEB. NeuroImage, 200, 12–25. 10.1016/j.neuroimage.2019.06.032

Zhao, Q., Hautamaki, V., & Fränti, P. (2008). Knee Point Detection in BIC for Detecting the Number of Clusters. In J. Blanc-Talon, S. Bourennane, W. Philips, D. Popescu, & P. Scheunders (Eds.), Advanced Concepts for Intelligent Vision Systems (Vol. 5259, pp. 664–673). Springer Berlin Heidelberg. 10.1007/978-3-540-88458-3_60

Zhou, D., Huang, J., & Schölkopf, B. (2005). Learning from labeled and unlabeled data on a directed graph. Proceedings of the 22nd International Conference on Machine Learning, 1036–1043.

Zhou, Q., & Guan, Y. (2018). On the null distribution of bayes factors in linear regression. Journal of the American Statistical Association, 113(523), 1362–1371.

Zhou, Z., Li, H., Srinivasan, D., Abdulkadir, A., Nasrallah, I. M., Wen, J., Doshi, J., Erus, G., Mamourian, E., Bryan, N. R., Wolk, D. A., Beason-Held, L., Resnick, S. M., Satterthwaite, T. D., Davatzikos, C., Shou, H., & Fan, Y. (2023). Multiscale functional connectivity patterns of the aging brain learned from harmonized rsfMRI data of the multi-cohort iSTAGING study. NeuroImage, 269, 119911. 10.1016/j.neuroimage.2023.119911

Zimmern, V. (2020). Why brain criticality is clinically relevant: A scoping review. Frontiers in Neural Circuits, 14, 54.

Zou, T., Chen, C., Chen, H., Wang, X., Gan, L., Wang, C., Gao, Q., Zhang, C., Liao, W., Cheng, J., & Li, R. (2024). Structural-functional connectivity decoupling in multiscale brain networks in Parkinson’s disease. BMC Neuroscience, 25(1), 78. 10.1186/s12868-024-00918-4

