## Supplementary Material for "Multiscale Parcellation of Dynamic Causal Models of the Brain"

Tahereh S. Zarghami[1*]

[1] Bio-Electric Department, School of Electrical and Computer Engineering, University of Tehran, Iran.

#### Supplementary Figures

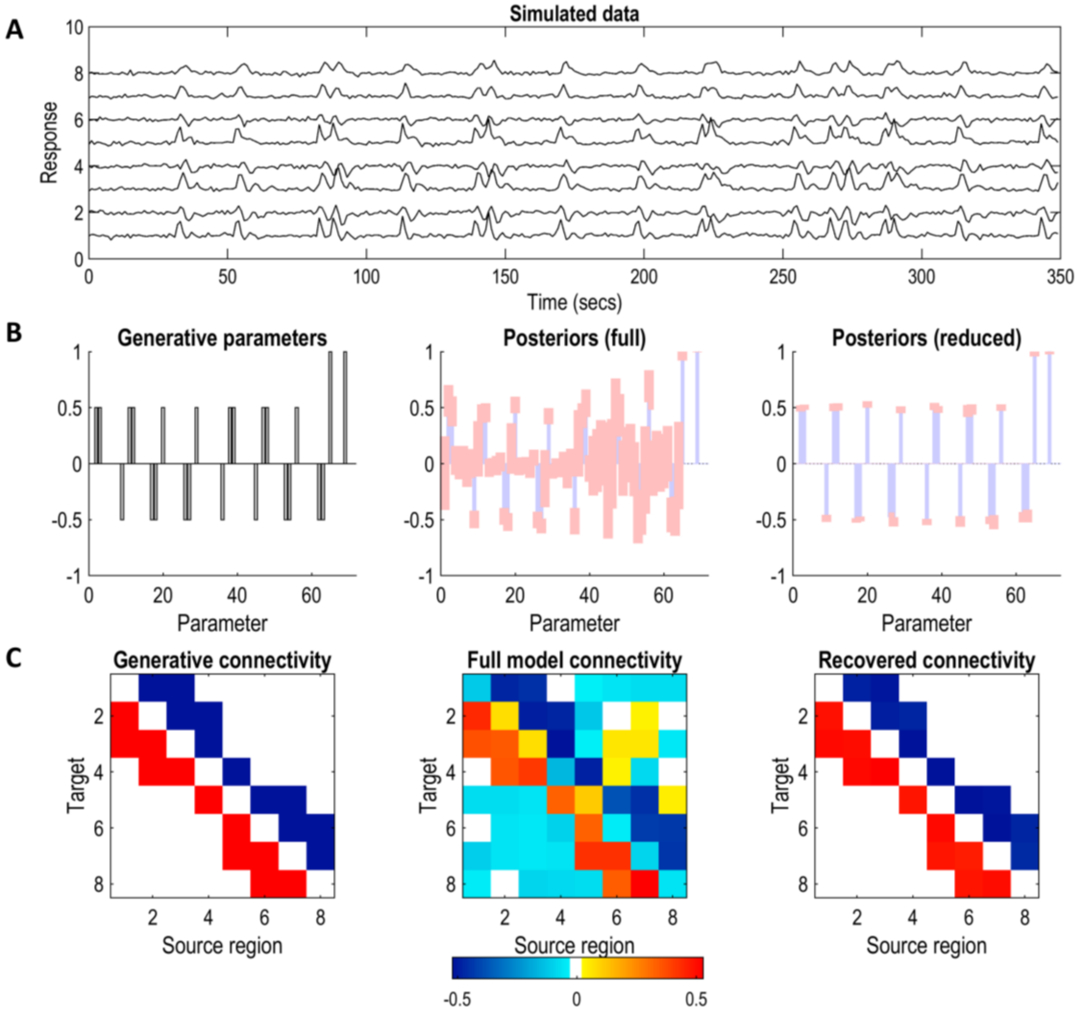


Fig. S1: Synthetic data generation and DCM model inversion results, for the causal network in Fig. 3-B. For the details, please refer to the caption of Fig. 4.

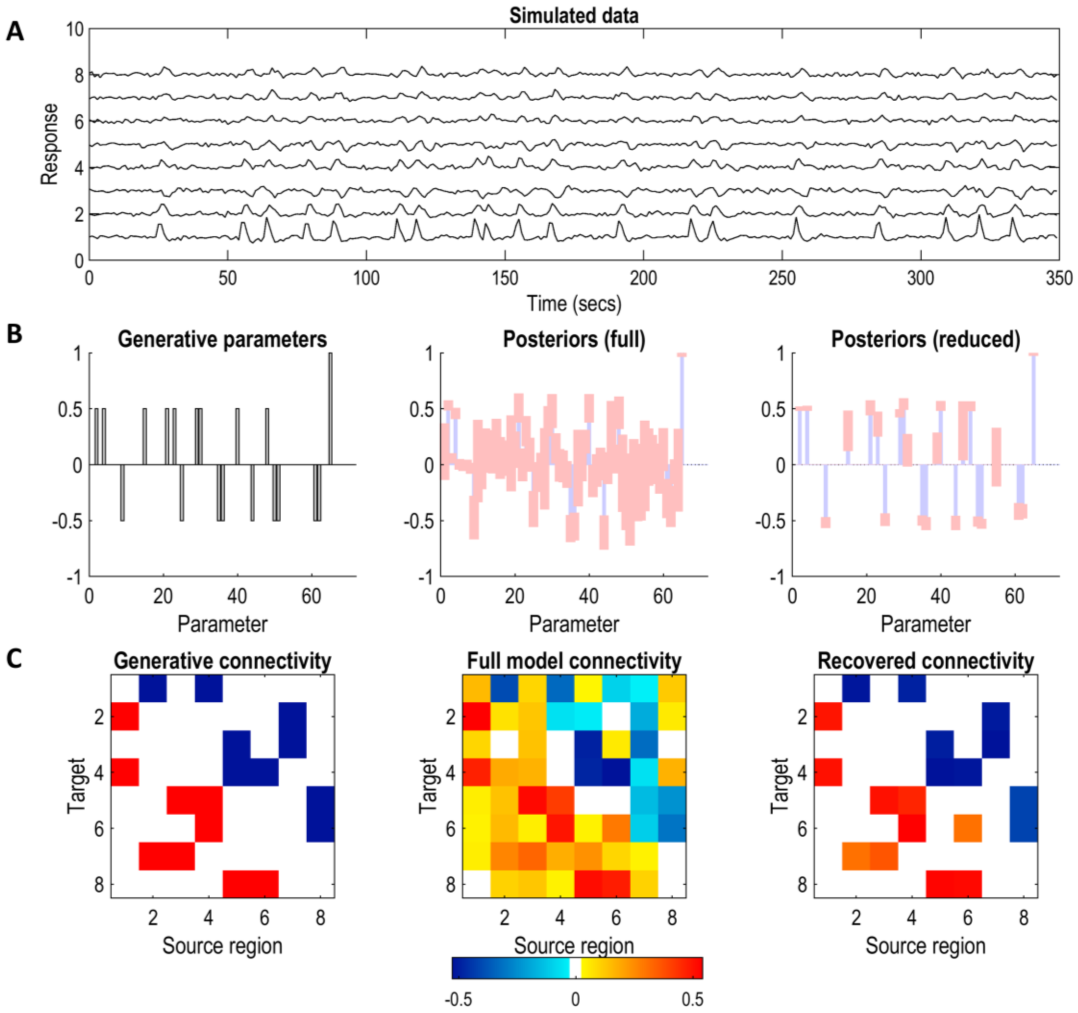


Fig. S2: Synthetic data generation and DCM model inversion results, for the causal network in Fig. 3-C. For the details, please refer to the caption of Fig. 4.

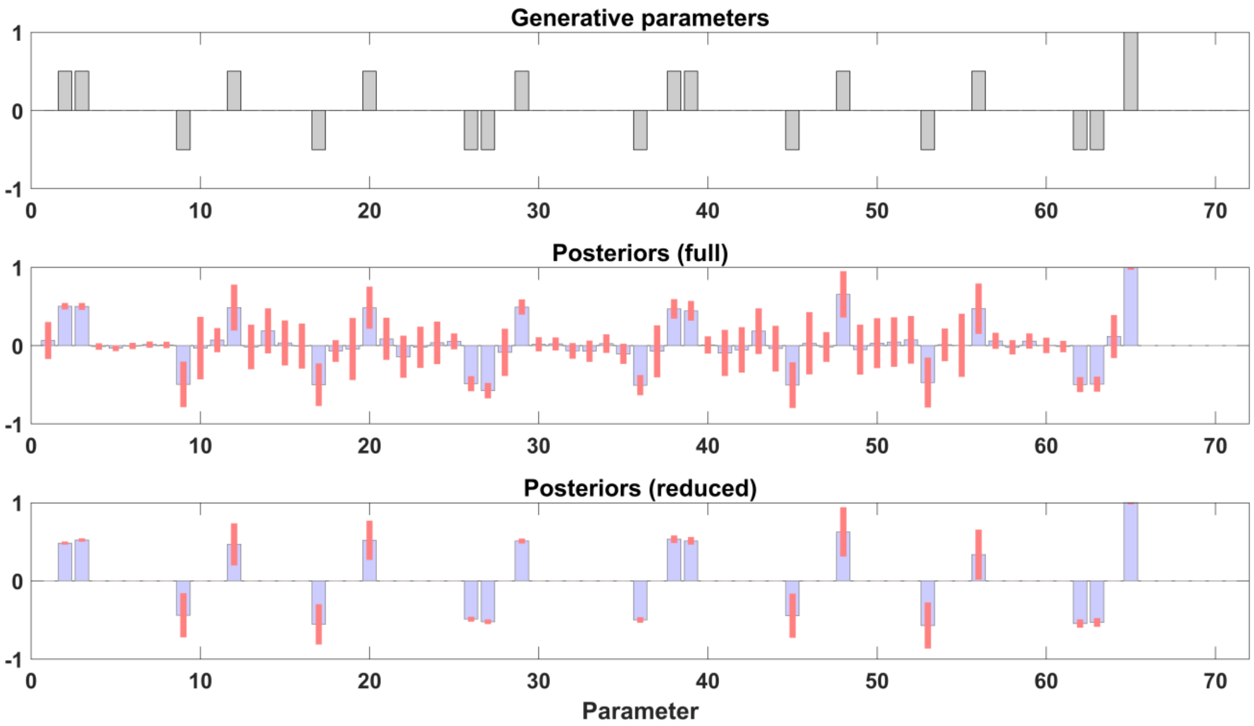


Fig. S3: An unpacked version of the bar plots in Fig. 4-B. (Top) Generative model parameters (i.e. ground truth connections). (Middle): Posterior parameter strengths, from fitting a fully-connected DCM to the synthetic timeseries. Blue bars represent expected values and pink error bars represent 95% credible intervals. (Bottom): Posterior parameter strengths following Bayesian model reduction. Weak and imprecise parameters have been pruned automatically, using exploratory BMR (*spm_dcm_bmr_all.m*), to improve model evidence.

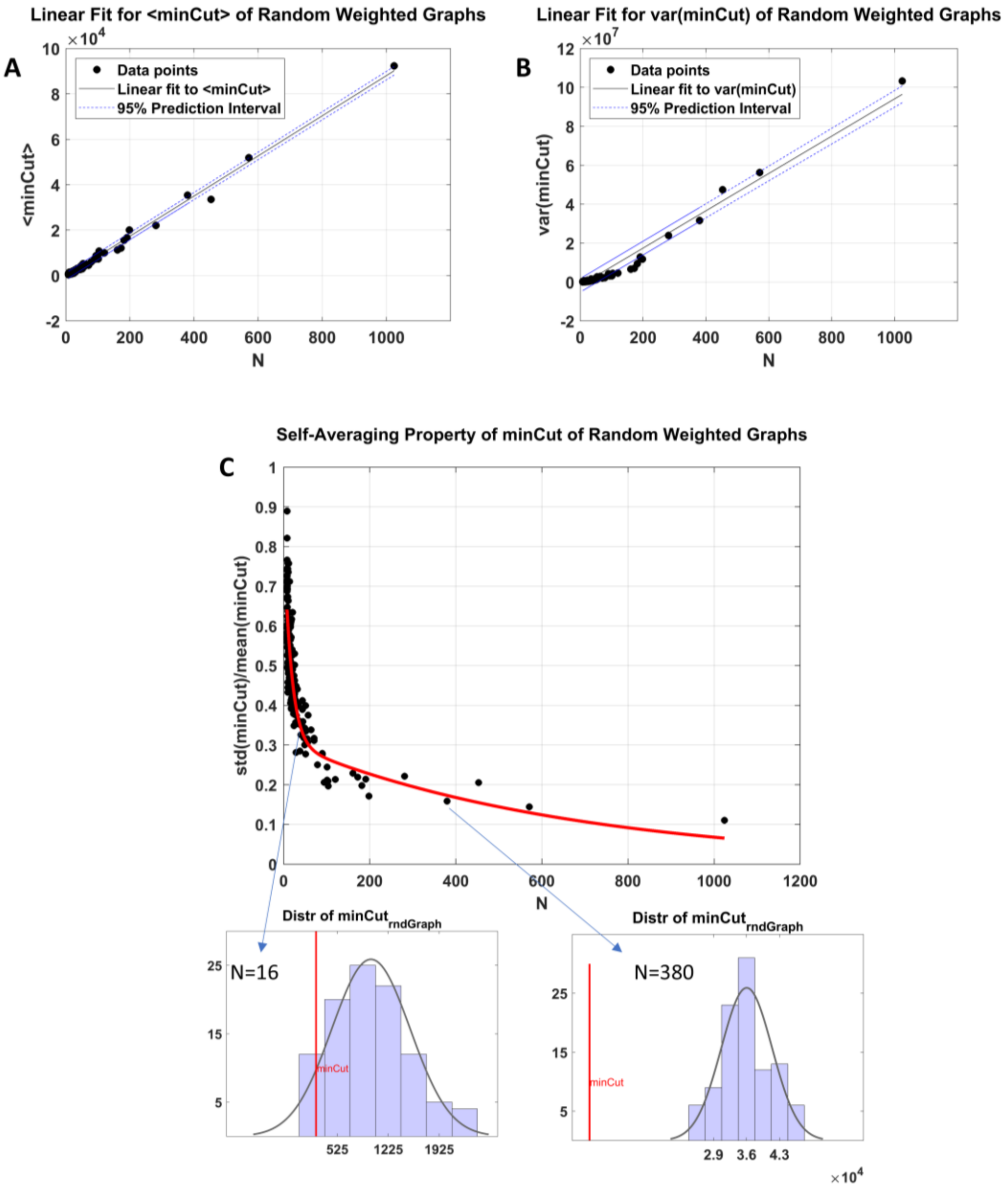


Fig. S4: Statistical properties of the minCut distribution in random weighted graphs. (A) Mean minCut as a function of the number of nodes $N$, exhibiting an approximately linear scaling, in agreement with theoretical predictions for random graphs (Schreiber & Martin, 1999). (B) Variance of the minCut as a function of $N$, which also increases approximately linearly. (C) Self-averaging behavior of the minCut in random weighted graphs. The ratio of the standard deviation to the mean minCut is plotted against $N$, for randomized graphs matched to transformed empirical networks. This ratio decreases rapidly with increasing $N$, approaching zero from above, suggesting that the results of (Schreiber & Martin, 1999) extend to *weighted* random graphs. Insets show representative minCut distributions for two randomized networks ($N = 16$ and $N = 380$). As network size increases, relative fluctuations of the minCut around its mean diminish, and eventually tend towards zero as $N \to \infty$, a property known as "self-averaging" that is typical of processes to which many terms contribute (Schreiber & Martin, 1999).

### Appendices

#### Appendix A: Proof for Eq. (4): $F(H_a) = F_1 + F_2$

In the free energy expression,

$$F = E_Q[\log P(y|\theta)] - D_{KL}[Q(\theta)||P(\theta)] \quad \text{(A1)}$$

substitute the factorized form of the prior, approximate posterior and data likelihood of the partitioned model (i.e. the alternative hypothesis, $H_a$):

$$P(\theta|H_a) = P(\theta_1)P(\theta_2) \quad \text{(A2)}$$

$$P(\theta|y, H_a) \approx Q(\theta|H_a) = Q(\theta_1)Q(\theta_2) \quad \text{(A3)}$$

$$P(y|\theta, H_a) = P(y_1, y_2|\theta_1, \theta_2) = P(y_1|\theta_1)P(y_2|\theta_2) \quad \text{(A4)}$$

which yields:

$$F(H_a) = \iint Q(\theta_1)Q(\theta_2)\log[P(y_1|\theta_1)P(y_2|\theta_2)]\,d\theta_1 d\theta_2 - \iint Q(\theta_1)Q(\theta_2)\log\frac{Q(\theta_1)Q(\theta_2)}{P(\theta_1)P(\theta_2)}\,d\theta_1 d\theta_2$$

$$F(H_a) = \iint Q(\theta_1)Q(\theta_2)\log P(y_1|\theta_1)\,d\theta_1 d\theta_2 + \iint Q(\theta_1)Q(\theta_2)\log P(y_2|\theta_2)\,d\theta_1 d\theta_2 - \iint Q(\theta_1)Q(\theta_2)\log\frac{Q(\theta_1)}{P(\theta_1)}\,d\theta_1 d\theta_2 - \iint Q(\theta_1)Q(\theta_2)\log\frac{Q(\theta_2)}{P(\theta_2)}\,d\theta_1 d\theta_2$$

$$F(H_a) = \int Q(\theta_1)\log P(y_1|\theta_1)\,d\theta_1 \int Q(\theta_2)d\theta_2 + \int Q(\theta_2)\log P(y_2|\theta_2)\,d\theta_2 \int Q(\theta_1)d\theta_1 - \int Q(\theta_1)\log\frac{Q(\theta_1)}{P(\theta_1)}\,d\theta_1 \int Q(\theta_2)d\theta_2 - \int Q(\theta_2)\log\frac{Q(\theta_2)}{P(\theta_2)}d\theta_2 \int Q(\theta_1)\,d\theta_1$$

$$F(H_a) = E_{Q(\theta_1)}[log\,P(y_1|\theta_1)] + E_{Q(\theta_2)}[log\,P(y_2|\theta_2)] - D_{KL}[Q(\theta_1)||P(\theta_1)] - D_{KL}[Q(\theta_2)||P(\theta_2)] \quad \text{(A5)}$$

$$\rightarrow F(H_a) = F_1 + F_2 \quad \text{(A6)}$$

#### Appendix B: Bayesian Model Reduction (BMR)

Bayesian model reduction refers to the analytic inversion of a reduced model using the prior and posterior of a full model, together with the prior of the reduced model. Reduced models are nested within a full model; that is, they include only a subset of the parameters of the full model after "switching off" the other parameters—by imposing very precise null priors, which shrinks them to zero. Consider Bayes' rule written for the full and reduced models, denoted by $m$ and $m^R$, respectively (Friston et al., 2019, 2016; Friston & Penny, 2011) :

$$P(\theta|y,m) = \frac{P(y|\theta,m)P(\theta|m)}{P(y|m)} \tag{B1}$$

$$P(\theta|y,m^R) = \frac{P(y|\theta,m^R)P(\theta|m^R)}{P(y|m^R)} \tag{B2}$$

Since the models differ only in terms of their priors, the likelihood terms are identical:

$$P(y|\theta,m^R) = P(y|\theta,m) \tag{B3}$$

Hence, equating Eqs. B1 and B2 via the likelihood yields:

$$\frac{P(\theta|y,m^R)P(y|m^R)}{P(\theta|m^R)} = P(y|\theta,m) = \frac{P(\theta|y,m)P(y|m)}{P(\theta|m)} \tag{B4}$$

By re-arranging Eq. B4, we obtain the posterior distribution over the parameters of the reduced model:

$$P(\theta|y,m^R) = P(\theta|y,m)\frac{P(\theta|m^R)}{P(\theta|m)}\frac{P(y|m)}{P(y|m^R)} \tag{B5}$$

On the right-hand side of Eq. B5, only $P(y|m^R)$ is unknown, which is the model evidence for the reduced model. This term can be obtained by integrating both sides of Eq. B5 over $\theta$:

$$\int P(\theta|y,m^R)\,d\theta = 1 = \int P(\theta|y,m)\frac{P(\theta|m^R)}{P(\theta|m)}\frac{P(y|m)}{P(y|m^R)}\,d\theta \tag{B6}$$

$$\rightarrow \frac{P(y|m^R)}{P(y|m)} = \int P(\theta|y,m)\frac{P(\theta|m^R)}{P(\theta|m)}\,d\theta \tag{B7}$$

$$\rightarrow \log P(y|m^R) = \log P(y|m) + \log\int P(\theta|y,m)\frac{P(\theta|m^R)}{P(\theta|m)}\,d\theta \tag{B8}$$

$$\rightarrow F^R = F + \log\int Q(\theta|y,m)\frac{P(\theta|m^R)}{P(\theta|m)}\,d\theta \tag{B9}$$

In the last line, optimized variational density and free energy have replaced the exact posterior density and log model evidence, respectively: $Q(\theta|Y,m) \approx P(\theta|Y,m)$ and $F \approx \log P(Y|m)$. Under Gaussian assumptions for the distributions in variational Laplace, the reduced posterior and free energy take simple forms, as elaborated in (Friston et al. 2016; Friston et al. 2018; Friston and Penny 2011). In short:

- Prior of the full model: $\mathcal{N}(\eta_F, \Sigma_F = \Pi_F^{-1})$
- Posterior of the full model: $\mathcal{N}(\mu_F, C_F = P_F^{-1})$
- Prior of the reduced model: $\mathcal{N}(\eta_R, \Sigma_R = \Pi_R^{-1})$
- Posterior of the reduced model (to be computed): $\mathcal{N}(\mu_R = ?, C_R = P_R^{-1} = ?)$

$$P_R = P_F + \Pi_R - \Pi_F \tag{B10}$$

$$\mu_R = C_R(P_F\mu_F + \Pi_R\eta_R - \Pi_F\eta_F) \tag{B11}$$

$$\Delta F = F_R - F_F = \frac{1}{2} Ln\, |\Pi_R P_F C_R \Sigma_F| - \frac{1}{2}(\mu_F^T P_F \mu_F + \eta_R^T \Pi_R \eta_R - \eta_F^T \Pi_F \eta_F - \mu_R^T P_R \mu_R) \tag{B12}$$

#### Appendix C: Naïve BMR

Here we introduce a naïve version of BMR, which assumes statistical independence among causal connections. This assumption is realized by imposing a diagonal structure on the covariance and precision matrices of the normally distributed connection strengths. In what follows, we derive the posterior of the reduced model and the reduced free energy under such naïve assumptions. As shown below, a key implication of this formulation for the graph partitioning problem is the elimination of matrix operations, which considerably reduces the computational burden and enhances scalability.

- Prior distribution of the full model: $\mathcal{N}(\eta_F = 0, \Sigma_F = \Pi_F^{-1} = \sigma_0^2 I)$
- Posterior distribution of the full model: $\mathcal{N}(\mu_F, C_F = P_F^{-1})$

  The posterior covariance and precision matrices of the full model (denoted by $C_F$and $P_F$, respectively) are diagonal by design. $\sigma_i^2$ denotes the posterior variance of connection $i \in \{1, \dots, N = n^2\}$, where $n =$ number of nodes/regions.

$$C_F = \begin{bmatrix} \sigma_1^2 & & 0 \\ & \ddots & \\ 0 & & \sigma_N^2 \end{bmatrix}, P_F = \begin{bmatrix} \sigma_1^{-2} & & 0 \\ & \ddots & \\ 0 & & \sigma_N^{-2} \end{bmatrix}$$

- Prior distribution of the reduced model: $\mathcal{N}\left(\eta_R = 0, \Sigma_R = \Pi_R^{-1} = \begin{bmatrix} \sigma_0^2 I & 0 \\ 0 & \varepsilon^2 I \end{bmatrix}\right)$

  $\rightarrow \Pi_R = \begin{bmatrix} \sigma_0^{-2} I & 0 \\ 0 & \varepsilon^{-2} I \end{bmatrix}$

  Since $(\varepsilon^2 \to 0^+) \Rightarrow (\varepsilon^{-2} \to +\infty)$, we shall represent $\varepsilon^2$ by $0^+$ and $\varepsilon^{-2}$ by $+\infty$, for brevity. In practice, typical values for $\varepsilon^2$ are on the order of $e^{-16}$, in the BMR functions of SPM12.

- Posterior distribution of the reduced model (to be computed):

  $\mathcal{N}(\mu_R = ?, C_R = P_R^{-1} = ?)$

We now substitute the above priors and posteriors in the BMR formulae from Appendix B, to derive the reduced posterior and free energy, under the Laplace approximation. The following points apply throughout the subsequent derivations:

- $\{C\}$ denotes the set of connections to 'cut out' and $\setminus$ stands for exclusion; hence, $\mu_{F\{C\}}$ is the vector holding the mean values of the removed (between-partition) connections and $\mu_{F\setminus\{C\}}$ contains the means of the retained (within-partition) connections. Without loss of generality, we assume that the connections have been ordered such that $\mu_F = \begin{bmatrix} \mu_{F\setminus\{C\}} \\ \mu_{F\{C\}} \end{bmatrix}$, and that the same order is preserved in the corresponding covariance and precision matrices. The cardinality of $\{C\}$ is represented by $N_C$.

- 0 denotes a matrix or vector of all zeros.
- Multiplication of diagonal matrices is commutative, i.e. $A_{diag}B_{diag} = B_{diag}A_{diag}$.
- The following relationships will be used:

$$\Sigma_R \mu_F = \begin{bmatrix} \sigma_0^2 I & 0 \\ 0 & 0^+ \end{bmatrix} \begin{bmatrix} \mu_{F\setminus\{C\}} \\ \mu_{F\{C\}} \end{bmatrix} \approx \begin{bmatrix} \sigma_0^2 \mu_{F\setminus\{C\}} \\ 0 \end{bmatrix} \quad \text{(C1)}$$

$$\Pi_R \begin{bmatrix} \mu_{F\setminus\{C\}} \\ 0 \end{bmatrix} = \begin{bmatrix} \sigma_0^{-2} I & 0 \\ 0 & \infty \end{bmatrix} \begin{bmatrix} \mu_{F\setminus\{C\}} \\ 0 \end{bmatrix} = \begin{bmatrix} \sigma_0^{-2} \mu_{F\setminus\{C\}} \\ 0 \end{bmatrix} \quad \text{(C2)}$$

We now proceed to derive the posterior of the reduced model, $\mathcal{N}(\mu_R = ?, C_R = P_R^{-1} = ?)$, and the reduced free energy, $\Delta F$, based on Eqs. B10-B12 from Appendix B:

**I)** $$P_R = P_F + \Pi_R - \Pi_F = P_F + \begin{bmatrix} \sigma_0^{-2} I & 0 \\ 0 & \infty \end{bmatrix} - \begin{bmatrix} \sigma_0^{-2} I & 0 \\ 0 & \sigma_0^{-2} I \end{bmatrix}$$

$$= P_F + \begin{bmatrix} 0 & 0 \\ 0 & \infty \end{bmatrix} = \begin{bmatrix} P_{F\setminus\{C\}} & 0 \\ 0 & \infty \end{bmatrix} \approx P_F \sigma_0^2 \Pi_R \quad \text{(C3)}$$

$$\rightarrow C_R = P_R^{-1} = \begin{bmatrix} C_{F\setminus\{C\}} & 0 \\ 0 & 0^+ \end{bmatrix} \approx C_F \sigma_0^{-2} \Sigma_R \quad \text{(C4)}$$

**II)** $$\mu_R = C_R(P_F\mu_F + \Pi_R\eta_R - \Pi_F\eta_F) = C_F\sigma_0^{-2}\Sigma_R\, (P_F\mu_F + \Pi_R * 0 - \Pi_F * 0)$$

$$= \sigma_0^{-2}\Sigma_R\mu_F \approx \begin{bmatrix} \mu_{F\setminus\{C\}} \\ 0 \end{bmatrix} \qquad \text{(using Eq. C1)} \quad \text{(C5)}$$

**III)** $$\Delta F = F_R - F_F = \frac{1}{2}\log\frac{|\Sigma_F|}{|C_F|}\frac{|C_R|^\dagger}{|\Sigma_R|^\dagger} - \frac{1}{2}(\mu_F^T P_F \mu_F + \eta_R^T \Pi_R \eta_R - \eta_F^T \Pi_F \eta_F - \mu_R^T P_R \mu_R) \quad \text{(C6)}$$

where $|\ \ |$ and $|\ \ |^\dagger$ denote the determinant and pseudo-determinant of the enclosed matrices, respectively. We will simplify the log-ratio term in Eq. C6:

$$\frac{1}{2}\log\frac{|\Sigma_F|}{|C_F|}\frac{|C_R|^\dagger}{|\Sigma_R|^\dagger} = \frac{1}{2}\log\left(\frac{\prod_{i=1}^N \sigma_0^2}{\prod_{i=1}^N \sigma_i^2} \times \frac{\prod_{i\in\setminus\{C\}} \sigma_i^2}{\prod_{i\in\setminus\{C\}} \sigma_0^2}\right) = \frac{1}{2}\log\left(\frac{\prod_{i\in\{C\}} \sigma_0^2}{\prod_{i\in\{C\}} \sigma_i^2}\right) = \frac{1}{2}\sum_{i\in\{C\}}\log\left(\frac{\sigma_0^2}{\sigma_i^2}\right) \quad \text{(C7)}$$

Substitute Eqs. C7 and C3 in Eq. C6:

$$\Delta F = \frac{1}{2} \sum_{i\in\{C\}} \log\left(\frac{\sigma_0^2}{\sigma_i^2}\right) - \frac{1}{2}\left(\mu_F^T P_F \mu_F - \mu_R^T (P_F \sigma_0^2\, \Pi_R)\mu_R\right) \tag{C8}$$

Using Eqs. C2 and C5, we get:

$$\Pi_R \mu_R = \sigma_0^{-2} \mu_R \tag{C9}$$

Substitute Eq. C9 in Eq. C8:

$$\Delta F = \frac{1}{2} \sum_{i\in\{C\}} \log\left(\frac{\sigma_0^2}{\sigma_i^2}\right) - \frac{1}{2}\left(\mu_F^T P_F \mu_F - \mu_R^T P_F \mu_R\right) \tag{C10}$$

Since $\mu_F = \begin{bmatrix}\mu_{F\backslash\{C\}}\\ 0\end{bmatrix} + \begin{bmatrix}0\\ \mu_{F\{C\}}\end{bmatrix}$, $P_F = \begin{bmatrix}P_{F\backslash\{C\}} & 0\\ 0 & P_{F\{C\}}\end{bmatrix}$ and $\mu_R \approx \begin{bmatrix}\mu_{F\backslash\{C\}}\\ 0\end{bmatrix}$, the last term in Eq. C10 can be written as: $\mu_R^T P_F \mu_R = \mu_{F\backslash\{C\}}^T P_{F\backslash\{C\}} \mu_{F\backslash\{C\}} = \mu_F^T P_F \mu_F - \mu_{F\{C\}}^T P_{F\{C\}} \mu_{F\{C\}}$.

Therefore:

$$\Delta F = \frac{1}{2} \sum_{i\in\{C\}} \log\left(\frac{\sigma_0^2}{\sigma_i^2}\right) - \frac{1}{2}\left\{\mu_F^T P_F \mu_F - \left(\mu_F^T P_F \mu_F - \mu_{F\{C\}}^T P_{F\{C\}} \mu_{F\{C\}}\right)\right\}$$

$$\Delta F = \frac{1}{2} \sum_{i\in\{C\}} \log\left(\frac{\sigma_0^2}{\sigma_i^2}\right) - \frac{1}{2}\left(\mu_{F\{C\}}^T P_{F\{C\}} \mu_{F\{C\}}\right)$$

$$\Delta F = -\frac{1}{2} \sum_{i\in\{C\}} \left(\sigma_i^{-2} \mu_F^2(i) + \log\left(\frac{\sigma_i^2}{\sigma_0^2}\right)\right) \tag{C11}$$

The final equality follows from the diagonality of the precision matrix. Notably, this illustrates how large matrix multiplications can be avoided under the naïve mean-field formulation.

Based on Eq. C11, in order to maximize $\Delta F$, the graph should be partitioned to minimize the sum of between-partition[1] connections squared and weighted by their precisions, plus the logarithm of the ratio of their posterior to prior variances. That is:

$$\max_{\{C\}} \Delta F \Leftrightarrow \min_{\{C\}} \sum_{i\in\{C\}} \left(\sigma_i^{-2} \mu_F^2(i) + \log\left(\frac{\sigma_i^2}{\sigma_0^2}\right)\right) \tag{C12}$$

Note that, for typical prior and posterior variances of DCM connections ($\sigma_i^2 \ll \sigma_0^2 \ll 1$), the (negative) log-ratio term is usually much smaller than the (positive) squared term, rendering the former negligible in practice.

---

[1] Since $\mu_F^T \mu_F = \mu_F^2 = \mu_{F\backslash\{C\}}^T \mu_{F\backslash\{C\}} + \mu_{F\{C\}}^T \mu_{F\{C\}}$ is constant for a given graph, minimizing a function of the removed (between-partition) connections maximizes a corresponding function of the retained (within-partition) connections. Together, these objectives achieve the goal of partitioning or clustering.

#### Appendix D: Linearized DCM

To apply dynamic causal modelling to hundreds or thousands of neuronal states, one can commit to low-order (first-order) approximations (Frässle et al., 2017, 2018; Friston et al., 2021). Consider the state space model describing the coupling among a large number of states $x(t)$, which are then converted to some observable (here, hemodynamic) measurements $y(t)$, through convolution with a (hemodynamic) kernel $k$. The states and observations are subject to system and observation noise, $\omega_x$ and $\omega_y$, respectively. Mathematically, the model is expressed as follows (dropping the time dependence for clarity):

$$\begin{aligned} \dot{x} &= f(x) + \omega_x \\ y &= k * x + \omega_y \end{aligned} \tag{D1}$$

Assuming that an observation is available for each relevant state, and linearizing this state space model, we arrive at the following linear system, where $J = \partial_x f(x)$ and † denotes conjugate transpose (Friston et al., 2021):

$$\left.\begin{matrix} Dx = xJ^\dagger + \omega_x \\ y = Kx + \omega_y \end{matrix}\right\} \Rightarrow \left\{\begin{matrix} KDx = KxJ^\dagger + K\omega_x \\ Dy = KDx + D\omega_y \end{matrix}\right\} \Rightarrow \left\{\begin{matrix} Dy = yJ^\dagger + \omega \\ \omega = K\omega_x + D\omega_y - \omega_y J^\dagger \end{matrix}\right. \tag{D2}$$

Hence, we can approximate the system with a general linear model:

$$\begin{aligned} Dy &= yJ^\dagger + \omega \\ cov(\omega) &= \gamma_1 KK^\dagger + \gamma_2 DD^\dagger + \gamma_3 I \end{aligned} \tag{D3}$$

This approximation assumes that $J^\dagger J \propto I$, since the Jacobian of stable states is dominated by large negative leading diagonals. Equation D3 is a general linear model, with random fluctuations whose covariance components are characterized by the (hemodynamic) convolution kernel and the amplitudes of state and observation noise. Applying variational inference to this linearized system provides approximate Gaussian posteriors for the unknown Jacobian elements (i.e., causal connections) and the unknown covariances of the random effects (Friston et al., 2007).

#### Appendix E: Characterizing the intrinsic dynamics of partitions

The intrinsic dynamics of a partition are characterized by its intrinsic timescales, characteristic frequencies, and the kinetic energy of its eigenmodes. These quantities are defined in terms of the eigenvalues of the partition's Jacobian (i.e. effective connectivity) matrix.

Let the K dominant eigenvalues of the Jacobian matrix $J$ be denoted by $\lambda_1 \geq \lambda_2 \geq \cdots \geq \lambda_K$, ordered according to their (negative) real parts. In linear systems theory, the real part of each eigenvalue determines the exponential rate of decay of perturbations along the corresponding eigenmode. Accordingly, the intrinsic timescales of the dissipative flow associated with the partition are given by the inverse of the negative real parts of the eigenvalues:

$$\tau_i = -1/Re\{\lambda_i\} \tag{E1}$$

Conversely, the imaginary parts of the eigenvalues characterize oscillatory (solenoidal) components of the flow. The intrinsic frequencies associated with these modes are therefore defined as:

$$f_i = Im\{\lambda_i\}/2\pi \tag{E2}$$

The kinetic energy of each dynamical mode can likewise be expressed in terms of the eigenvalues of the Jacobian. Specifically, the kinetic energy associated with eigenmode $i$ is given by:

$$KE_i = \frac{\lambda_i^{\dagger}\lambda_i}{-2Re\{\lambda_i\}} = \frac{Re\{\lambda_i\}^2 + Im\{\lambda_i\}^2}{-2Re\{\lambda_i\}} \tag{E3}$$

Definitions of dissipative and solenoidal flows are provided in Appendix F. In practice, these dynamical quantities are computed by performing an eigen-decomposition of the effective connectivity restricted to the Markov blanket states of a given partition, yielding the eigenvalues required for Eqs. (E1–E3). For further theoretical background and empirical motivation, see (Friston et al., 2021).

#### Appendix F: Computing inverse covariance from effective connectivity

Starting with a Langevin formulation of neuronal dynamics, one can express the flow $f(x)$ at nonequilibrium steady state in terms of a Helmholtz decomposition of the solution to density dynamics—as described by the Fokker Planck equation (Ao, 2004; Benozzo et al., 2024; Friston et al., 2021; Qian & Beard, 2005):

$$\dot{x} = f(x) + \omega$$
$$f(x) = (Q - \Gamma)\,\nabla\Im(x) \qquad \text{(F1)}$$

Here, $\Im(x) = -\ln p\,(x)$ denotes the surprisal or self-information and the antisymmetric (skew) matrix $Q = -Q^{\dagger}$ mediates solenoidal flow. The positive definite matrix $\Gamma \propto I$ is a diffusion tensor describing the amplitude of random fluctuations, $\omega$ (assumed to be a Wiener process). In this Helmholtz decomposition, the flow $f(x)$ can be decomposed into a dissipative (curl-free) gradient flow, $-\Gamma\nabla\Im$, and a solenoidal (divergence-free) flow, $Q\nabla\Im$.

Differentiating the Helmholtz decomposition with respect to the states $x$ yields:

$$\nabla f(x) = (Q - \Gamma)\,\nabla^2\Im(x) = (Q - \Gamma)\,H(x) \Longrightarrow$$
$$J(x) = (Q - \Gamma)\,\Pi(x) \Rightarrow$$
$$\Pi(x) = -(\Gamma - Q)^{-1}J(x) \approx -(\Gamma + Q)\,J(x) \qquad \text{(F2)}$$

Equation F2[2] reveals the relationship between the Jacobian $J(x) = \nabla f(x)$ encoding the causal connections and the conditional independencies specified by a Hessian $H(x) = \nabla^2\Im(x)$, which equals the inverse covariance (i.e. precision) matrix of the neuronal states $\Pi(x) = \Sigma_x^{-1}$ under Gaussian assumptions.

To compute the solenoidal matrix $Q$, note that the symmetry of the Hessian matrix imposes linear constraints on $Q$:

$$(Q - \Gamma)^{-1}\,J(x) = \Pi(x) = \Pi(x)^T = J(x)^T(Q - \Gamma)^{-T} \Longrightarrow$$
$$Q\,J(x)^T + J(x)Q = \Gamma J(x)^T - J(x)\Gamma \qquad \text{(F3)}$$

---

[2] The approximate equality in the final expression arises from a first-order Taylor expansion of the inverse of a mixture of matrices.

Eq. F3 is a Sylvester equation ($AX + XB = C$), which can be solved numerically in MATLAB using the sylvester or lyap functions—based on the Bartels–Stewart algorithm (Bartels & Stewart, 1972)—to derive $Q$ from $J(x)$ and $\Gamma$. Once $Q$ is calculated, it is substituted in Eq. F2, along with $J(x)$ and $\Gamma$, to compute the precision matrix $\Pi(x)$.

In summary, the step-by-step procedure for deriving the inverse covariance from the Jacobian is as follows:

1. Estimate $J(x)$ and $\Gamma$ using DCM
2. Substitute $J(x)$ and $\Gamma$ in Eq. F3, to derive $Q$ (by solving the Sylvester equation)
3. Substitute $Q, J(x)$ and $\Gamma$ in Eq. F2, to derive $\Pi(x)$

A MATLAB implementation of this procedure is available as *spm_ness* function in SPM12. Alternatively, a modified implementation (*spm_ness_mod*) may be accessed from: https://github.com/tszarghami/Parcellation.
